# Munc13-4 mediates tumor immune evasion by regulating the sorting and secretion of PD-L1 via exosomes

**DOI:** 10.1101/2025.03.22.644518

**Authors:** Chuqi Liu, Dexiang Liu, Xiang Zheng, Jiali Guan, Xinyan Zhou, Haikun Zhang, Shen Wang, Qiubai Li, Lu Gan, Jun He, Cong Ma

**Author notes:** Correspondence (Jun He), (Cong Ma). These authors contributed equally (Chuqi Liu, Dexiang Liu, Xiang Zheng).

## Abstract

Tumor-derived extracellular vesicles primarily carry PD-L1 via exosomes, which interact with PD-1 receptors on T cells, impacting immune responses in the tumor microenvironment and beyond, leading to a more extensive immunosuppressive landscape. However, the mechanisms governing exosomal PD-L1 sorting and secretion remain elusive. In this study, we identified Munc13-4 as a crucial regulator of exosomal PD-L1 sorting and secretion. Deletion of Munc13-4 in breast tumors enhances T cell-mediated anti-tumor immunity, suppresses tumor growth, and improves the efficacy of immune checkpoint inhibitors. Our results illustrate how Munc13-4 collaborates with HRS, Rab27, and SNAREs to facilitate PD-L1 sorting and secretion via exosomes. The cryo-EM structure of the Munc13-4–Rab27a complex provide new insights into its role in exosome secretion. Importantly, we discovered that Munc13-4 has a novel role in sorting PD-L1 onto exosomes, which relies on the formation of a ternary complex with PD-L1 and HRS. In addition, IFNγ stimulation modifies Munc13-4 and HRS, establishing a dynamic regulatory mechanism that enables tumor cells to adapt to immune pressure by modulating PD-L1 sorting. Using a specially designed peptide to disrupt the Munc13-4–PD-L1 interaction and impede PD-L1 sorting significantly enhances anti-tumor immunity and slows tumor growth in vivo. These results highlight the potential of targeting the Munc13-4–PD-L1 axis to suppress tumor immune evasion.

## Introduction

Tumor cells evade immune surveillance by increasing the surface expression of programmed death-ligand 1 (PD-L1), which interacts with the programmed death protein 1 (PD-1) receptor on T cells to trigger the immune checkpoint response and leading to T cell inhibition^1,2^. Inhibitors targeting the PD-1/PD-L1 have shown promise in cancer therapy by restoring T cell function and enhancing anti- tumor immunity^3^. Nonetheless, a significant proportion of patients do not respond to anti-PD-L1/PD- 1 therapies^3,4^. A primary reason for this resistance is the secretion of PD-L1 into the bloodstream via extracellular vesicles (EVs), particular exosomes, where it can disrupt immune function remotely^5–7^. Recent studies have demonstrated that genetic blockade of exosomal PD-L1 biogenesis and/or secretion not only suppresses local tumor growth but also elicits a durable systemic anti-tumor immune response^8^. Therefore, the role of exosomal PD-L1 in immune modulation highlights the urgent need for targeted strategies to inhibit exosomal PD-L1 biogenesis and secretion, which could significantly improve the efficacy of cancer immunotherapies.

The secretion of tumor-derived exosomes begins with the formation of early endosomes, which mature into multivesicular bodies (MVBs) containing intraluminal vesicles (ILVs). MVBs fuse with the plasma membrane, releasing ILVs as exosomes into the extracellular space^9^. Endosomal sorting complexes required for transport (ESCRTs) are critical for cargo sorting within the endosomal system. Hepatocyte growth factor-regulated tyrosine kinase substrate (HRS), a key component of the ESCRT machinery, mediates cargo recognition and sorting into MVBs^10^. In addition, the Rab family of small GTPases (Rabs) is essential for membrane trafficking, with Rab27 specifically controlling various steps of the exosome secretion pathway, particularly the docking of MVBs to the plasma membrane^11^. Moreover, the SNARE machinery mediates the fusion of MVBs with the plasma membrane, with syntaxin-4, SNAP-23, and VAMP-7 forming the SNARE complex to drive the secretion of exosomes in various tumor cells^12^. Despite these advances, the mechanisms governing PD-L1 sorting onto exosomes and their secretion remain elusive. Understanding these processes is essential for developing targeted molecules that inhibit the PD-L1 secretion pathway, and it is urgent to identify and investigate key players in the endosomal sorting pathway that mediate PD-L1 recognition and sorting onto exosomes.

In this study, we identified Munc13-4, a member of the Munc13 protein family known for its regulatory role in membrane trafficking, as upregulated in various tumor tissues, where it mediates tumor immune evasion by regulating exosomal sorting and secretion of PD-L1. Deleting Munc13-4 in breast tumors significantly enhances T cell-mediated anti-tumor immunity, suppresses tumor growth, and boosts the efficacy of immune checkpoint inhibitors. We elucidated a coherent mechanism whereby Munc13-4 collaborates with HRS, Rab27, and SNAREs to regulate PD-L1 sorting and secretion via exosomes. Notably, we discovered a novel function of Munc13-4 in PD-L1 sorting that depends on its direct interaction with PD-L1; disrupting this interaction with a specifically designed peptide markedly impaired PD-L1 sorting, leading to enhanced T cell-mediated anti-tumor responses *in vivo*. These findings position Munc13-4 as a promising therapeutic target for boosting immune responses against tumors.

## Results

### Munc13-4 Deficiency in Tumor Cells Inhibits Tumor Growth in An Immunity-Dependent Way

The Munc13 protein family functions as critical regulators of vesicle trafficking and exocytosis across various cell types. While Munc13-1, Munc13-2, and Munc13-3 are involved in the exocytosis of synaptic vesicles and dense-core vesicles in neurons and neuroendocrine cells^13–15^, Munc13-4 has specialized roles in cytotoxic granule exocytosis in immune cells^16–19^ and has recently been implicated in exosome secretion in tumor cells^20^. To explore the role of Munc13-4 in tumor progression, we assessed its expression in tumor and adjacent normal tissues using TIMER2.0 (cistrome.shinyapps.io/timer)^21,22^, a platform that analyzes genomic data from The Cancer Genome Atlas (TCGA). Our analysis revealed significant upregulation of Munc13-4 across various tumor types (**Figure S1A**). Immunohistochemical staining of tumor and adjacent normal tissues on tissue microarrays demonstrated increased Munc13-4 expression in tumors, including breast cancer, thyroid cancer, cholangiocarcinoma, gastrointestinal stromal tumor, pancreatic cancer, and hepatocellular carcinoma (**Figure 1A** and **S1B–S1E**), suggesting a crucial role for Munc13-4 in tumor progression. Given the high global incidence of breast cancer, we focused on investigating the specific role of Munc13-4 in breast cancer.

**Figure 1.**
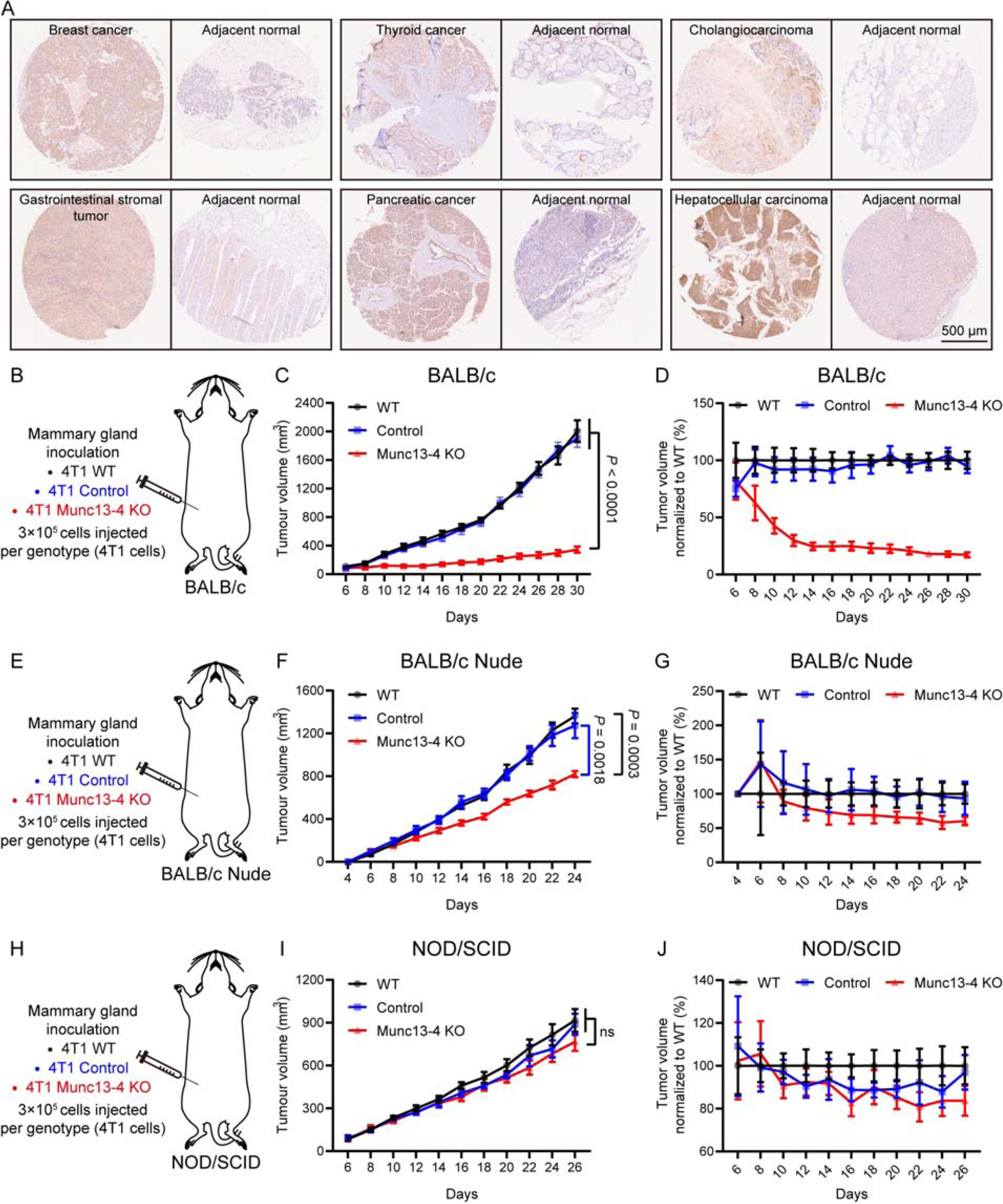
Munc13-4 deficiency in tumor cells inhibits tumor growth in an immunity-dependent way **(A)** Representative immunohistochemical images showing Munc13-4 expression in breast cancer, thyroid cancer, cholangiocarcinoma, gastrointestinal stromal tumors, pancreatic cancer, and hepatocellular carcinoma tissues, along with their corresponding adjacent normal tissues, assessed using a multi-organ carcinoma tissue array. Scale bar, 500 μm. **(B–D)** Tumor growth in BALB/c mice inoculated with wild-type (WT), control or Munc13-4 knockout (KO) 4T1 cells (n = 9). **(B)** Schematic of experimental design. **(C)** Tumor growth curves following mammary gland inoculation. **(D)** Percentage change in tumor volume, normalized to WT group. **(E–G)** Tumor growth in BALB/Nude mice inoculated with WT, control or Munc13-4 KO 4T1 cells (n = 8). **(E)** Schematic of experimental design. **(F)** Tumor growth curves following mammary gland inoculation. **(G)** Percentage change in tumor volume, normalized to WT group. **(H–J)** Tumor growth in NOD/SCID mice inoculated with WT, control or Munc13-4 KO 4T1 cells (n = 6). **(H)** Schematic of experimental design. **(I)** Tumor growth curves following mammary gland inoculation. **(J)** Percentage change in tumor volume, normalized to WT group. Data are presented as means ± SEM, *p*-values were calculated by one-way ANOVA with multiple comparisons (C, F and I), ns, not significant. See also Figure S1 and S2.

Using the CRISPR-Cas9 system, we generated Munc13-4 knockout 4T1 murine mammary carcinoma cells, with control cells infected with a lentivirus carrying Cas9 without sgRNA (**Figure S2A**). The deletion of Munc13-4 did not influence the proliferation of 4T1 cells *in vitro* (**Figure S2B**). We then conducted *in-vivo* studies by creating orthotopic mouse models of breast cancer using wild-type (WT), control and Munc13-4 knockout 4T1 cells (**Figure 1B**). Mice inoculated with Munc13-4 knockout 4T1 cells showed a significant delay in tumor growth compared to those implanted with WT or control cells (**Figure 1C**, **S2C** and **S2D**), with average tumor volume in the knockout group stabilizing at approximately 17.17% of that in the WT group (**Figure 1D**). These data demonstrate that Munc13-4 knockout substantially impairs the oncogenic potential of 4T1 cells, indicating a pivotal role for Munc13-4 in breast tumor progression.

Next, we identified differentially expressed proteins in the proteomes of control and Munc13-4 knockout 4T1 cells, and conducted enrichment analysis based on the KEGG database to pinpoint relevant cellular processes and organismal systems affected by Munc13-4 knockout. KEGG enrichment analysis suggests significant involvement in transport and catabolic pathways, along with a critical association with the immune system (**Figure S2E**). Given that immune evasion is a hallmark of cancer, we explored the relationship between Munc13-4 and immune evasion by assessing the oncogenicity of Munc13-4 knockout 4T1 cells in immunodeficient mouse models. In BALB/Nude mice lacking T cells, the difference in tumor growth between those with Munc13-4 knockout cells and WT or control cells became less pronounced (**Figure 1E, 1F**, **S2F** and **S2G**), with Munc13-4 knockout tumors approaching 60.19% of the size of WT tumors (**Figure 1G**). Moreover, in NOD/SCID mice with severe combined immunodeficiency, tumors in the Munc13-4 knockout group grew comparably to those in the WT group (**Figure 1H, 1I**, **S2H** and **S2I**), with the Munc13-4 knockout tumors reaching 83.67% of the size of the WT tumors (**Figure 1J**). These results suggest that the role of Munc13-4 in breast tumor progression is closely linked to its capacity to modulate immune responses within the tumor microenvironment.

### Munc13-4 Deficiency in Tumor Cells Enhances T Cell Infiltration and Activation

We next explored whether deletion of Munc13-4 in breast tumor cells influences the quantity and activity of T cells within tumors, spleens and lymph nodes of tumor-bearing mice inoculated with either control or Munc13-4 knockout 4T1 cells. Flow cytometry showed a significant increase in the infiltration of both CD4^+^ and CD8^+^ T cells within the tumors of mice implanted with Munc13-4 knockout 4T1 cells (**Figure 2A** and **S3A**), which was corroborated by immunofluorescent staining (**Figure S3B**). In addition, the populations of both CD4^+^ and CD8^+^ cells in the spleens and lymph nodes of these mice were notably elevated compared to the control group (**Figure 2B** and **2C**). These results indicate that the deletion of Munc13-4 in tumor cells enhances T cell infiltration in the tumor, spleen, and lymph nodes.

**Figure 2.**
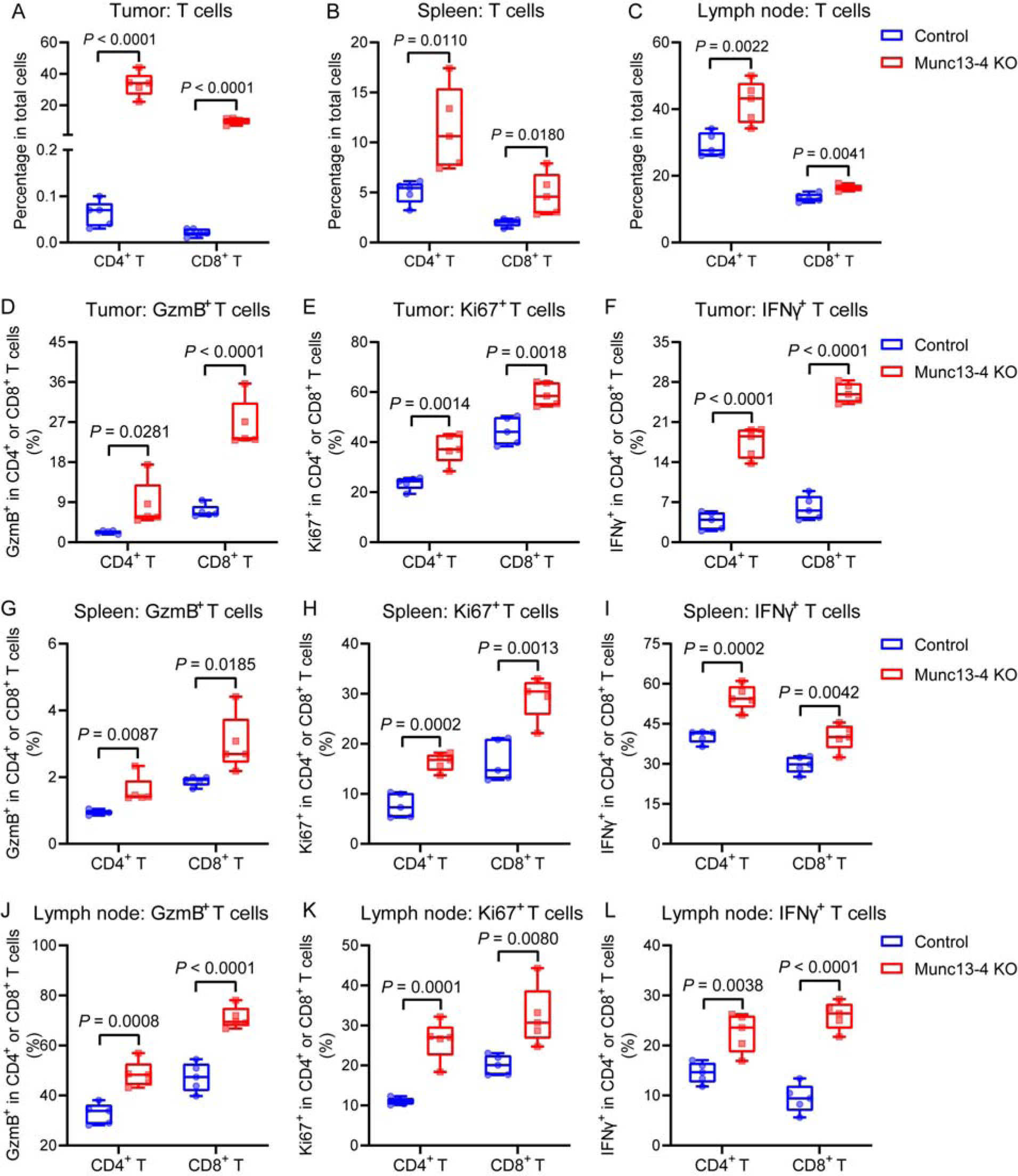
Munc13-4 deficiency in tumor cells enhances T cell infiltration and activation **(A–C)** Flow cytometric quantification of the percentage of CD45^+^CD3^+^CD4^+^ and CD45^+^CD3^+^CD8^+^ T cells among total cells in the tumors **(A)**, spleens **(B)**, and draining lymph nodes **(C)** of BALB/c mice (n = 5), 21 days after mammary gland injection with 3 × 10^5^ control or Munc13-4 KO 4T1 cells per mouse. **(D–F)** Quantification of the percentage of granzyme B^+^ (GzmB^+^) **(D)**, Ki67^+^ **(E)** and IFNγ^+^ **(F)** cells among CD45^+^CD3^+^CD4^+^ and CD45^+^CD3^+^CD8^+^ T cells within tumors from orthotopic mouse models of breast cancer generated by control or Munc13-4 KO 4T1 cells (n = 5). **(G–I)** Quantification of the percentage of granzyme B^+^ **(G)**, Ki67^+^ **(H)** and IFNγ^+^ **(I)** cells among CD45^+^CD3^+^CD4^+^ and CD45^+^CD3^+^CD8^+^ T cells within spleens from orthotopic mouse models of breast cancer generated by control or Munc13-4 KO 4T1 cells (n = 5). **(J–L)** Quantification of the percentage of granzyme B^+^ **(J)**, Ki67^+^ **(K)** and IFNγ^+^ **(L)** cells among CD45^+^CD3^+^CD4^+^ and CD45^+^CD3^+^CD8^+^ T cells within the draining lymph nodes from orthotopic mouse models of breast cancer generated by control or Munc13-4 KO 4T1 cells (n = 5). Box plots show all data points, all *p*-values were calculated by Multiple t tests. See also Figure S3.

We further assessed T cell activation markers, including the cytotoxic molecule granzyme B, the proliferation marker Ki67, and the cytokine interferon gamma (IFNγ), in CD4^+^ and CD8^+^ T cell populations from tumors, spleens, and lymph nodes of tumor-bearing mice. Mice implanted with Munc13-4 knockout 4T1 cells exhibited a significant increase in the expression of granzyme B, Ki67, and IFNγ in both CD4^+^ and CD8^+^ T cells across all examined tissues compared to those inoculated with control 4T1 cells (**Figure 2D–2L** and **S3C–S3E**). Collectively, these results demonstrate that Munc13-4 deficiency in breast tumor cells augments T cell infiltration and activation, thereby enhancing the systemic T cell-mediated immune response.

### Facilitating PD-L1 Secretion by Munc13-4 Suppresses Anti-Tumor Efficacy of T Cells

PD-L1 binding to PD-1 on T cells is a crucial mechanism for tumor evasion of immune surveillance. To explore whether reduced tumor-induced immunosuppression in Munc13-4-deficient models was linked to PD-L1 changes, we examined the effect of Munc13-4 knockout on PD-L1 expression. Our results showed no impact on PD-L1 protein levels in 4T1 and SUM159 breast tumor cells (**Figure 3A**).

**Figure 3.**
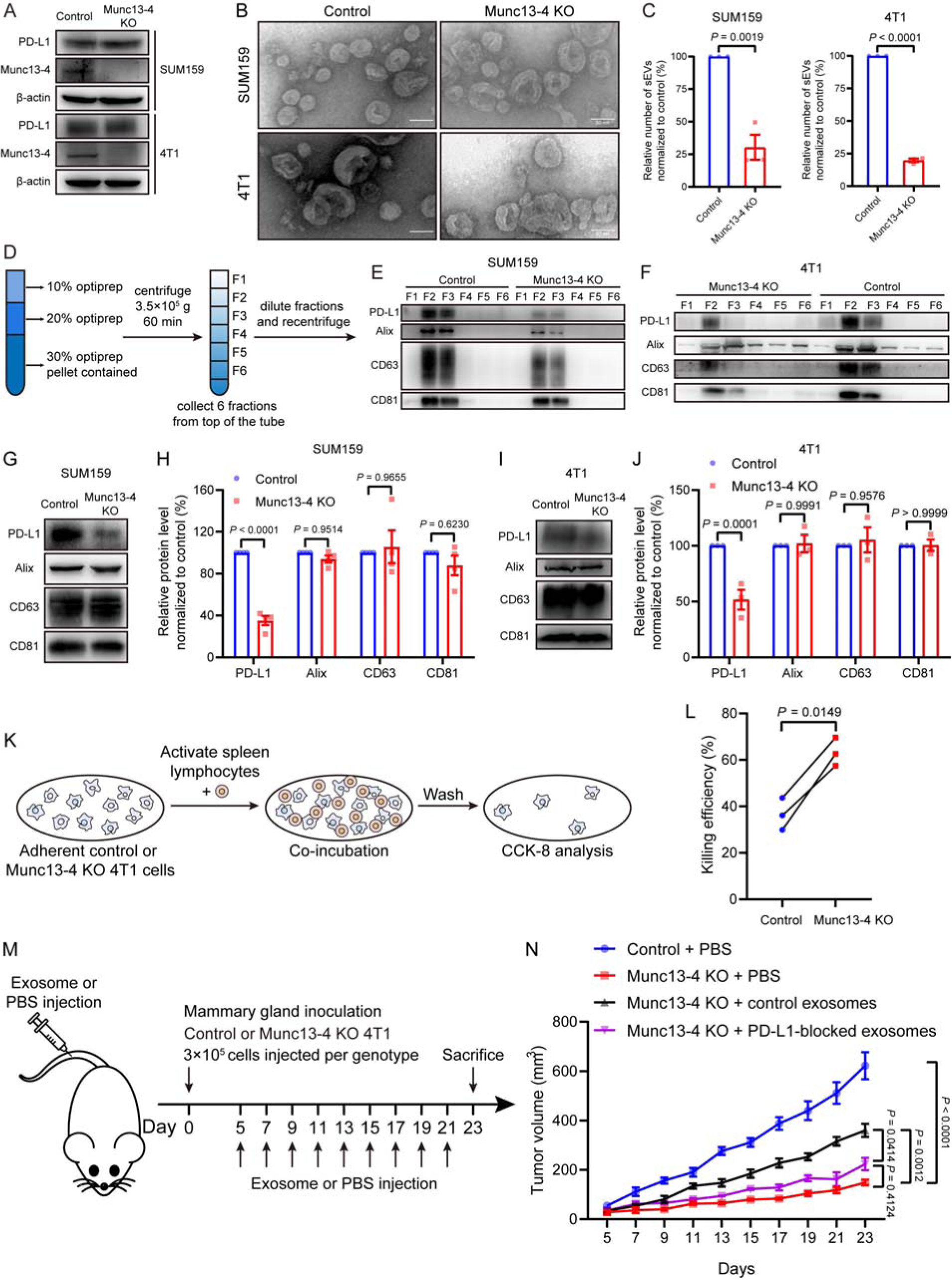
Facilitating PD-L1 secretion by Munc13-4 suppresses anti-Tumor efficacy of T cells **(A)** Western blot analysis of total PD-L1 level in control and Munc13-4 KO SUM159 or 4T1 cells (n = 3). **(B)** Representative TEM images of EVs secreted by control and Munc13-4 KO SUM159 or 4T1 cells. Scale bar, 50 nm. **(C)** Quantification of exosomes secreted by equal numbers of control and Munc13-4 KO SUM159 (left) or 4T1 (right) cells through NTA (n = 3). **(D–F)** Analysis of EVs by optiprep^TM^ density gradient centrifugation. **(D)** Schematic of experimental design. Western blot analysis of PD-L1, Alix, CD63 and CD81 in EVs secreted by equal numbers of control and Munc13-4 KO SUM159 **(E)** or 4T1 **(F)** cell, collected from factions 1–6 (F1–6) (n = 3). **(G and H)** Western blot analysis of PD-L1, Alix, CD63 and CD81 abundance on equal numbers of exosomes secreted by control and Munc13-4 KO SUM159 cells **(G)** and corresponding quantification of blot band intensities **(H)** (n = 3). **(I and J)** Western blot analysis of PD-L1, Alix, CD63 and CD81 abundance on equal numbers of exosomes secreted by control and Munc13-4 KO 4T1 cells **(I)** and corresponding quantification of blot band intensities **(J)** (n = 3). **(K and L)** Assessment of cytotoxicity elicited by activated mouse spleen lymphocytes against control and Munc13-4 KO 4T1 cells. **(K)** Schematic of experimental design. **(L)** Quantification of killing efficiency against control and Munc13-4 KO 4T1 cells (n = 3). **(M and N)** Evaluation of the relationship between decreased oncogenicity and impaired PD-L1 secretion in Munc13-4-deficient 4T1 cells. **(M)** Schematic of experimental design. **(N)** Tumor growth curves following mammary gland inoculation of control or Munc13-4 KO 4T1 cells, with subsequent injection of PBS or the indicated exosomes (n = 6). Data are presented as means ± SEM (C, H, J and N), *p*-values were calculated by unpaired t test (C), two-way ANOVA (H and J), paired t test (L) and one-way ANOVA with multiple comparisons (N). See also Figure S4 and S5.

PD-L1 transport to the plasma membrane and sorting onto exosomes enables its interaction with PD-1 on T cells, thus inhibiting T cell-mediated anti-tumor immunity. Proteomics analysis suggests the involvement of Munc13-4 in transport and catabolic pathways (**Figure S2E**), we thus examined the effect of Munc13-4 knockout on the transport of PD-L1 to the plasma membrane and to exosomes. No change in PD-L1 on the plasma membrane was observed after Munc13-4 knockout (**Figure S4A** and **S4B**). Transmission electron microscopy (TEM) and nanoparticle tracking analysis (NTA) showed that EVs isolated from control and Munc13-4 knockout cell culture supernatants displayed exosome-like morphology (**Figure 3B**) and were within the 50–200 nm size range characteristic of exosomes (**Figure S4C** and **S4D**). However, NTA indicated a significant decrease in the total number of exosomes secreted by Munc13-4 knockout cells compared to control cells (**Figure 3C, S4C** and **S4D**). Western analysis showed a marked reduction in PD-L1 and exosome marker proteins (Alix, CD63, CD81) in EVs from Munc13-4 knockout cells (**Figure S4E** and **S4F**). Isolation of EVs via Optiprep^TM^ density gradient centrifugation confirmed the association of PD-L1 with exosomes and indicate disrupted exosome secretion in Munc13-4 knockout cells (**Figure 3D– 3F**).

To assess PD-L1 levels on exosomes, we collected equivalent numbers of exosomes from both control and knockout cells. Western analysis showed that PD-L1 levels were significantly lower in exosomes from Munc13-4 knockout cells, while levels of Alix, CD63, and CD81 remained unchanged (**Figure 3G–3J**). These results indicate that Munc13-4 deletion does not affect the total PD-L1 levels or its presence on the plasma membrane but significantly reduces secreted PD-L1 by inhibiting exosomes secretion and decreasing its enrichment onto exosomes.

We further assessed the role of exosomes from control and Munc13-4 knockout cells in T cell suppression. Again, equivalent numbers of exosomes from both control and knockout cells were collected. Exosomes from Munc13-4-deficient 4T1 cells showed reduced inhibitory effects on the cytotoxicity of CD8^+^ T cells compared to exosomes from control cells (**Figure S4G** and **S4H**).

Notably, anti-PD-L1 treatment significantly decreased the suppression of CD8^+^ T cell activation by control exosomes, while it had minimal effect on exosomes from Munc13-4 knockout cells (**Figure S4G** and **S4H**). Taken together, the reduced PD-L1 presence on exosomes due to Munc13-4 knockout in tumor cells leads to decreased immunosuppressive capacity.

To directly characterize the anti-tumor efficacy of T cells influenced by Munc13-4 deficiency in tumor cells, we examined the cytotoxicity of T cells primed by either control or Munc13-4-deficient 4T1 cells *in vitro*. Mouse spleen lymphocytes activated with anti-CD3 and anti-CD28 antibodies displayed significantly enhanced cytotoxicity against Munc13-4 knockout 4T1 cells compared to control cells (**Figure 3K** and **3L**). Meanwhile, we explored the relationship between decreased oncogenicity and impaired exosomal PD-L1 secretion in Munc13-4-deficient 4T1 cells *in vivo* (**Figure 3M**). Infusion of exosomes from control 4T1 cells markedly accelerated tumor growth in mice bearing Munc13-4 knockout 4T1 cell transplants (**Figure 3N, S4I** and **S4J**). In contrast, exosomes pre-treated with anti-PD-L1 antibody had minimal impact on tumor growth (**Figure 3N, S4I** and **S4J**). Collectively, these findings underscore the essential role of Munc13-4 in T cell suppression and tumor progression through its regulation of PD-L1 secretion via exosomes.

### Munc13-4 Deficiency in Tumor Cells Boosts Immune Checkpoint Blockade Therapy Effectiveness

Immune checkpoint blockade (ICB) therapy, which targets the PD-1/PD-L1 interaction using antibodies, has become a common approach in cancer treatment. However, it faces challenges such as limited durability of remission and a low overall response rate, restricting its benefits to a small subset of patients^3,4^. Recent studies indicate that PD-L1 on exosomes secreted by tumor cells may antagonize ICB therapy^5–7^. Given our findings that Munc13-4 promotes PD-L1 secretion and inhibits immune surveillance *in vivo*, we investigated whether Munc13-4 deletion in tumor cells could enhance ICB therapy efficacy (**Figure S5A**).

In experiments, neither anti-PD-1 nor anti-PD-L1 treatment slowed tumor growth in mice inoculated with control 4T1 cells compared to IgG isotype controls (**Figure S5B–S5D**). In contrast, tumor growth was significantly delayed in mice implanted with Munc13-4 deficient 4T1 cells, and this effect was further enhanced by anti-PD-1 and anti-PD-L1 treatments (**Figure S5B–S5D**). These results suggest that Munc13-4 depletion in tumor cells improves the therapeutic efficacy of immune checkpoint inhibitors.

### Munc13-4 Does Not Influence MVB Biogenesis

The above findings indicate that Munc13-4 deletion in breast tumor cells has two detrimental effects on PD-L1 secretion: (i) reduced PD-L1 sorting on exosomes and ii) impaired exosome secretion.

This may result from impaired MVB biogenesis and/or MVB fusion with the plasma membrane. To explore the underlying mechanisms, we utilized the human breast tumor cell line SUM159 and examined whether Munc13-4 deficiency disrupts MVB biogenesis. TEM analysis of control and Munc13-4 knockout SUM159 cells revealed a significant accumulation of MVBs in the knockout cells (**Figure S6A** and **S6B**). This was corroborated by immunofluorescence and western analyses showing a marked increase in CD63, a well-established MVB marker (**Figure S6F–S6H**). In addition, TEM analysis indicated an increase in ILVs within MVBs in Munc13-4 knockout cells (**Figure S6C**), suggesting that reduced PD-L1 secretion is not due to impaired MVB biogenesis.

Further TEM analysis showed a significant increase in hybrid structures formed between MVBs and lysosomes in Munc13-4 knockout cells, characterized by electron-dense compartments and double-membrane autophagosomes (**Figure S6A** and **S6D**). Immunofluorescence analysis confirmed increased colocalization of CD63 with the lysosomal marker LAMP1 (**Figure S6I** and **S6J**). Moreover, TEM data indicated that the diameter of MVBs in Munc13-4 knockout cells was significantly enlarged (**Figure S6E**). Together, these observations suggest an increased prevalence of both homotypic fusion among MVBs and heterotypic fusion between MVBs and lysosomes in the absence of Munc13-4.

### Munc13-4 Facilitates MVB Docking and Fusion with The Plasma Membrane

Since Munc13-4 deletion does not affect MVB biogenesis, we investigated its role in downstream processes related to the docking and fusion of MVBs with the plasma membrane. Rab GTPase Rab27a plays a critical role in various stages of the exosome secretion pathway, particularly in the transport and docking of MVBs to the plasma membrane^11^. As an effector of Rab27a, Munc13-4 collaborates with Rab27a to regulate exocytosis in various immune cells^19,23,24^. Consistent with this role, our *in-vitro* binding experiment detected significant binding between Munc13-4 and Rab27a (**Figure S6K**). In contrast, only minimal interactions were detected with Rab5 or Rab7 (**Figure S6K**), which are primarily associated with early and late endosomes, respectively. To study the role the Munc13-4–Rab27a complex in MVB docking, we expressed CD63 tagged with orange fluorescent protein in both control and Munc13-4 (or Rab27a) knockout SUM159 cells. By using total internal reflection fluorescence (TIRF) microscopy, we tracked the movement of MVBs near the plasma membrane^11^. Either Munc13-4 or Rab27a deficiency increased MVB mobility (**Figure S6L–S6O**), indicating the functional significance of Munc13-4 and Rab27a in MVB docking.

To investigate the molecular mechanisms by which Munc13-4 and Rab27a contribute to MVB docking, we determined the cryo-EM structure of the Munc13-4–Rab27a complex (**Figure 4A** and **S7**). Note that the non-hydrolyzable GTP analog GppNHp was added to maintain Rab27a in its active state for efficient binding to Munc13-4 (**Figure S8A–S8D**). The structure was resolved to 3.4 Å in the core region of the complex, while the two C_2_ domains at the N- and C-termini of Munc13-4 exhibited a resolution of 4.4 Å due to their inherent flexibility (**Figure S8E–S8G**). Compared to the solved structure of core domain (C_1_-C_2_B-MUN) of Munc13-1^25^, the overall architecture of Munc13- 4 is more curved (**Figure S8H**). The binding interface of the complex comprises residues F46, W73, and F88 on Rab27a, along with N739, T740, V660, and K661 on Munc13-4 (**Figure 4A**). Mutations at these sites significantly impaired the interaction (**Figure 4B, 4C, S8I** and **S8J**). Consistent with these findings, such mutations resulted in increased MVB mobility (**Figure 4D** and **4E**) and a marked decrease in the total number of exosomes (**Figure 4F**). Hence, the structure information provides new insight into how Munc13-4 works with Rab27a to promote MVB docking.

**Figure 4.**
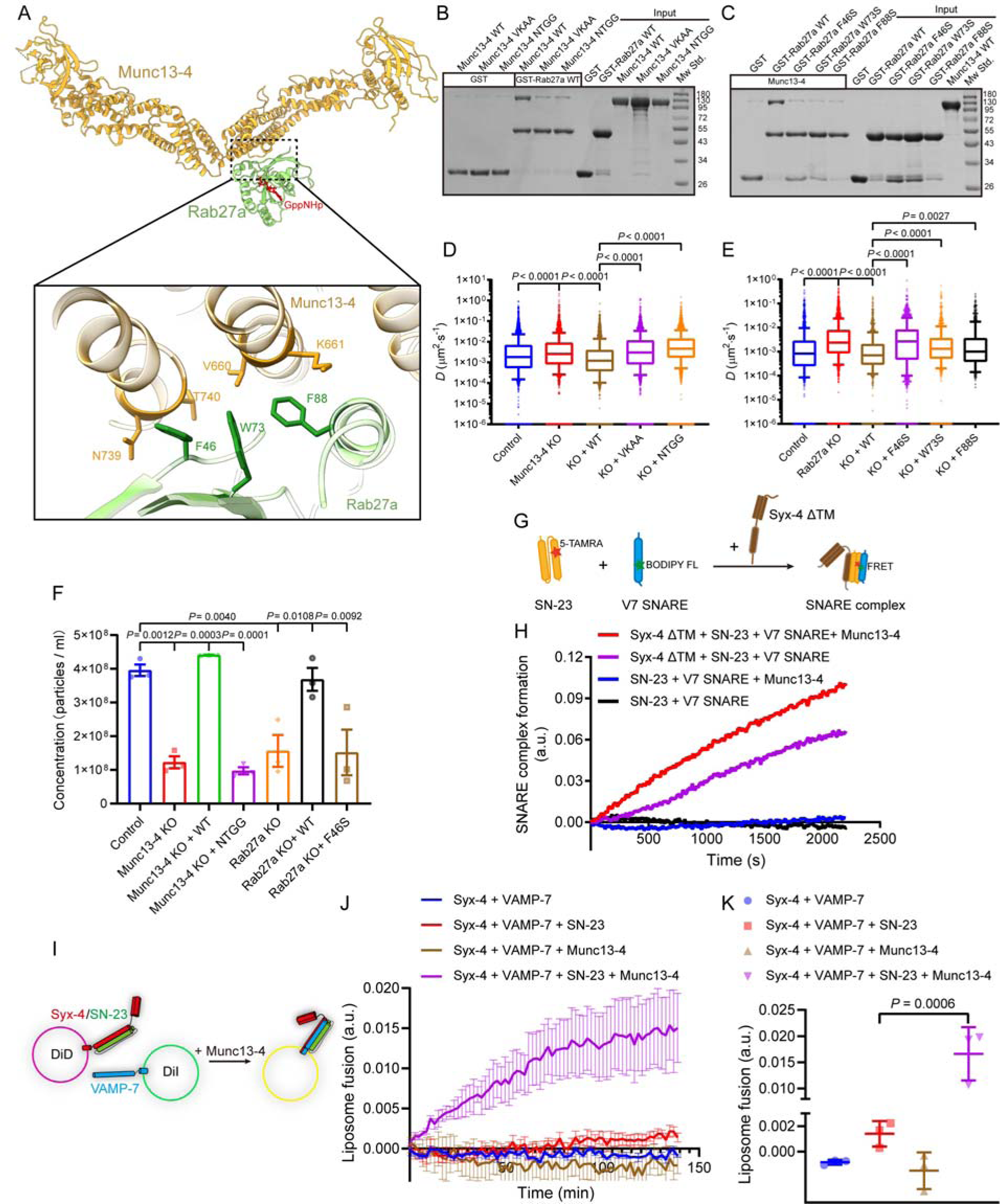
Munc13-4 facilitates MVB docking and fusion with the plasma membrane **(A)** Cryo-EM structure of the Munc13-4–Rab27a complex (upper panel) and detailed interface view (lower panel). Munc13-4 is represented in orange and Rab27a is shown in light green with the GppNHp nucleotide (red) bound. Residues on Rab27a (F46, W73, F88) and Munc13-4 (V660, K661, N739, T740), which are implicated in stabilizing the complex, are highlighted in darker colors. **(B and C)** GST pull-down assays examining the effects of VKAA and NTGG mutations in Munc13- 4 **(B)**, and F46S, W73S, F88S mutations in Rab27a **(C)**, on the formation of the Munc13-4–Rab27a complex (n = 3). **(D and E)** TIRF microscopy analysis of the effects of mutations in Munc13-4 and Rab27a on MVB mobility. Quantification of the mean diffusion coefficient (*D*), an index of MVB mobility, in SUM159 cells with the indicated mutations in Munc13-4 **(D)** or Rab27a **(E)** (n ≥ 962 for each group from triplicate experiments). **(F)** Quantification of exosomes secreted by equal numbers of indicated SUM159 cells though NTA (n = 3). **(G and H)** FRET-based detection of the role of Munc13-4 in SNARE complex assembly. **(G)** Illustration of FRET assay used to detect SNARE complex assembly. VAMP-7 SNARE motif (V7 SNARE) labeled with donor dye BODIPY FL, SNAP-23 (SN-23) labeled with acceptor dye 5- TAMRA, and syntaxin-4 (with its transmembrane domain deleted, termed as Syx4 ΔTM) together form a SNARE complex, leading to FRET between V7 SNARE and SN-23. **(H)** Representative graph of time-dependent SNARE complex assembly measured by the development of FRET between the 5-TAMRA labeled SN-23 and the BODIPY FL labeled V7 SNARE (n = 3). **(I–K)** FRET-based detection of the role of Munc13-4 in liposome fusion. **(I)** Illustration of the liposome fusion experiment. Syntaxin-4 (Syx-4) was incorporated into DiD-labeled liposomes and VAMP-7 was incorporated into DiI-labeled liposomes. Munc13-4 accelerates liposome fusion mediated by SNARE complex, leading to FRET between two liposome populations. **(J)** Time- dependent liposome fusion measured from the development of FRET between the DiD-labeled liposomes and the DiI-labeled liposomes. **(K)** Quantification of the FRET efficiency at the end of the detection (n = 3). Box plots show 10–90% percentile range of all data, with outliers represented as individual dots (D and E), data are represented as means ± SEM (F, J and K), *p*-values were calculated by Kruskal- Wallis test (D and E) and one-way ANOVA with multiple comparisons (F and K). See also Figure S6, S7 and S8.

Recent studies have identified a SNARE complex composed of syntaxin-4, SNAP-23, and VAMP-7 as mediators of MVB fusion with the plasma membrane for exosome release in various tumor cells^12^. Our binding assays demonstrated a direct interaction between Munc13-4 and both SNAP-23 and VAMP-7, and with the assembled SNARE complex (**Figure S6P**). We further assessed the regulatory role of Munc13-4 in SNARE complex assembly and membrane fusion using FRET-based assembly and fusion assays. Our results indicated that Munc13-4 significantly facilitates the assembly of syntaxin-4, SNAP-23, and VAMP-7 into the SNARE complex (**Figure 4G** and **4H**) and promotes the fusion between liposomes bearing syntaxin-4/SNAP-23 and liposomes containing VAMP-7 (**Figure 4I–4K**), underscoring its critical role in the fusion of MVBs with the plasma membrane. Altogether, these results suggest that Munc13-4 works with Rab27a and SNAREs to complete exosome secretion by facilitating the docking and fusion of MVBs with the plasma membrane.

### Exosomal Sorting of PD-L1 by Munc13-4 and HRS

Consistent to our finding that Munc13-4 knockout reduces PD-L1 abundance on exosomes, PD-L1 showed decreased colocalization with CD63 and increased colocalization with LAMP-1 in Munc13- 4 knockout SUM159 cells (**Figure 5A** and **5B**), indicating improper translocation of PD-L1 from MVBs to lysosomes and strengthening the role of Munc13-4 in sorting PD-L1 to exosomes.

**Figure 5.**
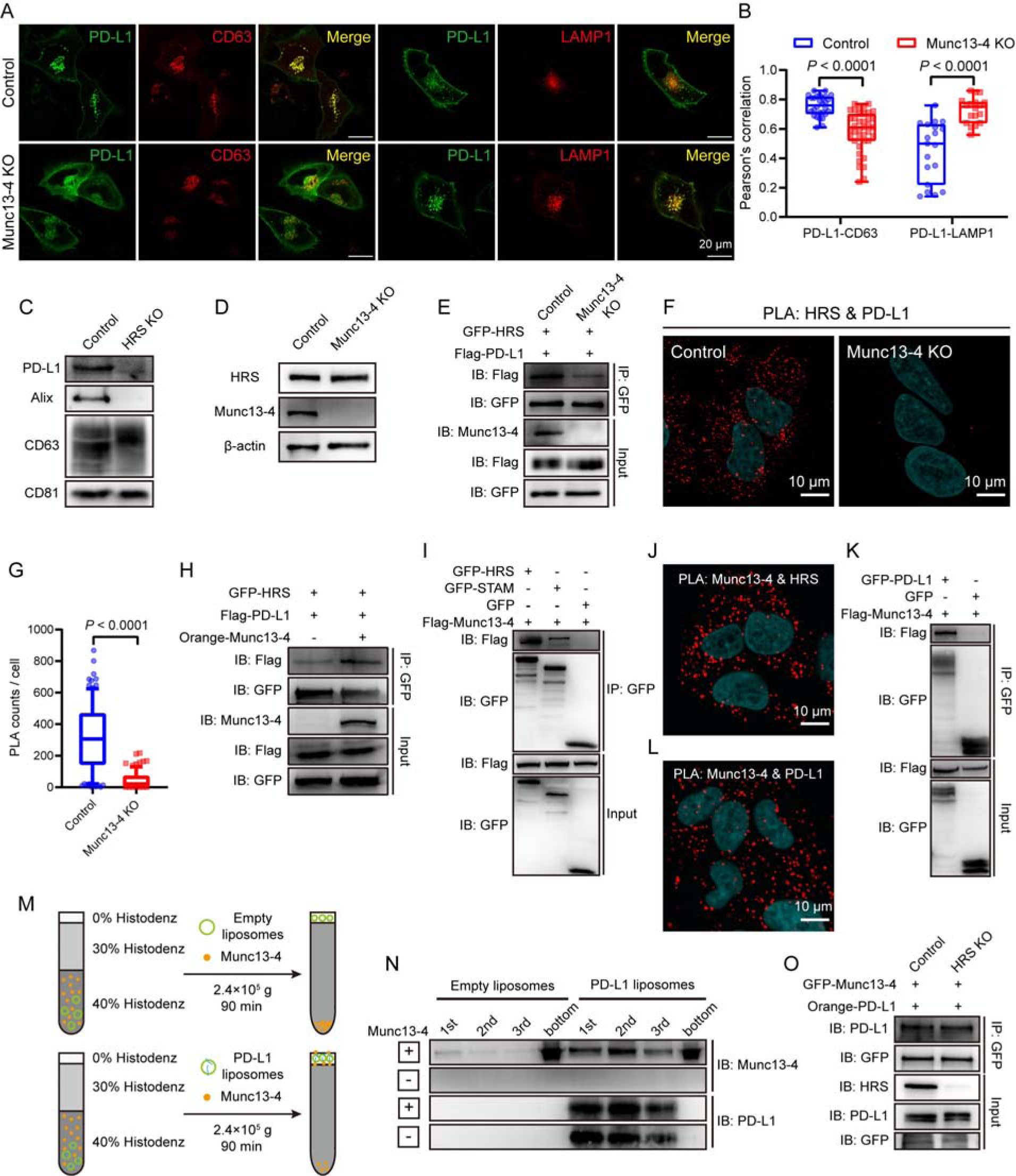
Exosomal sorting of PD-L1 by Munc13-4 and HRS **(A)** Representative confocal microscopy images of control and Munc13-4 KO SUM159 cells co- expressing GFP-tagged PD-L1 with Orange-tagged CD63 or Orange-tagged LAMP1. Scale bar, 20 μm. **(B)** Quantification of Pearson’s correlation coefficient between PD-L1 and CD63, as well as between PD-L1 and LAMP1, in control and Munc13-4 KO SUM159 cells (n = 30 from triplicate experiments). **(C)** Western blot analysis of PD-L1, Alix, CD63 and CD81 abundance on equal numbers of exosomes secreted by control and HRS KO SUM159 cells (n = 2). **(D)** Western blot analysis of total HRS amount in control and Munc13-4 KO SUM159 cells (n = 3). **(E)** Co-IP and immunoblotting (IB) analysis in control and Munc13-4 KO SUM159 cells transfected with indicated constructs to investigate the effect of Munc13-4 knockout on HRS–PD-L1 interaction (n = 3). **(F and G)** PLA to assess the effect of Munc13-4 knockout on the endogenous interaction between HRS and PD-L1. **(F)** Representative confocal microscopy images of control and Munc13-4 KO SUM159 cells in the PLA. Scale bar, 10 μm. **(G)** Quantification of puncta (n = 207 for control group and 175 for Munc13-4 KO group, from triplicate experiments). **(H)** Co-IP and IB analysis in Munc13-4 KO SUM159 cells transfected with indicated constructs to demonstrate that expression of Munc13-4 restored HRS–PD-L1 interaction (n = 3). **(I)** Co-IP and IB analysis in HEK293T cells transfected with indicated constructs to test the interaction of Munc13-4 with HRS and STAM (n = 3). **(J)** Representative confocal microscopy image of SUM159 cells subjected to PLA, detecting the endogenous interaction between Munc13-4 and HRS (n = 3). Scale bar, 10 μm. **(K)** Co-IP and IB analysis in HEK293T cells transfected with indicated constructs to explore the interaction between Munc13-4 and PD-L1 (n = 3). **(L)** Representative confocal microscopy image of SUM159 cells subjected to PLA, showing the endogenous interaction between Munc13-4 and PD-L1 (n = 3). Scale bar, 10 μm. **(M and N)** Direct binding between Munc13-4 and PD-L1 determined by *in vitro* liposome co- flotation assay. **(M)** Schematic of experimental design. **(N)** Western blot analysis of Munc13-4 and PD-L1 in top three fractions and bottom fraction of the mixture. **(O)** Co-IP and IB analysis in control and HRS KO SUM159 cells transfected with indicated constructs to examine the effect of HRS knockout on Munc13-4–PD-L1 interaction (n = 3). Box plots show all data points (B and G), *p*-values were calculated by two-way ANOVA (B) and Mann-Whitney U test (G).

Hepatocyte growth factor-regulated tyrosine kinase substrate (HRS), a key component of ESCRT-0, has been found to mediate PD-L1 sorting^26,27^. This function was verified by the significant reduction in PD-L1 abundance on exosomes following the deletion of HRS (**Figure 5C**). To further explore this, we examined the relationship between HRS and PD-L1 in both the presence and absence of Munc13-4. In Munc13-4 knockout SUM159 cells, the expression of PD-L1 and HRS was unaffected (**Figure 3A** and **5D**); while the interaction between HRS and PD-L1 was markedly diminished compared to control cells, regardless of whether they were exogenously or endogenously expressed (**Figure 5E–5G**). Reintroducing Munc13-4 into knockout cells significantly restored the HRS–PD-L1 interaction (**Figure 5H**), suggesting that this interaction relies on Munc13-4.

Supporting these findings, co-immunoprecipitation (co-IP) assay showed a marked binding preference of Munc13-4 for HRS over STAM (another ESCRT-0 component) under exogenous expression conditions (**Figure 5I**). In addition, proximity ligation assay (PLA) indicated an interaction between endogenous expressed Munc13-4 and HRS in SUM159 cells (**Figure 5J**). We also examined whether Munc13-4 directly interacts with PD-L1. Both co-IP and PLA assays revealed this interaction in SUM159 cells (**Figure 5K** and **5L**). This interaction was further verified through *in-vitro* liposome co-flotation assay (**Figure 5M** and **5N**). Importantly, the deletion of HRS in SUM159 cells did not influence the interaction between Munc13-4 and PD-L1 (**Figure 5O**), indicating that the Munc13-4–PD-L1 interaction is independent of HRS. Altogether, these results suggest the formation of a ternary complex comprising HRS, Munc13-4, and PD-L1, which is critical for sorting of PD-L1 to exosomes.

### IFNγ-induced Modifications of Munc13-4 and HRS Exert Opposing Effects on PD-L1 Sorting

IFNγ, a cytokine produced by NK and T cells, contributes substantially to immunosurveillance against tumors by activating immune cells and inducing apoptosis in tumor cells^28,29^. Conversely, tumor cells can exploit IFNγ signaling to evade immune destruction through the elevation of PD-L1 expression, which inhibits immune cell activity^28,29^. Indeed, IFNγ stimulation dramatically increased the protein level of PD-L1 (**Figure S9A** and **S9B**), leading to an elevated presence of PD-L1 on both the plasma membrane (**Figure S9C** and **S9D**) and exosomes (**Figure S9E** and **S9F**), without altering the total quantity of secreted exosomes (**Figure S9E–S9G**). Considering the significant effect of exosomal PD-L1 on the suppression of T cell activity, we hence explored whether and how HRS and Munc13-4 regulate PD-L1 sorting onto exosomes in response to IFNγ.

Unlike PD-L1, the overall levels of HRS and Munc13-4 remained unchanged under IFNγ treatment in SUM159 cells (**Figure S9A** and **S9B**). Strikingly, IFNγ stimulation significantly decreased the ubiquitylation of HRS (**Figure S9H**), without affecting its acetylation and phosphorylation (**Figure S9I–S9K**). Meanwhile, IFNγ stimulation significantly increased the acetylation of Munc13-4 (**Figure S9L**), without influencing its phosphorylation and ubiquitylation (**Figure S9M–S9O**). These results indicate that both HRS ubiquitylation and Munc13-4 acetylation induced by IFNγ may cooperate to regulate PD-L1 sorting onto exosomes.

To explore the mechanism that regulates Munc13-4 acetylation, we individually expressed six common acetyltransferases—GCN5, PCAF, CBP, P300, TIP60, and HBO1—with Munc13-4 in HEK293T cells and found that CBP and P300 are able to acetylate Munc13-4 (**Figure S10A**). Knockout of CBP—rather than P300—reduced the acetylation of endogenous Munc13-4 in SUM159 cells, both in the absence and presence of IFNγ (**Figure S10B**). In addition, IFNγ stimulation promoted the translocation of CBP from the nucleus to the cytoplasm (**Figure S10C** and **S10D**).

These data indicate that CBP acts as a physiological acetyltransferase for Munc13-4. Then, we screened for the deacetylase mediating Munc13-4 deacetylation using deacetylase inhibitors. We found that trichostatin A (TSA), an inhibitor of histone deacetylases (HDACs)^30^, increased Munc13-4 acetylation, while nicotinamide (NIC), an inhibitor of class-III sirtuin deacetylases (SIRTs)^30^, had no significant effect (**Figure S10E**). Among the HDAC1–8 isoforms, expression of HDAC3 and HDAC4, but not the other HDACs, reduced Munc13-4 acetylation in HEK293T cells (**Figure S10F– S10H**). Notably, knockout of HDAC3, rather than HDAC4, increased the acetylation level of endogenous Munc13-4, regardless of IFNγ treatment (**Figure S10I**), suggesting that HDAC3 serves as a physiological deacetylase for Munc13-4. It is noteworthy that the deletion of CBP and HDAC also influenced Munc13-4 expression, as evidenced by a reduction in Munc13-4 transcription (**Figure S10J** and **S10L**) and a corresponding decrease in the total amount of Munc13-4 (**Figure S10K** and **S10M**). We clarify that to ensure the accuracy of the data, our evaluation of the acetylation level of Munc13-4 in the experiments mentioned above was performed with the condition that the total amounts of Munc13-4 samples remained consistent (See **Methods**) We proceeded to investigate the impact of Munc13-4 acetylation on the sorting of PD-L1 onto exosomes. In the presence of IFNγ, the knockout of CBP and HDAC3 did not influence the total amount of PD-L1 (**Figure S10N** and **S10O**), but displayed opposing effects on PD-L1 sorting.

Specifically, the knockout of CBP, which reduces Munc13-4 acetylation, led to a significant increase in the abundance of PD-L1 on exosomes (**Figure 6A** and **6B**). In contrast, the knockout of HDAC3, which enhances Munc13-4 acetylation, resulted in a remarkable decrease in PD-L1 abundance on exosomes (**Figure 6C** and **6D**). Hence, these results consistently suggest that Munc13-4 acetylation inhibits the sorting of PD-L1 onto exosomes.

**Figure 6.**
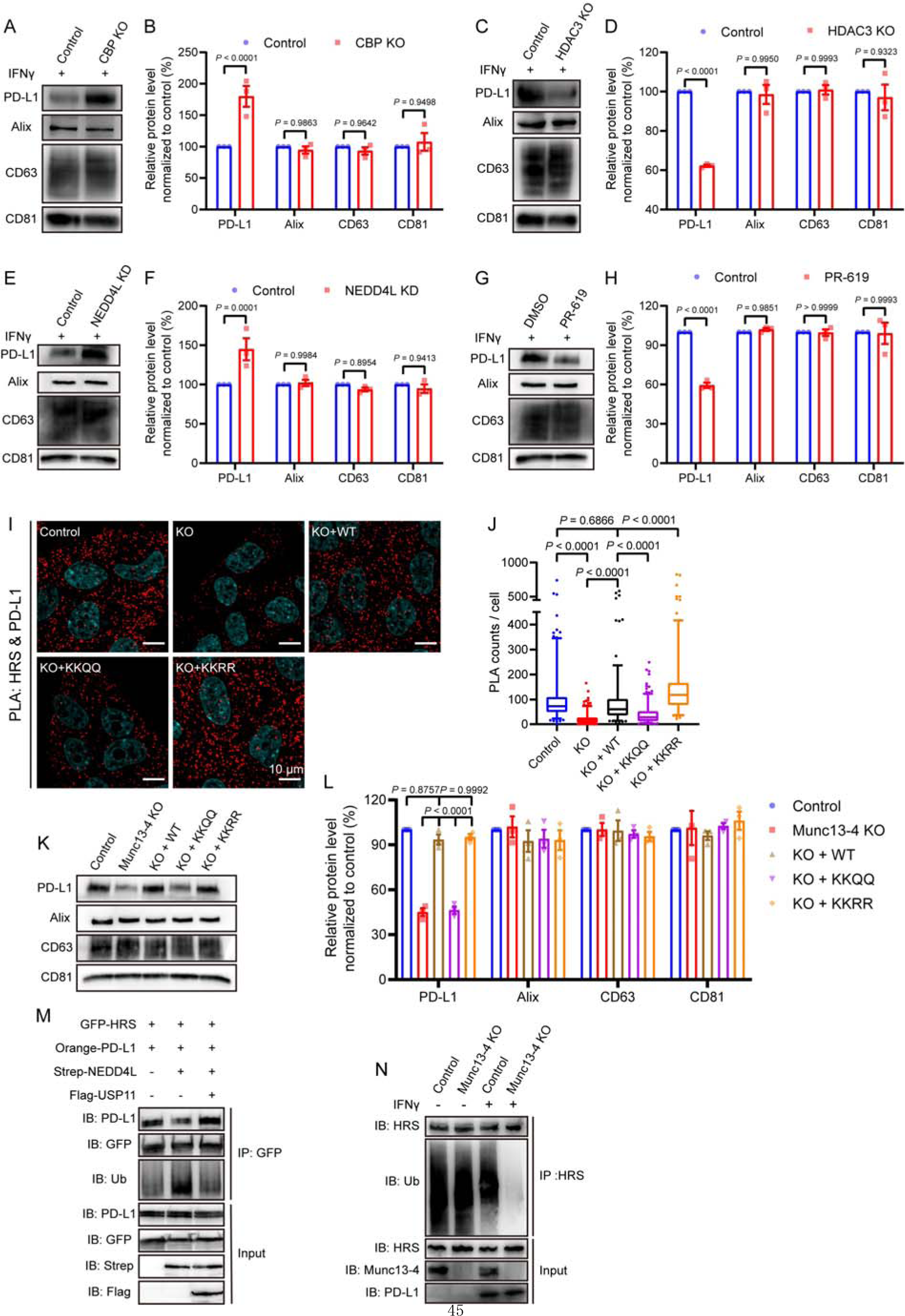
IFN**γ**-induced modifications of Munc13-4 and HRS exert opposing effects on PD-L1 sorting **(A and B)** Western blot analysis of PD-L1, Alix, CD63 and CD81 abundance on equal numbers of exosomes secreted by control and CBP KO SUM159 cells under IFNγ treatment **(A)** and corresponding quantification of blot band intensities **(B)** (n = 3). **(C and D)** Western blot analysis of PD-L1, Alix, CD63 and CD81 abundance on equal numbers of exosomes secreted by control and HDAC3 KO SUM159 cells under IFNγ treatment **(C)** and corresponding quantification of blot band intensities **(D)** (n = 3). **(E and F)** Western blot analysis of PD-L1, Alix, CD63 and CD81 abundance on equal numbers of exosomes secreted by control and NEDD4L knockdown (KD) SUM159 cells under IFNγ treatment **(E)** and corresponding quantification of blot band intensities **(F)** (n = 3). **(G and H)** Western blot analysis of PD-L1, Alix, CD63 and CD81 abundance on equal numbers of exosomes secreted by SUM159 cells co-treated with IFNγ and PR-619 or DMSO **(G)** and corresponding quantification of blot band intensities **(H)** (n = 3). **(I and J)** PLA to assess the effects of Munc13-4 mutations on the interaction between endogenous HRS and PD-L1 in SUM159 cells. Representative confocal images of indicated SUM159 cells in the PLA **(I)** and quantification of PLA puncta **(J)** (n = 181 for control group, 252 for KO group, 174 for KO + WT group, 240 for KO + KKQQ group and 185 for KO + KKRR group from triplicate experiments). Scale bar, 10 μm. **(K and L)** Western blot analysis of PD-L1, Alix, CD63 and CD81 abundance on equal numbers of exosomes secreted by indicated SUM159 cells **(K)** and corresponding quantification of blot band intensities **(L)** (n = 3). **(M)** Co-IP and IB analysis in HEK293T cells transfected with indicated constructs to investigate the effect of HRS ubiquitylation on its interaction with PD-L1 (n = 3). **(N)** IP and IB analysis in control and Munc13-4 KO SUM159 cells, with or without IFNγ treatment, to explore the ubiquitylation of HRS (n = 3). Data are represented as means ± SEM (B, D, F, H and L), box plot shows 5–95% percentile range of all data, with outliers represented as individual dots (J), *p*-values were calculated by two-way ANOVA (B, D, F, H and L) and Kruskal-Wallis test (J). See also Figure S9–S11.

On the other hand, IFNγ stimulation induces HRS deubiquitylation, and we next investigate the effect of HRS ubiquitylation on PD-L1 sorting. To identify the E3 ligases responsible for HRS ubiquitylation, we utilized the UbiBrowser 2.0 database (http://ubibrowser.bio-it.cn/ubibrowser_v3/)^31^. Among the identified and predicted E3 ligases, including NEDD4^32^, NEDD4L^33^, SH3RF1^34^, ITCH, CBL and PRKN, we found that NEDD4L efficiently catalyzed HRS ubiquitylation in HEK293T cells (**Figure S10P**). In SUM159 cells, NEDD4L knockdown led to a significant reduction in HRS ubiquitylation, regardless of IFNγ stimulation (**Figure S10Q**). Notably, under IFNγ treatment, NEDD4L knockdown resulted in a marked increase in the amount of PD-L1 on exosomes (**Figure 6E** and **6F**) without altering total amount of PD-L1 (**Figure S10Q**).

Furthermore, expression of several common deubiquitinases, including STAMBPL1, CYLD, USP11, USP7, and USP8, was found to reduce NEDD4L-mediated HRS ubiquitylation in HEK293T cells (**Figure S10R**), suggesting that the regulation of HRS deubiquitylation involves multiple deubiquitinases. Treatment with PR-619, a broad-spectrum reversible inhibitor of ubiquitin isopeptidases^35,36^, induced a substantial increase in HRS ubiquitylation in SUM159 cells independent of IFNγ (**Figure S10S**). However, PR-619 treatment resulted in a significant reduction in the abundance of PD-L1 on exosomes under IFNγ stimulation (**Figure 6G** and **6H**), without influencing the overall PD-L1 level (**Figure S10S**). Collectively, these findings demonstrate that HRS deubiquitylation facilitates the sorting of PD-L1 onto exosomes.

### IFN**γ**-induced Modifications of Munc13-4 and HRS Regulate PD-L1 Binding

The finding that Munc13-4 acetylation inhibits PD-L1 sorting suggests that it impairs the interaction between Munc13-4 and PD-L1. Indeed, the enhanced acetylation of Munc13-4, driven by the expression of CBP, led to a reduction in its interaction with PD-L1 (**Figure S11A**), as well as a weakened association of PD-L1 with HRS (**Figure S11B**), while not affecting the interaction between HRS and Munc13-4 (**Figure S11B**). This suggests that Munc13-4 serves as a central hub for the formation of the HRS–Munc13-4–PD-L1 complex. Interestingl<u>y</u>, the mutant deleting the C-terminus of Munc13-4 (termed Munc13-ΔC, lacking residues1049–1090) significantly diminished the acetylation by CBP (**Figure S11C** and **S11D**), indicating that the acetylation sites are located within this region. Furthermore, Munc13-ΔC failed to bind PD-L1 (**Figure S11E**), suggesting that the residues within 1049–1090 are crucial for both acetylation and PD-L1 binding. However, Munc13-ΔC retained its ability to bind HRS. Further screening revealed that residues 546–782 within the MUN domain of Munc13-4 mediate HRS interaction (**Figure S11F**).

We aimed to identify the acetylation sites on Munc13-4. The region between residues 1049 and 1090 contains two lysine (K) residues, and mass spectrometry analysis revealed that both K1062 and K1079 were acetylated in the presence of CBP (**Figure S11G** and **S11H**). Single mutations of either K1062 or K1079 to arginine (K1062R or K1079R) preserved the positive charge but impaired acetylation. Moreover, the double mutant (K1062R/K1079R, termed KKRR) nearly completely abolished CBP-mediated acetylation of Munc13-4 (**Figure S11I**), confirming our mass spectrometry findings. Detected by co-IP, mutating either K1062 or K1079 in Munc13-4 to glutamine (Q), which mimics acetylation, did not affect its interaction with PD-L1 (**Figure S11J**). However, double mutations (K1062Q/K1079Q, termed KKQQ) significantly impaired this interaction (**Figure S11J** and **S11K**). In contrast, the KKRR mutant, which abolishes acetylation, did not affect the Munc13-4– PD-L1 interaction (**Figure S11K**).

In Munc13-4 knockout SUM159 cells, where the endogenous interaction between HRS and PD- L1 was severely damaged, expression of Munc13-4 WT rescued their endogenous interaction to the level comparable to that in control cells (**Figure 6I** and **6J**). In contrast, Munc13-4 KKQQ only slightly restored the HRS–PD-L1 interaction (**Figure 6I** and **6J**), while Munc13-4 KKRR substantially rescued this interaction (**Figure 6I** and **6J**). Similar results were observed in HEK293T cells (**Figure S11L**). Furthermore, the decrease in the PD-L1 abundance on exosomes observed upon Munc13-4 deletion was fully restored by the expression of either Munc13-4 WT or the KKRR mutant, but not by the KKQQ mutant (**Figure 6K** and **6L**). Overall, CBP-mediated acetylation of Munc13-4 at K1062 and K1079 disrupts its interaction with PD-L1, which in turn impairs the HRS– PD-L1 interaction and hinders the sorting of PD-L1 onto exosomes.

We next investigated the mechanism by which HRS deubiquitylation promotes PD-L1 sorting onto exosomes. Expression of NEDD4L in HEK293T cells, which significantly enhanced HRS ubiquitylation, notably suppressed the interaction between HRS and PD-L1 (**Figure 6M**). In contrast, additional expression of USP11, which decreased HRS ubiquitylation, restored the HRS–PD-L1 interaction (**Figure 6M**). These findings suggest that HRS ubiquitylation negatively regulates its interaction with PD-L1. In SUM159 cells, deletion of Munc13-4 also led to a reduction in HRS ubiquitylation, independent of IFNγ stimulation (**Figure 6N**), which likely compensates for the diminished HRS–PD-L1 interaction caused by Munc13-4 deficiency, suggesting a potential compensatory mechanism in PD-L1 sorting in the absence of Munc13-4.

### A Peptide that Disrupts PD-L1**–**Munc13-4 Interaction Inhibits Tumor Growth

Given the critical role of Munc13-4 in sorting PD-L1 to exosomes, we sought to disrupt their interaction in tumor cells to mitigate immune evasion. Targeting the PD-L1–Munc13-4 interaction might be a promising therapeutic strategy. Our co-IP analysis identified both the cytoplasmic motif and the transmembrane domain of PD-L1 as essential for binding to Munc13-4 (**Figure S12A**).

Using AlphaFold, we modeled five 3D structures of the PD-L1 (253–290)–Munc13-4 (1049–1080) complex and determined that the PD-L1 region (256–273) is crucial for this interaction (**Figure 7A** and **S12B**).

**Figure 7.**
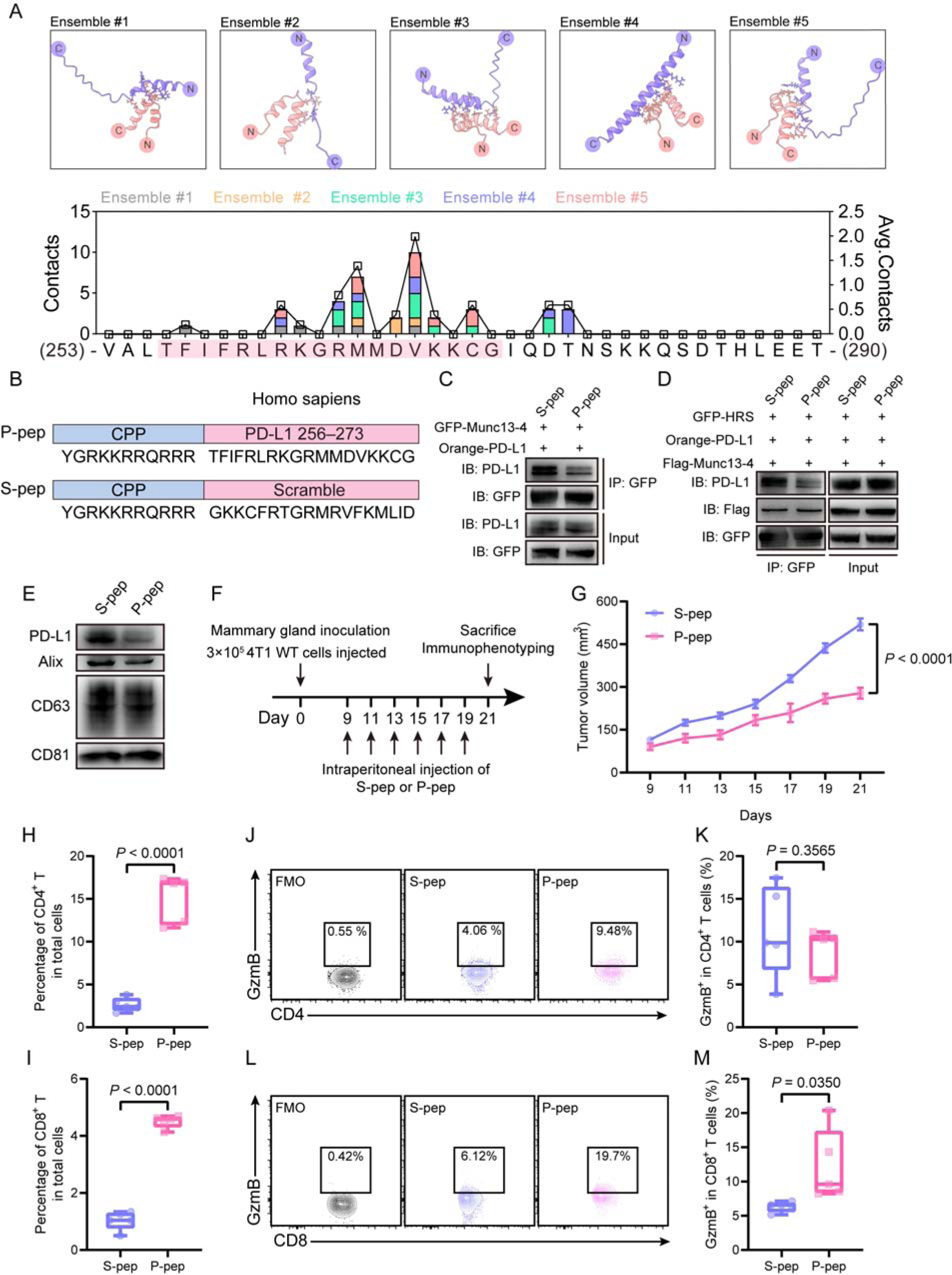
A peptide disrupting PD-L1**–**Munc13-4 interaction inhibits tumor growth **(A)** Prediction of the interaction between PD-L1 (residues 253–290) and Munc13-4 (residues 1049–1080). Top insets show five ensembles generated by AlphaFold multimer, where N- and C-terminus of the proteins are indicated. PD-L1 and Munc13-4 are colored in blue and red, respectively. Residues that show potential contacts with each other (within 4 Å) are shown in sticks. Bottom inset displays the statistics of the residues in PD-L1 that show contacts with Munc13-4. The absolute and averaged contact numbers for each ensemble are illustrated by stacked histograms and line-scatter plot. The potential Munc13-4–interacting sequence of PD-L1 is shaded in magenta. **(B)** Diagram of the sequences for P-pep and S-pep. P-pep comprises a cell-penetrating peptide (CPP) fused to the human PD-L1 256–273 motif, whereas S-pep consists of a CPP linked to a scrambled sequence containing the same amino acid composition as the human PD-L1 256–273 motif. **(C)** Co-IP and IB analysis in HEK293T cells transfected with indicated constructs and incubated with P-pep or S-pep to examine the effect of P-pep on Munc13-4–PD-L1 interaction (n = 3). **(D)** Co-IP and IB analysis in HEK293T cells transfected with indicated constructs and incubated with P-pep or S-pep to assess the effect of P-pep on the interactions of HRS with PD-L1 and Munc13-4 (n = 3). **(E)** Western blot analysis of PD-L1, Alix, CD63 and CD81 abundance on the same number of exosomes secreted from equal number of SUM159 cells treated with S-pep or P-pep. **(F and G)** Assessment of *in vivo* anti-tumor efficacy of P-pep. **(F)** Schematic of experimental design. **(G)** Tumor growth curves of orthotopic mouse models of breast cancer treated with P-pep or S-pep (n = 9). **(H and I)** Flow cytometric quantification of the percentage of CD45^+^CD3^+^CD4^+^ **(H)** and CD45^+^CD3^+^CD8^+^ **(I)** T cells among total cells in tumors (n = 5). **(J and L)** Representative contour plots depicting CD45^+^CD3^+^CD4^+^ **(J)** and CD45^+^CD3^+^CD8^+^ **(L)** T cell populations within tumors, showing the expression of granzyme B. **(K and M)** Quantification of the percentage of granzyme B^+^ cells among CD45^+^CD3^+^CD4^+^ **(K)** and CD45^+^CD3^+^CD8^+^ **(M)** T cells within tumors (n = 5). Data are represented as means ± SEM (G), box plots show all data points (H, I, K and M), *p*-values were all calculated by unpaired t test. See also Figure S12.

Next, we tested whether the 256–273 sequence could competitively inhibit the interaction between PD-L1 and Munc13-4. Overexpressing this sequence in HEK293T cells significantly disrupted their interaction (**Figure S12C**). This disruption also affected the interaction between HRS and PD-L1, but did not impact the HRS–Munc13-4 interaction (**Figure S12D**). Further exploration utilized cell-penetrating peptide (CPP)^37,38^ to deliver the PD-L1 256–273 peptide (P-pep) and a scrambled version (S-pep) (**Figure 7B**). Treatment of HEK293T cells with P-pep, compared to S-pep, significantly inhibited the ectopic interaction between PD-L1 and Munc13-4 (**Figure 7C**) and consequently disrupted the interaction between HRS and PD-L1 (F**igure 7D**), while the HRS– Munc13-4 interaction remained unaffected (F**igure 7D**). In addition, P-pep treatment of SUM159 cells led to a marked decrease in PD-L1 levels on exosomes (**Figure 7E**), suggesting that competitive inhibition of PD-L1 sorting is a viable approach.

We also evaluated the in vivo effects of P-pep on tumor growth using orthotopic breast cancer mouse models (**Figure 7F** and **S12E**). Mice treated with P-pep exhibited a significant delay in tumor growth compared to those receiving S-pep (**Figure 7G, S12F** and **S12G**). Immunophenotyping revealed increased infiltration of both CD4^+^ and CD8^+^ T cells in the tumors of P-pep-treated mice (**Figure 7H** and **7I**), along with enhanced cytotoxicity of tumor-infiltrating CD8^+^ T cells (**Figure 7L** and **7M**), while CD4^+^ T cell cytotoxicity remained unchanged (**Figure 7J** and **7K**). Comprehensive assessments, including body weight, blood routine tests, biochemical analysis, and histological evaluations of major organs (**Figure S12H–S12Q**), indicated no noticeable toxicity from either P- pep or S-pep. Collectively, these results demonstrate that P-pep effectively targets the PD-L1– Munc13-4 interaction, reducing tumor-induced immunosuppression and inhibiting tumor growth without systemic side effects.

## Discussion

Tumor-derived exosomes carry PD-L1, which engages with PD-1 receptors on T cells, influencing immune responses within the tumor microenvironment and in distant sites, resulting in a more extensive immunosuppressive environment^5,8,39^. Therapeutic strategies aimed at counteracting the immunosuppressive effects of exosomal PD-L1 require the development of molecules that can effectively suppress its extracellular secretion. Hence, it is essential to identify the regulatory factors governing the secretory pathway of exosomal PD-L1 and to elucidate the underlying mechanisms. In this study, we reveal a critical role for Munc13-4 in modulating the immunosuppressive effects of exosomal PD-L1 by influencing its sorting and secretion. Specifically, deleting Munc13-4 in breast tumor cells significantly reduces both the number of secreted exosomes and the abundance of PD-L1 on these exosomes, which systematically enhances T cell-mediated anti-tumor responses, suppresses tumor progression and improves the efficacy of immune checkpoint inhibitors. The underlying mechanisms involve i) formation of the Munc13-4–PD-L1–HRS ternary complex, which promotes efficient sorting of PD-L1 to MVBs and loading onto exosomes; ii) assembly of the Munc13-4–Rab27a complex, which enables proper MVB docking; and iii) cooperation between Munc13-4 and the SNARE complex comprising syntaxin-4, SNAP-23 and VAMP-7, which facilitates MVB fusion with the plasma membrane to release PD-L1-containing exosomes. Notably, employing a specially designed peptide to disrupt the Munc13-4–PD-L1 interaction, thereby impairing PD-L1 sorting, significantly enhances anti-tumor immunity and slows tumor growth *in vivo* (**Figure 8**). This underscores the potential of targeting the Munc13-4–PD-L1 axis as a novel approach to augment the efficacy of immune checkpoint inhibitors.

**Figure 8.**
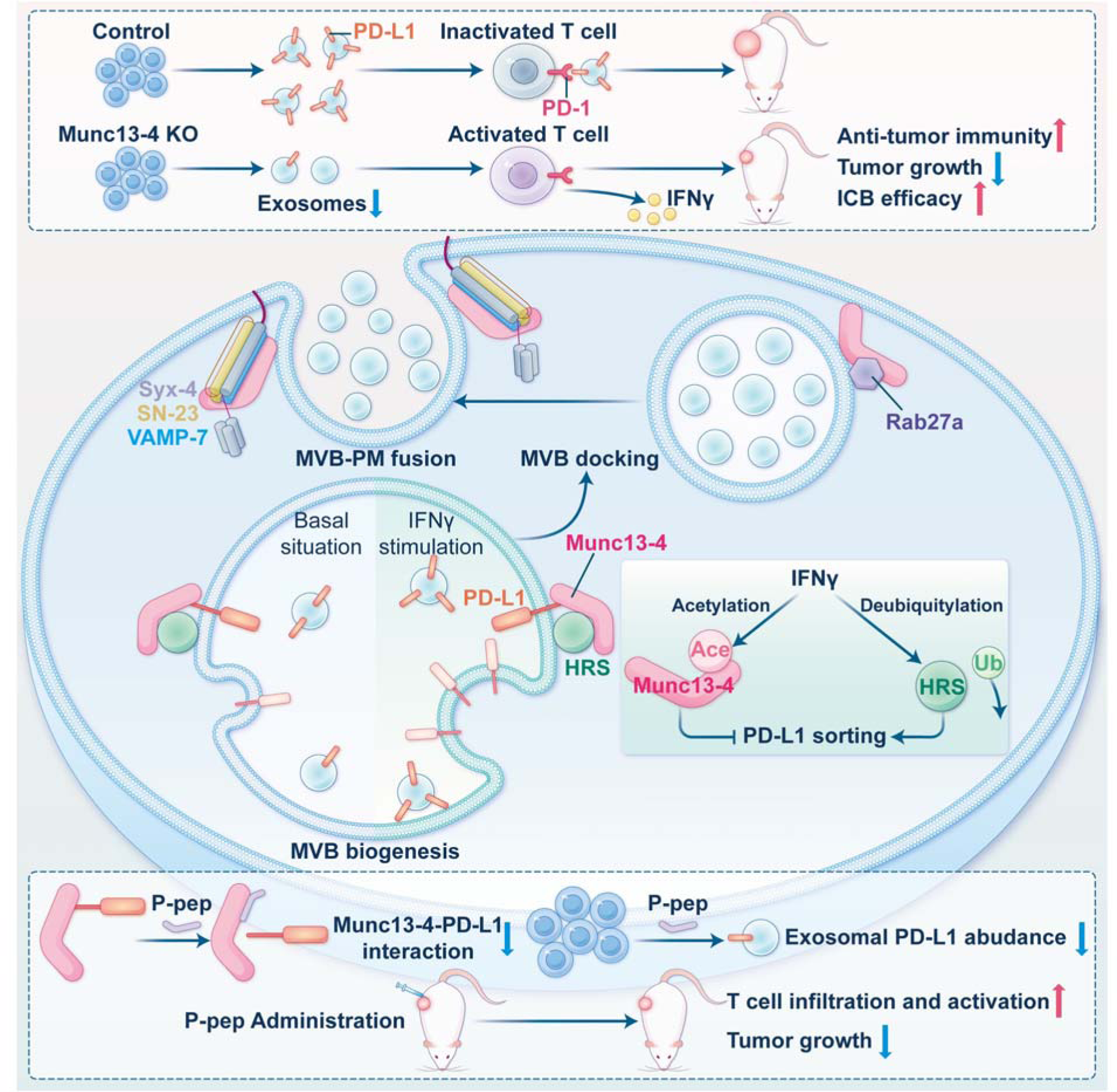
**Mechanistic model of Munc13-4-mediated tumor immune evasion through the regulation of PD-L1 sorting and secretion via exosomes**

Munc13-4 is ubiquitously expressed in various types of cells and contains a MUN domain flanked by two C_2_ domains^40^. Munc13-4 has attracted significant attention for its role in the exocytosis of secretory granules in immune cells. Inherited variants of Munc13-4 have been associated with Familial Hemophagocytic Lymphohistiocytosis Type 3 (FHL3), a rare autosomal recessive disorder characterized by impaired granule exocytosis^16,41^. Munc13-4 is also involved in exosome secretion in tumor cells^20^. Building upon this existing knowledge, our work contributes new insights by revealing that Munc13-4 is upregulated in various tumor cells and regulates multiple steps toward MVB exocytosis leading to exosome secretion, including MVB docking and fusion with the plasma membrane. This emphasizes its broader role in exocytosis across different cell types and highlights its potential as a biomarker for cancer diagnosis and prognosis. By regulating exosome secretion in tumor cells, Munc13-4 impacts the tumor microenvironment and facilitate communication between tumor and immune cells, influencing tumor growth, metastasis, and anti-tumor immunity.

The accumulation of MVBs in Munc13-4 knockout cells suggests that Munc13-4 is crucial for the docking of MVBs with the plasma membrane. While previous studies have underscored the importance of the Munc13-4–Rab27a complex in regulating secretory granule docking^19,23,24^, the underlying mechanisms remain unclear due to the absence of structural information. In this study, we present, for the first time, the cryo-EM structure of the Munc13-4–Rab27a complex, revealing a previously unexplored binding interface essential for complex stability. Specifically, residues F46, W73, and F88 in Rab27a, along with residues N739/T740 and V660/K661 in Munc13-4, maintain this stability and mediate MVB docking with the plasma membrane. Notably, the Rab27a-binding surface on Munc13-4 is located within the middle of the MUN domain (spanning residues 651–778), rather than in an N-terminal sequence connecting the C_2_A and MUN domain as reported previously^19,24^. Indeed, mutations in F46 and W73 in Rab27a and T740 in Munc13-4 have been previously linked to Griscelli Syndrome type 2 (GS2) ^42,43^ and FHL3^44^. Consistent with the structural data, introducing the mutations that disrupt the binding surface between Munc13-4 and Rab27a significantly increases the mobility of MVBs beneath the plasma membrane, which leads to impaired exosome secretion. Therefore, our results support the notion that Munc13-4 serves as the effector of Rab27a, stabilizing MVB docking close to the plasma membrane, thereby promoting exosome secretion. Together, Munc13-4–Rab27a-mediated MVB docking model may represent a common mechanism governing exosome secretion in various tumor cells.

Our previous study has identified that the SNARE complex that mediates MVB fusion with the plasma membrane in tumor cells is composed of syntaxin-4, SNAP-23 and VAMP-7^12^. Here, we observed an accelerated effect of Munc13-4 on SNARE complex assembly and SNARE-mediated membrane fusion, which is dependent on its interaction with SNAP-23 (Q_bc_-SNARE) and VAMP-7 (R-SNARE). This suggests an important role for Munc13-4 in chaperoning the proper conformation of SNAREs and/or in stabilizing the SNARE complex. The SNARE-chaperoning role of Munc13s, particularly the well-studied isoform Munc13-1, has been extensively documented^45–49^. While Munc13-4 shares similar domain structures with Munc13-1, their mechanisms in assisting SNARE complex assembly and membrane fusion may differ significantly. Syntaxin-4 is known to adopt a closed conformation similar to syntaxin-1^50,51^. However, Munc13-4 lacks the hydrophobic core (the NF pocket) in its MUN domain^46^, making it unlikely to catalyze the opening of syntaxin-4. In this regard, the activation of syntaxin-4 may depend on the interaction between its N-peptide and the corresponding SM protein, Munc18-3^52,53^. Our observation that Munc13-4 does not interact with syntaxin-4 further corroborates the notion. As a member of the CATCHR protein family, our finding that Munc13-4 binds to Q_bc_- and R-SNAREs is expected. For example, Munc13-1 binds to Q_bc_- SNARE SNAP-25^54^ and R-SNARE VAMP2^48^, aiding their assembly into the SNARE complex. The Dsl1 complex interacts with Q_b_-SNARE Sec20 and Q_c_-SNARE Use1, stabilizing the complex’s conformation^55^. The GARP complex subunit Vps51 binds to Q_c_-SNARE Tlg1, likely promoting its connection to SNARE bundles^56,57^. Similarly, the Exocyst complex subunit Sec6 interacts with Q_bc_- SNARE Sec9, facilitating both binary and ternary SNARE complex formation^58,59^. Together with our findings, these evidences suggest that Munc13-4 promotes SNARE complex formation during MVB fusion through mechanisms shared by various CATCHR family members.

The observation that Munc13-4 deletion does not alter the overall levels of PD-L1 on the plasma membrane but reduces the abundance of PD-L1 on secreted exosomes indicates a previously unrecognized role for Munc13-4 in cargo sorting within the endosomal system. HRS recognizes ubiquitinated proteins through its ubiquitin-interacting motif (UIM), which is essential for initiating cargo sorting during ESCRT-mediated MVB biogenesis^60,61^. While HRS was reported to sort cargoes such as interleukin-2 receptor beta (IL-2Rβ)^62^ and PD-L1^27^ independently of the UIM-ubiquitin interaction, the mechanism is not well understood. Our findings show that Munc13-4 independently binds both PD-L1 and HRS, but HRS cannot bind to PD-L1 without Munc13-4, highlighting Munc13-4’s critical role in recognizing PD-L1 before HRS interaction. This suggests that Munc13-4 mediates the recruitment of HRS to PD-L1, initiating a series of events that involve other ESCRT complexes and proteins required for the formation of PD-L1-containing ILVs. Therefore, the formation of a ternary complex consisting of Munc13-4, HRS, and Munc13-4-binding cargoes (e.g., PD-L1) may represent a novel ubiquitin-independent cargo sorting mechanism, where Munc13-4 and HRS work together to enable the proper sorting and packaging of cargoes into exosomes, facilitating its subsequent secretion. Further research is required to identify the full spectrum of exosomal cargo proteins sorted by Munc13-4. Overall, Munc13-4 works in conjunction with HRS, Rab27a, and SNAREs to establish an “assembly line” that effectively manages the processes of cargo sorting, packaging, trafficking, and release. This collaborative mechanism is critical for ensuring the efficient secretion of various proteins, including those involved in immune responses, highlighting the importance of Munc13-4 in exosomal biology.

In the tumor microenvironment, tumor-infiltrating lymphocytes can secret IFNγ to stimulate anti-tumor immune responses and induce tumor cell apoptosis^28,29^. However, tumor cells can exploit IFNγ to reduce anti-tumor immunity by increasing PD-L1 levels on exosomes. Our findings reveal that IFNγ exerts dual and opposing effects on PD-L1 sorting onto exosomes by modulating Munc13- 4 acetylation and HRS deubiquitylation. IFNγ triggers the translocation of the acetyltransferase CBP from the nucleus to the cytoplasm, leading to acetylation of K1062/K1079 at the C-terminal end of Munc13-4. The acetylation of K1062/K1079 disrupts the Munc13-4–PD-L1 interaction, thereby reducing PD-L1 sorting. Notably, these two lysine residues are unique to Munc13-4 and not conserved among Munc13s, underscoring Munc13-4’s distinct functional role in cargo recognition. Meanwhile, IFNγ stimulates HRS deubiquitylation, which enhances the HRS–PD-L1 interaction to increase PD-L1 sorting, likely due to a conformational change that leads to the activation of HRS^63^. These opposing effects indicate a nuanced regulatory mechanism where tumor cells may adapt to immune pressure by manipulating the sorting of PD-L1. Besides ubiquitylation, ERK-mediated phosphorylation of HRS also influences PD-L1 sorting^27^. Understanding these subtle regulatory mechanisms can help develop targeted therapies, such as modulating Munc13-4 acetylation through CBP or HDAC3, or altering HRS ubiquitylation and/or phosphorylation via NEDD4L, deubiquitinases or ERK. Recent studies suggest that exosomal PD-L1 levels in the blood of cancer patients could serve as a potential biomarker for predicting responses to immune checkpoint blockade (ICB) therapies^5–7^. The variability in PD-L1 levels on circulating exosomes among different patients may be linked to differences in Munc13-4 acetylation and HRS ubiquitylation/phosphorylation. Our findings could provide valuable insights for guiding ICB treatment decisions by analyzing Munc13-4 acetylation and HRS ubiquitylation in pathological tissue sections.

Tumor-derived exosomes containing PD-L1 have systemic immunosuppressive effects^8,39^, making the reduction of exosomal PD-L1 secretion a promising therapeutic strategy. Genetic blockade of overall exosome secretion has proven effective in slowing tumor growth and enhancing immunotherapy outcomes. For instance, loss of Rab27a in tumor cells inhibits exosome secretion, decreasing the release of all exosomal cargoes, which leads to reduced tumor growth and improved T cell anti-tumor activity^8,64^, similar to the effects observed with Munc13-4 in our study. However, while reducing overall exosome secretion can lessen their immunosuppressive effects, some evidence suggests that exosomes may also offer therapeutic benefits in immunotherapy. For example, exosomes can deliver tumor-associated antigens (TAAs) to dendritic cells, boosting T cell activation and promoting anti-tumor responses^65^. Exosomes carrying TAA-MHC complexes can directly stimulate antigen-specific T cell activation^66^. Therefore, selectively targeting PD-L1 sorting onto exosomes could eliminate their immunosuppressive effects while retaining their immunostimulatory potential. Our findings pinpoint the binding sites necessary for the Munc13-4–PD-L1 interaction that is specific for PD-L1 sorting. Disrupting this interaction with a designed peptide effectively reduces PD-L1 enrichment on exosomes without affecting overall exosome secretion. In vivo, this peptide treatment significantly enhances T cell function and inhibits tumor growth without major side effects. The peptide targets the PD-L1 motif that interacts with Munc13-4, allowing for selective PD-L1 targeting and minimizing the risk of immune dysfunction from non-specific peptide uptake by immune cells. Overall, this peptide holds great promise as a therapeutic agent for modulating immune responses.

In conclusion, we elucidate the Munc13-4-dependent mechanisms that govern the secretion of PD-L1 via exosomes in breast cancer cells, highlighting the functional complexity of Munc13-4 and enhancing our understanding of exosome biogenesis and secretion. Our in vivo findings offer valuable insights into the regulation of PD-L1 secretion, suggesting promising therapeutic strategies to improve patient outcomes in cancer treatment.

### Limitations of the study

Our study sheds light on Munc13-4’s role in tumor progression, specifically how it regulates PD-L1 secretion through exosomes. While we focus on breast cancer cells, the relevance of our findings to other cancer types needs further exploration. Although Munc13-4 is upregulated in various cancers, its precise functions in these contexts require additional validation. The interaction between Munc13- 4 and PD-L1 is crucial for sorting PD-L1 into exosomes. Future research should investigate the structural details of this complex to inform targeted interventions. In addition, there is an urgent need for specialized drug-delivery systems that can selectively target tumor tissues, protect our designed peptide that inhibits exosomal PD-L1 sorting from degradation, and avoid non-specific uptake by normal cells.

## Resource availability

### Lead contact

Requests for further information and resources should be directed to and will be fulfilled by the lead contact, Cong Ma.

### Materials availability

All materials generated in this study are available from the lead contact upon request.

### Data and code availability

Proteomics data generated in this study are available via ProteomeXchange with identifier PXD059882. Coordinate for the Munc13-4–Rab27a complex with GppNHp has been deposited in the RCSB Protein Data Bank (PDB: 9LA9). EM density maps for full map and composite map have been deposited in the Electron Microscopy Data Bank under accession codes EMD-62922 and EMD-63239, respectively. The home-written matlab script used for PLA data process is available from https://github.com/shenwang3333/PLA_Counting. All data reported in this paper are available from the lead contact upon request.

## Acknowledgments

We acknowledge Yueguang Rong (Huazhong University of Science and Technology) for providing the CRISPR-Cas9 plasmids and offering valuable suggestions on this study. We are grateful to Xiangliang Yang (Huazhong University of Science and Technology), Quan Chen (Nankai University) and Da Jia (Sichuan University) for their valuable advice on this study. We acknowledge the staff at the Cryo-EM Facilities of GIBH-CAS and the Guangzhou National Laboratory Bio-imaging Technology Platform for their assistance in cryo-EM sample preparation and data acquisition. This work was funded by the National Key R&D Program of China (2022YFA1206000), National Natural Science Foundation of China (32225024; 32421003; 92254302; 32401035; 32241021; 3231163669), National Science and Technology Major Project of China (2021ZD0202500), and Hubei Provincial Natural Science Foundation of China (2024AFA005).

## Author contributions

C.M. conceived the study. C.M., J.H., C.L. and D.L. designed experiments and wrote the paper. C.L. and D.L. performed the majority of the experiments. X.Z. expressed and purified proteins for structural studies and performed in vitro experiments. J.G. performed cryo-EM data collection and data processing. J.H. and J.G. built and refined models of Munc13-4–Rab27a complex. X.Z. helped with the isolation of exosomes, H.Z. helped with experiments related to protein modifications, S.W. helped with in vitro experiments and data analysis, and L.G. helped with animal experiments.

## STAR METHODS

### EXPERIMENTAL MODEL AND STUDY PARTICIPANT DETAILS

#### Bacterial strains

Escherichia coli strains BL21(DE3) (Thermo), DH10Bac (Gibco) and DH5α (Thermo) were cultured in Luria-Bertani (LB) broth at 37 °C with shaking at 200 rpm. The media was supplemented with appropriate antibiotics: ampicillin (100 µg/ml), kanamycin (50 µg/ml), gentamicin (7 µg/ml), or tetracycline (10 µg/ml), as necessary.

#### Cell culture

HEK293T and 4T1 cells were obtained from American Tissue Type collection (ATCC). SUM159 were obtained from Pcocell. HEK293T and SUM159 cells were cultured in DMEM medium (Gibco, 11965092) with 10% FBS (Gibco, A5670701) and 1% penicillin-streptomycin solution (Proteintech, PR40022). 4T1 cells were cultured in RPMI-1640 medium (Gibco, 11875093) supplemented with 10% FBS (Gibco, A5670701) and 1% penicillin-streptomycin (Proteintech, PR40022). These cells were cultured in a humidified incubator (Thermo) at 37°C with 5% CO2. Sf9 cells (Gibco) were cultured in SIM-SF Expression Medium (SinoBiological, MSF1) at 27°C, 125 rpm.

#### Mice

BALB/c, BALB/Nude and NOD/SCID mice (female, 5-7-week-old) were purchased from Beijing Vital River Laboratory Animal Technology Co., Ltd. (Beijing, China). The mice were housed in a controlled animal facility under consistent environmental conditions, including a room temperature of 22C±C1C°C, relative humidity of 40–70%, and a 12-hour light/dark cycle. Food and water were provided ad libitum. Mice were randomly assigned at the start of each experiment. All experimental procedures were conducted in compliance with the guidelines and with approval from the Institutional Animal Care and Use Committee (IACUC) of Tongji Medical College, Huazhong University of Science and Technology (Wuhan, China).

#### Patients’ samples

Formalin-fixed, paraffin-embedded human tissue arrays (HOrgC180PG01-2 and HBreD180Bc01-2) were obtained from Shanghai Outdo Biotech Co., Ltd. (China). The HOrgC180PG01-2 array comprises 180 tissue cores derived from 91 patients across 14 tumor types, and the HBreD180Bc01- 2 array includes 180 tissue cores from 150 patients with triple-negative breast cancer. Both tissue arrays were subjected to immunohistochemical (IHC) analysis, with detailed clinical information retrieved from the company’s website (https://www.superchip.com.cn/).

### METHOD DETAILS

#### Online data acquisition and analysis

The differential expression of Munc13-4 between tumor and normal tissues was analyzed utilizing TIMER2.0 (http://timer.cistrome.org/)^21,22^. The reported and predicted ubiquitin ligase (E3) of HRS was searched using UbiBrowser 2.0 (http://ubibrowser.bio-it.cn/ubibrowser_v3/)^31^.

#### Plasmids

The coding sequences for full-length human CD63, PD-L1, and Munc13-4 were inserted into the NPY-td-Orange2 vector. Full-length human Munc13-4, PD-L1, and various mutants of Munc13-4, including K1062R/K1079R, K1062Q/K1079Q, V660A/K661A, and N739G/T740G, along with truncated forms of Munc13-4 (residues 1–1048, 1–910, 1–782, 1–546, 1–287, and 1–108), were cloned into the pEGFP-N3 vector (Clontech). Similarly, full-length human HRS, STAM, and Rab27a, along with Rab27a mutants (F46S, W73S, and F88S), were cloned into the pEGFP-C1 vector (Clontech).

Constructs for full-length human Munc13-4, PD-L1, HRS, and mutant Munc13-4 (K1062R/K1079R, K1062Q/K1079Q), as well as truncated PD-L1 (residues 256–273), histone acetyltransferase domains of human GCN5 (residues 503–656), PCAF (residues 503–651), CBP (residues 1323–1700), P300 (residues 127–1663), TIP60 (residues 227–504), and HBO1 (residues 332–607), and full-length human NEDD4, ITCH, CBL, SH3RF1, PRKN, NEDD4L, STAMBPL1, CYLD, USP11, USP7, USP8, USP36, were generated in the pcDNA3.1- vector (Invitrogen) with an N-terminal Flag-tag. Histone deacetylase domains of human HDAC1 (residues 9–321), HDAC2 (residues 9–322), HDAC3 (residues 3–316), HDAC4 (residues 655–1084), HDAC5 (residues 684–1028), HDAC6 (residues 87–404 and 482–800), HDAC7 (residues 518–865), and HDAC8 (residues 14–324) were also cloned into the pcDNA3.1- vector with an N-terminal HA-tag. Coding sequence of full-length human NEDD4L were cloned into the pcDNA3.1- vector (Invitrogen) with an N-terminal Strep-tag.

Full-length human Munc13-4, Rab27a, and their respective mutants, including Munc13-4 (K1062R/K1079R, K1062Q/K1079Q, V660A/K661A, N739G/T740G) and Rab27a (F46S, W73S, F88S), were cloned into the pLV-EF1α-IRES-Hygro vector (Addgene). Full-length Munc13-4 and its mutants (V660A/K661A, N739G/T740G) were subcloned into the pFastBac^TM^HT B vector (Invitrogen).

Coding sequences for full-length human Rab27a, syntaxin-4, truncated syntaxin-4 (residues 1–275, Syx-4 ΔTM), SNAP-23 mutant (All cysteine residues were mutated to serine, SN-23-6CS), SN-23- 6CS (S161C), VAMP-7 SNARE motif (residues 123–187, A131C), VAMP-7 SNARE-TM motif (residues 123–208), and PD-L1 (residues 19–290) were cloned into the pET-28a vector (Novagen). Finally, full-length Rab27a, Rab5, Rab7, and their mutants (Rab27a (F46S, W73S, F88S)), Syx-4 ΔTM (1–275), SN-23-6CS, and VAMP-7 SNARE motif (123–187) were subcloned into the pGEX- 6P-1 vector (Cytiva).

#### Transfection

Recombinant bacmid was transfected into Sf9 cells using the X-tremeGENE™ 9 DNA Transfection Reagent (Roche). For all other plasmid transfections, Hieff Trans® Liposomal Transfection Reagent (Yeasen, 40802ES08) was employed, following the manufacturer’s instructions. Cells were prepared for subsequent experiments 24–36 hours post-transfection.

#### Cell viability assay

Cell proliferation of control and Munc13-4 knockout 4T1 cells was assessed using a CCK-8 kit (Vazyme, A311-01) according to the manufacturer’s instructions. Briefly, the same number of control and Munc13-4 knockout cells were plated onto 96-well culture plates, and cell viability was measured at 0, 24, and 48 hours using the CCK-8 kit.

#### Histological analyses

Tissue arrays were deparaffinized with heat at 60 °C for 30 min followed by two 15-minute washes with xylene. Then, the paraffin sections were rehydrated by washing for 5 min in absolute ethanol I, absolute ethanol II, 85% alcohol, 75% alcohol and distilled water in sequence. Following the procedures outlined in the “PTLink Quick Operation Guide” (Dako), slides were subjected to antigen retrieval using the specified instrument. Upon completion, slides were immersed in distilled water at room temperature for natural cooling for a minimum of 10 minutes. Subsequently, the slides were rinsed with PBST buffer. The diluted Munc13-4 primary antibody working solution (1:50, Santa Cruz, sc-271300) was applied, and the slides were incubated overnight at 4°C. The next day, the slides were removed from refrigeration and allowed to equilibrate to room temperature for 45 minutes before being washed with PBST buffer. Automated staining, including blocking, secondary antibody binding, and DAB color development, was performed using the DAKO automated immunohistochemistry staining system according to the ”Autostainer Link 48 User Guide.” Counterstaining was conducted using hematoxylin for 1 minute, followed by immersion in 0.25% hydrochloric acid alcohol (prepared with 400 ml of 70% ethanol and 1 ml of concentrated hydrochloric acid) for no less than 2 seconds. The slides were rinsed under running water for a minimum of 2 minutes, air-dried at room temperature, and mounted using neutral resin. Digitization of the slides was performed at ×20 magnification using the Aperio XT Scanner (Leica). To evaluate the protein expression levels of Munc13-4 in breast cancer tissues and adjacent normal tissues, immunohistochemistry (IHC) images were analyzed using IHC Profiler, an open-source plugin for ImageJ. The staining intensity for Munc13-4 was scored using a four-tier scoring system: 0 (negative), 1 (low positive), 2 (positive), and 3 (high positive).

Samples of mouse heart, liver, spleen, lung and kidney were fixed overnight in 4% formalin, embedded in paraffin, and cut into 4 mm consecutive sections. The paraffin sections were sequentially immersed in Environmental-Friendly Dewaxing Transparent Liquids I and II (Servicebio, G1128) for 20 minutes each, followed by treatment with anhydrous ethanol I and II for 5 minutes each. Subsequently, the sections were immersed in 75% ethanol for 5 minutes and thoroughly rinsed with tap water. Hematoxylin and eosin (H&E) staining was performed using the Hematoxylin-Eosin (H&E) HD Constant Dye Kit (Servicebio, G1076) according to the manufacturer’s instructions. The sections were then dehydrated through a graded series of absolute ethanol solutions (I, II, and III) for 2 minutes each, followed by sequential immersion in normal butanol I and II for 2 minutes each and clearing in xylene I and II for 2 minutes each. Finally, the sections were sealed with neutral gum and scanned using the NanoZoomer S360 Digital Slide Scanner (Hamamatsu).

For tissue immunofluorescence assays, paraffin-embedded tumor tissues from orthotopic mouse models of breast cancer sections were deparaffinized and rehydrated through a graded ethanol series, followed by washing in distilled water. Antigen retrieval was performed using EDTA Antigen Retrieval Solution (Beyotime, P0085) under high temperature and pressure conditions, and the sections were allowed to cool to room temperature before washing in Tris-buffered saline with 0.05%

Tween-20, pH 7.4 (TBST). Endogenous peroxidase activity was blocked using 3% H2O2, followed by washing in distilled water. The sections were then encircled with a hydrophobic pen and incubated with 10% goat serum (Boster, AR1009) at 37°C for blocking. For the first staining, a CD4 primary antibody (Abcam, RM1013, 1:50) diluted in TBST was applied, and the sections were incubated overnight at 4°C. After washing, a secondary antibody Goat Anti-Rabbit IgG H&L (HRP) (Abcam, ab205718, 1:4000) was added and incubated at 37°C. Tyramide signal amplification (TSA) staining was performed using iFluor^®^ 488 tyramide working solution (AAT Bioquest, 45100), followed by washing in TBST. The slides underwent a second round of antigen retrieval in Improved Citrate Antigen Retrieval Solution (Beyotime, P0083) using a microwave, were cooled to room temperature, and washed again. Blocking was repeated with 10% goat serum at 37°C.For the second staining, a CD8 primary antibody (Abcam, RM1129, 1:100) diluted in TBST was applied, and the sections were incubated overnight at 4°C. Following washing, the HRP-conjugated secondary antibody was added and incubated, and TSA staining was conducted using Cy3 tyramide working solution (AAT Bioquest, 11065). Nuclear staining was performed with DAPI (Solarbio, C0060) in the dark, followed by washing in TBST. Finally, the sections were mounted with Fluoromount-G^®^ (SouthernBiotech, 0100-01) and stored at 4°C in the dark. The sections were imaged using Pannoramic SCAN II (3D HISTECH).

#### iTRAQ-Based quantitative proteomics

4T1 cells (control and Munc13-4 knockout) were plated in triplicate for proteomics. To extract proteins, an appropriate volume of SDS-free L3 buffer supplemented with final concentration of 1 × Cocktail (EDTA contained) was added to the sample. The mixture was incubated on ice for 5 minutes, followed by the addition of DTT to achieve a final concentration of 10 mM. Ultrasonic disruption was performed to lyse the sample, and the lysate was centrifuged at 25,000 × g and 4°C for 15 minutes to remove insoluble debris. The supernatant was collected and further treated with DTT (final concentration of 10 mM), followed by incubation in a water bath at 56°C for 1 hour.

Subsequently, iodoacetamide was added to the solution to a final concentration of 55 mM, and the mixture was incubated in the dark for 45 minutes. A second centrifugation at 25,000 × g and 4°C for 15 minutes was performed, and the supernatant, containing the extracted protein solution, was collected for downstream analyses.

Protein samples (100 µg each) were digested with Trypsin Gold (Promega, V5280) at a protein-to-trypsin ratio of 20:1 (w/w) at 37°C for 16 hours. The resulting peptides were dried via vacuum centrifugation and reconstituted in 0.5 M TEAB. iTRAQ labeling was conducted following the manufacturer’s protocol for the 4-plex iTRAQ reagent kit (Sigma-Aldrich, 4374321). The labeled samples were combined in equal proportions and fractionated using high-performance liquid chromatography (HPLC) on a Thermo DIONEX Ultimate 3000 BioRS system equipped with a Durashell C18 column (5 µm, 100 Å, 4.6 × 250 mm, Welch Materials). A total of 20 fractions were collected for further analysis.

Peptides separated by liquid chromatography were ionized using a nanoESI source and analyzed on a Q-Exactive HF X mass spectrometer (Thermo Fisher Scientific) operating in data-dependent acquisition (DDA) mode. Instrument parameters were configured as follows: the ion source voltage was set to 1.9 kV; the MS1 scan range was 350–1,500 m/z with a resolution of 60,000; and the MS2 scan range started at a fixed m/z of 100 with a resolution of 15,000. Precursor ion selection criteria included charge states between 2+ and 6+ and the top 20 most intense ions with signal intensities exceeding 10,000. Fragmentation was performed using higher-energy collisional dissociation (HCD), and the resulting fragments were detected in the Orbitrap analyzer. The dynamic exclusion duration was set to 30 seconds, and the automatic gain control (AGC) targets were 3 × 10^6^ for MS1 and 1 × 10^5^ for MS2.

Protein identification was conducted utilizing the Mascot search engine (version 2.3.02; Matrix Science) against the Mus musculus subset of the NCBI non-redundant (NR) sequence databases. The search parameters were configured as follows: monoisotopic mass, peptide mass tolerance of 20 ppm, fragment mass tolerance of 0.05 Da, trypsin as the digestion enzyme, allowance for one missed cleavage, and charge states of +2 and +3 for peptides. Variable modifications included Gln-> pyro-Glu (N-terminal Q), oxidation (M), and deamidation (NQ), while fixed modifications comprised carbamidomethylation (C) and iTRAQ8plex labeling (N-terminal and K). Protein quantification was performed using the automated software IQuant. Peptides with a confidence interval of 95% were filtered based on a 1% false discovery rate (FDR), and confident proteins were required to include at least one unique peptide. Quantitative protein ratios were weighted and normalized using the median ratio in Mascot. Differentially expressed proteins (DEPs) between the control and Munc13-4 knockout groups were identified via t-tests, with results subjected to a 5% FDR correction. Proteins with expression fold changes ≥ 1.2 or ≤ 0.83 were classified as DEPs. Additionally, KEGG pathway enrichment analysis was performed for DEPs, and a heatmap was generated using an online platform for data analysis and visualization (https://www.bioinformatics.com.cn/)^67^.

#### Cell treatment with IFN**γ**, PR-619 and peptide

To explore the effects of IFNγ on the expression and exosomal sorting of PD-L1, 100 ng/ml Recombinant Human IFN-gamma Protein (Abclonal, RP01038) was added to SUM159 cells, and 100 ng/ml Recombinant Mouse IFN-gamma Protein (Abclonal, RP01070) was added to 4T1 cells, incubating 24 hours. To examine the effect of HRS ubiquitylation on the sorting of PD-L1 onto exosomes under IFNγ stimulation, SUM159 cells were co-treated with 100 ng/ml Recombinant Human IFN-gamma Protein (Abclonal, RP01038) and 8 μM PR-619 (MCE, HY-13814) for 24 hours. To investigate the effects of the designed peptide on protein interactions and exosomal PD-L1 sorting, HEK293T and SUM159 cells were incubated with 10 μg/ml P-pep (Homo sapiens) or S-pep.

#### Western blot

Cells or exosomes were lysed on ice in RIPA buffer (50 mM Tris-HCl, pH 7.5, 150 mM NaCl, 1% Triton X-100) supplemented with a protease inhibitor cocktail (Topscience, C0001). Following a 20- minute incubation, the lysates were centrifuged at 4°C, 12,000 × g for 10 minutes. The total protein concentration in the supernatant was determined using a BCA Protein Quantification Kit (Yeasen, 20201ES76) to ensure consistent loading of different samples. Protein samples were then denatured by heating in diluted 1× SDS-PAGE Sample Loading Buffer (Yeasen, 20315ES05) for 10 minutes at 100°C. Proteins were separated by SDS-PAGE and subsequently transferred to PVDF membranes (Millipore, ISEQ00010). The membranes were blocked with 5% non-fat bovine milk in Tris-Cl buffer (150 mM NaCl, pH 7.2) containing 0.1% Tween-20, followed by incubation with the indicated primary antibody and a subsequent incubation with an HRP-conjugated secondary antibody.

Immunodetection was carried out using the Super Sensitive ECL Luminescence Reagent (Meilunbio, MA0186-2). The integrated density of the blot bands was quantified and analyzed using ImageJ and Prism 6.0 software to assess the relative protein levels. The primary antibodies used in western blot assays are as follows: Munc13-4 Antibody (C-2) (Santa Cruz, sc-271300, 1:1000), PD-L1/CD274 Rabbit mAb (Abclonal, A19135, 1:2000), HGS Polyclonal antibody (Proteintech, 10390-1-AP, 1:10000), β-Actin Rabbit mAb (High Dilution) (Abclonal, AC026, 1:100000), Alix Monoclonal antibody (Proteintech, 67715-1-Ig, 1:5000), CD63 Antibody (MX-49.129.5) (Santa Cruz, sc-5275, 1:1000), CD81 Antibody (B-11) (Santa Cruz, sc-166029, 1:1000), Mouse anti GFP-Tag mAb (Abclonal, AE012, 1:10000), Rabbit anti GFP-Tag pAb (Abclonal, AE011, 1:10000), DYKDDDDK tag Polyclonal antibody (Proteintech, 20543-1-AP, 1:10000), StrepII Tag Mouse Monoclonal Antibody (Beyotim, AF2924, 1:1000), Mouse anti HA-Tag mAb (Abclonal, AE008, 1:5000), Pan Acetylation Monoclonal antibody (Proteintech, 66289-1-Ig, 1:1000), Ubiquitin Antibody (P4D1) (Santa Cruz, sc-8017, 1:1000), pan Phospho-Serine/Threonine Rabbit Polyclonal Antibody (Beyotim, AF5725, 1:1000), Pan Phospho-Tyrosine Mouse mAb (Abclonal, AP0973, 1:1000), CBP/KAT3A/CREBBP Antibody (C-1) (Santa Cruz, sc-7300, 1:1000), P300 Antibody (F-4) (Santa Cruz, sc-48343, 1:1000), Histone Deacetylase 3 (HDAC3) Antibody (A-3) (Santa Cruz, sc-376957, 1:1000), Histone Deacetylase 4 (HDAC4) Antibody (B-5) (Santa Cruz, sc-365093, 1:1000). The secondary antibodies used in western blot experiments are as follows: HRP-conjugated Goat anti-Rabbit IgG (H+L) (Abclonal, AS014, 1:10000), HRP-conjugated Goat anti-Mouse IgG (H+L) (Abclonal, AS003, 1:10000). The integrated density of blot strips was analyzed by ImageJ software to characterize the relative protein level.

#### Generation of gene-edited cell lines

To generate gene knockout cell lines, the CRISPR-Cas9 system was employed. HEK293T cells were transfected with the lentiCRISPR v2 plasmid, which contained a single-guide RNA (sgRNA) targeting the gene of interest, along with the psPAX2 and pMD2.G plasmids to produce lentivirus.

After 36–48 hours of transfection, the culture medium of the HEK293T cells was collected, centrifuged to remove cell debris, and the supernatant containing the lentivirus-sgRNA was used to infect SUM159 or 4T1 cells. Following 48 hours of infection, cells were selected with puromycin (MCE, HY-B1743) at a concentration of 2.0 μg/mL for 5–7 days. Limiting dilution was then performed to isolate single-cell clones from the infected SUM159 or 4T1 cells. Cells infected with lentivirus produced by HEK293T cells co-transfected with an empty lentiCRISPR v2 vector, psPAX2, and pMD2.G plasmids served as controls. Successful knockout of the target SNARE protein was verified by western blot analysis.

The sgRNAs used in this study are as follows:

Mus-Munc13-4: GTGGCCTTCAGGCAAAATAC Hs-Munc13-4: TGAAGGTCTCGTCCCAGACG Hs-Rab27a: CCAAAGCTAAAAACTTGATG

Hs-HRS: CTGCCTGCAGAGACAAGTGG Hs-P300: GTTCAATTGGAGCAGGCCGA Hs-CBP: CGCGTGACCAGTCATTTGCG Hs-HDAC3: GGTGAAGCCTTGCATATTGG

Hs-HDAC4: GGAGCCCATTGAGAGCGATG Hs-NEDD4L: GGAGCCCATTGAGAGCGATG

For the generation of gene-complemented cell lines, the pLV-EF1a-IRES-Hygro plasmid containing the full-length sequence of the gene of interest was utilized. HEK293T cells were transfected with the pLV-EF1a-IRES-Hygro plasmid along with the psPAX2 and pMD2.G plasmids to produce lentivirus. At 36–48 hours post-transfection, the culture medium was collected, centrifuged to remove cell debris, and the resulting lentivirus-containing supernatant was used to infect the gene knockout SUM159 cells. Infected cells were selected using Hygromycin B (Sangon Biotech, A100607) at a final concentration of 500 μg/ml for 5–7 days. Single-cell clones were subsequently isolated through limiting dilution.

#### Immunoprecipitation (IP) and co-IP

To investigate the post-transcriptional modifications of Munc13-4 and HRS, IP assays were conducted. Equal numbers of SUM159 cells were seeded onto 15-cm culture dishes, with one plate treated with 100 ng/ml IFNγ (ABclonal, RP01038) for 24 hours. Following the incubation, cells were lysed on ice using RIPA buffer (50 mM Tris-HCl, pH 7.5, 150 mM NaCl, 1% Triton X-100) supplemented with a protease inhibitor cocktail (Topscience, C0001). The lysates were then centrifuged at 4°C for 10 minutes at 12,000 × g. The supernatant was incubated overnight at 4°C with Munc13-4 Antibody (C-12) (Santa Cruz, sc-271301, 1:100) or HGS Polyclonal antibody (Proteintech, 10390-1-AP, 1:500), in conjunction with Protein A/G magnetic beads (Biolinkedin, L- 1004), on a rotator. To identify the acetyltransferase of Munc13-4, GFP-tagged Munc13-4 was co-expressed with Flag-labelled GCN5 503–656, PCAF 503–651, CBP 1323–1700, P300 127–1663, TIP60 227–504 or HBO1 332–607 in HEK293T cells. To investigate the type of deacetylase, HEK293T cells co-expressing GFP- Munc13-4 and Flag- CBP 1323–1700 was treated with 0.2% DMSO, 5mM nicotinamide (NIA) (MCE, HY-B0150) or 1 μM trichostatin A (TSA) (MCE, HY- 15144) for 24 hours. To identify the deacetylase of Munc13-4, GFP- Munc13-4 and Flag- CBP 1323–1700 was co-expressed with HA-fused HDAC1 9–321, HDAC2 9–322, HDAC3 3–316, HDAC4 655–104, HDAC5 684–1028, HDAC6 87–404, HDAC6 482–800, HDAC7 518–865 or HDAC8 14–324 in HEK293T cells. To identify the E3 ligase of HRS, GFP-HRS was co-expressed with Flag-tagged NEDD4, ITCH, CBL, SH3RF1, PRKN or NEDD4L in HEK293T cells. To identify the deubiquitinase of HRS, GFP-HRS and strep-NEDD4L was co-expressed with Flag-fused STAMBPL1, CYLD, USP11, USP7, USP8 or USP36 in HEK293T cells. The transfection of plasmids mentioned above was performed using Hieff Trans^®^ Liposomal Transfection Reagent (Yeasen, 40802ES03) according to the manufacturer’s protocol.GFP-Munc13-4 or GFP-HRS was immunoprecipitated from cell lysates using anti-GFP magnetic beads (Biolinkedin, L-1016) according to the manufacturer’s guidance. After incubation, the beads were washed three times with PBS. Subsequently, 1× SDS-PAGE Sample Loading Buffer (Yeasen, 20315ES05) was added, and the samples were heated at 100°C for 10 minutes. The protein samples were then analyzed by western blotting.

The deletion of either CBP or HDAC3 in SUM159 cells resulted in a significant decrease in Munc13-4 expression. To examine the effects of CBP and HDAC3 knockout on Munc13-4 acetylation, Munc13-4 was enriched from a cell population three times larger in the knockout groups compared to the control cells. The amount of immunoprecipitated Munc13-4 used for acetylation analysis was standardized based on the results of preliminary western blot analysis.

To investigate protein-protein interactions, co-IP assays were conducted. Recombinant plasmids were transfected into SUM159 or HEK293T cells using Hieff Trans^®^ Liposomal Transfection Reagent (Yeasen, 40802ES03). GFP-fused proteins were then enriched from cell lysates using anti-GFP magnetic beads (Biolinkedin, L-1016). Following enrichment, the beads were washed three times with PBS and subsequently heated in 1× SDS-PAGE Sample Loading Buffer (Yeasen, 20315ES05) at 100°C for 10 minutes. Next, the samples were analyzed by western blotting.

#### Isolation of extracellular vesicles (EVs) / exosomes

To isolate EVs / exosomes from cell culture medium, equal numbers of cells with different genotypes were plated on 15-cm culture dishes. After cell adherence, the cells were rinsed with PBS and refreshed with an equivalent volume of DMEM containing 10% exosome-depleted FBS (bovine exosomes were removed by overnight centrifugation at 100,000 × g), followed by a 24-hour incubation. Then exosomes were isolated from the medium using a sequential centrifugation protocol. Briefly, the culture medium was first centrifuged at 4°C, 300 × g for 10 minutes to remove cells, followed by centrifugation at 4°C, 2000 × g for 20 minutes to eliminate cell debris. The medium was then centrifuged at 4°C, 10,000 × g for 30 minutes to remove larger extracellular vesicles. The resulting supernatant was ultracentrifuged at 100,000 × g for 70 minutes (Beckman Type 70Ti rotor). The exosome pellet was washed in cold PBS and subjected to another round of ultracentrifugation at 100,000 × g for 70 minutes to further purify the vesicles. Finally, the exosome pellet was resuspended in PBS or RIPA buffer for subsequent analysis.

#### Optiprep^TM^ density gradient centrifugation

Pellets of EVs, obtained through ultracentrifugation from cell culture supernatants, were washed and resuspended in 200 µL of buffer containing 0.25 M sucrose, 10 mM Tris-Cl, and 1 mM EDTA (pH 7.4). The suspension was then transferred to an SW55Ti rotor tube (Beckman, 344090), mixed in a 1:1 ratio with a 60% (wt/vol) Optiprep™ stock solution, and sequentially layered with 160 µL of a 20% (wt/vol) Optiprep™ solution and 150 µL of a 10% (wt/vol) Optiprep™ solution. Tubes were centrifuged for 1 hour at 4°C, 350,000 × g in an SW55Ti rotor (stopping without break). Following centrifugation, six 100 µL fractions were collected from the top of the gradient. These fractions were diluted with 600 µL of PBS and subjected to a second round of centrifugation for 1 hour at 4°C, 100,000 × g. The resulting pellets from the concentrated fractions were resuspended in 20 µL of PBS and analyzed by western blotting.

#### Transmission electron microscopy (TEM)

The morphology of EVs was characterized by TEM. 20 μL of EVs suspension was carefully deposited onto a copper grid (EMCN, BZ11262a) and incubated for 3–5 minutes. Excess liquid was then removed with filter paper. Subsequently, 2% uranyl acetate was applied to the copper grid for 2–3 minutes, after which the excess solution was absorbed using filter paper, and the sample was allowed to air-dry at room temperature. The samples were then observed with TEM (HITACHI, HT7800). The number of ILVs and MVBs and the percentage of MVB-lysosome hybrids among total MVBs was scored manually and the diameter of MVBs was measured by ImageJ software.

#### Nanoparticle tracking analysis (NTA)

The number and size of exosomes were directly measured using the NS300 instrument (Malvern) equipped with a high-sensitivity sCMOS camera. Particle tracking and sizing were performed automatically based on Brownian motion and the diffusion coefficient. Three 60-second videos were recorded for each sample. The videos were then analyzed using NanoSight NTA software, which tracked the center of each particle under Brownian motion and calculated the average distance traveled by the particles on a frame-by-frame basis.

#### nanoLCMS/MS analysis

nanoLCMS/MS analysis was performed for the identification of Munc13-4 acetylation. 24 hours following the co-expression of GFP-Munc13-4 and Flag-CBP 1323–1700 in HEK293T cells, GFP- Munc13-4 was immunoprecipitated from cell lysates using anti-GFP magnetic beads (Biolinkedin, L-1016), following the manufacturer’s protocol. The beads were subsequently washed three times with PBS, after which diluted 1 × SDS-PAGE Sample Loading Buffer (Yeasen, 20315ES05) was added, and the samples were heated at 100°C for 10 minutes. Proteins separated by SDS-PAGE were subjected to trypsin digestion (Promega, V5280) in 100 mM NH4HCO3 overnight at 37°C. The resulting peptides were extracted with extraction buffer (1:2, vol/vol, 5% formic acid/acetonitrile) and then vacuum-dried.

A total of 200 ng of peptides were separated and analyzed using a nano-UPLC system (Evosep One) coupled to a timsTOF Pro2 mass spectrometer (Bruker) equipped with a nano-electrospray ionization source. Peptide separation was achieved on a reversed-phase column (PePSep C18, 1.9 µm, 150 µm × 15 cm, Bruker) with mobile phases consisting of H2O containing 0.1% formic acid (phase A) and acetonitrile (ACN) with 0.1% formic acid (phase B). A 44-minute gradient was used for the separation. Data acquisition was performed in DDA PaSEF mode, with the mass spectrometer scanning within a range of 100 to 1700 m/z for MS1. During PASEF MS/MS acquisition, the collision energy was linearly increased in correlation with ion mobility, ranging from 20 eV (1/K0 = 0.6 Vs/cm²) to 59 eV (1/K0 = 1.6 Vs/cm²).

The raw MS data files provided by the vendor were processed using SpectroMine software (version 4.2.230428.52329) in conjunction with the integrated Pulsar search engine. The MS spectra were queried against the species-specific UniProt FASTA database for *Homo sapiens* (uniprot_Homo sapiens_9606_reviewed_2023_09.fasta), with carbamidomethylation of cysteine (C) set as a fixed modification, and oxidation (M) and acetylation at the protein N-terminus as variable modifications. Trypsin was employed as the protease, with a maximum allowance of two missed cleavages. A false discovery rate (FDR) of 0.01 was applied at both the peptide-spectrum match (PSM) and peptide levels. Peptide identification was performed with an initial precursor mass tolerance of 20 ppm. All other parameters were left at their default settings.

#### Proximity ligation assay (PLA)

SUM159 cells were plated onto the wells of 29-mm glass-bottom dishes (Cellvis, D29-10-0-N). After fixation with 4% (wt/vol) paraformaldehyde at room temperature for 15 minutes, the cells were permeabilized with 0.2% (vol/vol) Triton X-100 in PBS for 10 minutes at room temperature, followed by three washes with PBS. Subsequently, in situ proximity ligation assays were performed using the Duolink^®^ In Situ Red Starter Kit Mouse/Rabbit (Sigma-Aldrich, DUO92101), following the manufacturer’s protocol. This included blocking, primary antibody incubation, Duolink^®^ PLA probe incubation, ligation, amplification, final washing, and nuclear staining in sequence. The cell samples were imaged using an FV3000 Confocal Laser Scanning Microscope (Olympus) equipped with a 60× oil-immersion objective (NA 1.42). DAPI was excited using a 405 nm laser, and Duolink^®^ In Situ Detection Reagents Red were excited with a 594 nm laser. To determine the interaction between Munc13-4 and PD-L1, Munc13-4 Antibody (C-2) (Santa Cruz, sc-271300, 1:200) and PD-L1/CD274 (C-terminal) Polyclonal antibody (Proteintech, 28076-1-AP, 1:200) were used. To explore the interaction between HRS and PD-L1, HGS Polyclonal antibody (Proteintech, 10390-1- AP, 1:500) and PD-L1/CD274 Monoclonal antibody (Proteintech, 66248-1-Ig, 1:200) were employed. Data were processed by home-written matlab script (https://github.com/shenwang3333/PLA_Counting).

#### Orthotopic mouse models of breast cancer and treatments

Mice were anesthetized via intraperitoneal injection of pentobarbital sodium at a dose of 40 mg/kg. Orthotopic breast cancer models were established by injecting control or Munc13-4 knockout 4T1 cells (3 × 10^5^ cells per mouse) into the right fourth mammary fat pad of BALB/c, BALB/c Nude, or NOD/SCID female mice. Tumor dimensions were measured every other day starting on either day 4 or day 6 post-inoculation using a digital caliper, and tumor volume was calculated using the formula: (width² × length × 0.5). At the end of the observation period, mice were euthanized, and tumors were harvested, weighed, and photographed for further analysis.

Exosomes (1 × 10^9^ particles) secreted by control 4T1 cells were pre-incubated with either an IgG isotype control antibody (Bioxcell, BE0090, 1:100) or *InVivo*MAb anti-mouse PD-L1 antibody (Bioxcell, BE0101, 1:100). To remove unbound antibodies, the exosome-antibody complexes were subjected to ultracentrifugation at 100,000 × g for 70 minutes at 4°C. The purified exosomes were then intravenously injected into mice via the tail vein. Injections were administered every other day for a total of nine treatments.

Peptide treatments were performed using P-pep (Mus musculus) or S-pep, administered via intraperitoneal injection at a dosage of 100 µg per mouse. Once the tumor volume reached approximately 100 mm³, peptides were administered every other day, with a total of six injections.

For immune checkpoint blockade (ICB) studies, mice were treated with IgG isotype control antibody (Bioxcell, BE0090, 100 µg/mouse), *InVivo*MAb anti-mouse PD-L1 antibody (Bioxcell, BE0101, 100 µg/mouse) or *InVivoMAb* anti-mouse PD-1 antibody (Bioxcell, BE0146, 100 µg/mouse). Antibodies were administered via intraperitoneal injection every three days for a total of six treatments.

#### Immune profiling

To analyze T cell infiltration and activation, tumors, spleens, and tumor-draining lymph nodes (TDLNs) were excised from orthotopic mouse models of breast cancer. Tumor tissues were minced into small pieces and incubated with RPMI 1640 medium containing 1 mg/ml collagenase D (Roche, COLLD-RO) and 0.2 mg/ml DNase I (BioFroxx, 112MG010) at 37°C for 1 hour, followed by mechanical dissociation using a mesh cell strainer. The cells were then centrifuged at 500 × g for 5 minutes, washed with PBS, and treated with red blood cell (RBC) lysis buffer (BD Biosciences, 555899) to remove RBCs. The resulting cell suspension was filtered twice through a 70 μm nylon mesh to obtain single-cell suspensions. Lymphocytes from the spleens and TDLNs were isolated by mechanically squashing the tissues through a 70 μm mesh and removing RBCs.

For stimulation, the single-cell suspensions were incubated with RPMI 1640 medium containing Leukocyte Activation Cocktail (BD Biosciences, 550583, 1:1000) at 37°C for 6 hours. After staining with viability dye FVS 575V (BD Biosciences, 565694, 1:1,000) to exclude dead cells, the cells were stained with the following antibodies. For surface marker analysis, cells were stained with anti- CD45-APC-Cy7 (BD Biosciences, 557659, 1:50), anti-CD3-BV421 (BD Biosciences, 562600, 1:50), anti-CD4-Alexa Fluor 700 (BD Biosciences, 557956, 1:50), and anti-CD8-Percp-Cy5.5 (BD Biosciences, 551162, 1:50). For intracellular cytokine staining, cells were fixed and permeabilized using Fix/Perm buffer (BD Biosciences, 562574) and Perm/Wash buffer (BD Biosciences, 562574), then re-stained with anti-KI67-BV510 (BD Biosciences, 563462, 1:50), anti-IFN-γ-BV650 (BD Biosciences, 563854, 1:50), or anti-Granzyme B-FITC (Invitrogen, 11-8898-82, 1:500). Flow cytometric analysis was performed using the CytoFLEX flow cytometer.

#### CD8^+^ T cell suppression assay

To block PD-L1 on the exosome surface, equal quantities of exosomes isolated from control and Munc13-4 knockout 4T1 cells were incubated with PD-L1 blocking antibodies (Bioxcell, BE0101, 1:100) or IgG isotype control antibody (Bioxcell, BE0090, 1:100) at room temperature for 5 hours. After incubation, the exosomes were washed with 25 ml PBS and subjected to ultracentrifugation to remove unbound antibodies.

Mouse CD8^+^ T cells were purified from splenocytes using the Mouse CD8 T Cell Isolation Kit (Vazyme, CS103-01) and stimulated for 24 hours with anti-CD3 (Biolegend, 300301, 1 μg/ml) and anti-CD28 (Biolegend, 117003, 1 μg/ml) antibodies. Post-stimulation, the CD8C T cells were incubated with the preprocessed exosomes for 16 hours in the continued presence of anti-CD3 and anti-CD28 antibodies.

Following treatment, CD8^+^ T cells were harvested, stained with anti-CD8-Percp-Cy5.5 (BD Biosciences, 551162, 1:50), and permeabilized using Fix/Perm buffer and Perm/Wash buffer (BD Biosciences, 562574). Fixed cells were subsequently stained with anti-Granzyme B-FITC (Invitrogen, 11-8898-82, 1:500) and analyzed by flow cytometry.

#### T cell-mediated tumor cell killing assay

Spleens were aseptically harvested from BALB/c mice and placed in sterile petri dishes containing cold PBS. The spleens were gently disrupted by grinding against a 70 μm cell strainer using a sterile syringe plunger, and the resulting cell suspension was centrifuged at 500 × g for 5 minutes at 4°C. The pellet was resuspended in 5 ml of PBS, and the suspension was carefully layered onto 5 ml of Ficoll-Paque™ PLUS (Cytiva, 17144002) in a 15 ml conical tube without mixing. The gradient was centrifuged at 1000 × g for 20 minutes at room temperature with the centrifuge brake turned off. The mononuclear cell layer at the interface was carefully collected using a pipette, transferred to a fresh tube, and washed twice with PBS by centrifugation at 500 × g for 5 minutes at 4°C to remove residual Ficoll. The purified lymphocytes were then resuspended and stimulated with anti-CD3 (Biolegend, 300301, 1 μg/ml) and anti-CD28 (Biolegend, 117003, 1 μg/ml) antibodies for 24 hours. Subsequently, the stimulated lymphocytes were co-cultured with adherent control or Munc13-4 knockout 4T1 cells in 96-well plates at an effector-to-target (E: T) ratio of 1:1 for 24 hours. The viability of control and Munc13-4 knockout 4T1 cells was assessed using the Cell Counting Kit-8 (Vazyme, A311-01) following the manufacturer’s instructions.

#### Quantitative reverse transcription (qRT)-PCR assay

Total RNA was extracted from each sample using the AFTSpin Tissue/Cell Fast RNA Extraction Kit for Animal (Abclonal, RK30120). The isolated RNA was eluted in nuclease-free water and reverse-transcribed into complementary DNA (cDNA) using the ABScript II cDNA First-Strand Synthesis Kit (Abclonal, RK20400). The resulting cDNA samples were subjected to quantitative PCR on a QuantStudio™ 6 Pro Real-Time PCR system using SYBR Green (Abclonal, RK21203) for detection. The primers used for qPCR were as follows:

Hs-Munc13-4 qPCR forward primer: CCCTTTGTCCAGCTGACCTT Hs-Munc13-4 qPCR reverse primer: AGCAGGCACCAGGAATTCAA Hs-Actin qPCR forward primer: GCCGCCAGCTCACCAT

Hs-Actin qPCR reverse primer: AGGAATCCTTCTGACCCATGC

#### Fluorescence imaging

To analyze the colocalization between PD-L1 and CD63/LAMP1, Munc13-4 knockout and control SUM159 cells were seeded onto glass-bottom dishes and transfected with the pEGFP-N3-PD-L1 plasmid along with either NPY-td-Orange2-CD63 or NPY-td-Orange2-LAMP1 plasmids. Following 24–36 hours of transfection, cells were fixed with 4% paraformaldehyde (PFA) for 15 minutes. For immunofluorescence staining, cells were permeabilized with 0.2% Triton X-100 for 10 minutes and subsequently blocked with 5% bovine serum albumin (BSA) for 1 hour at room temperature. After blocking, cells were incubated with primary antibodies overnight at 4°C and washed three times with PBS. Secondary antibody staining was performed for 2 hours at room temperature, followed by three PBS washes. Imaging was conducted using a Nikon confocal microscope equipped with a 60× oil-immersion objective lens (NA 1.40). The primary antibodies used in immunofluorescence experiments are as follows: CD63 Antibody (MX-49.129.5) (Santa Cruz, sc-5275, 1:100), LAMP1/CD107a Rabbit mAb (Abclonal, A21194, 1:200) and CBP/KAT3A/CREBBP Antibody (C-1) (Santa Cruz, sc-7300, 1:200). The secondary antibodies used in immunofluorescence experiments are as follows: Goat anti-Mouse IgG (H+L) Cross-Adsorbed Secondary Antibody, Alexa Fluor™ 488 (Invitrogen, A-11001, 1:500), Goat anti-Rabbit IgG (H+L) Highly Cross-Adsorbed Secondary Antibody, Alexa Fluor™ Plus 647 (Invitrogen, A-32733, 1:500). Fluorescence intensity was quantified using NIS-Elements AR 4.40 software, and colocalization was assessed by calculating the Pearson’s correlation coefficient, also employing NIS-Elements AR 4.40 software.

#### TIRF microscopy for monitoring MVB mobility

The mobility of exosomes in live cells was assessed using total internal reflection fluorescence (TIRF) microscopy (Nikon). To evaluate the effects of mutations in Munc13-4 on MVB mobility, control and Munc13-4 knockout SUM159 cells were seeded onto glass-bottom dishes. These cells were subsequently co-transfected with NPY-td-Orange2-CD63 and pEGFP-N3-Munc13-4 WT or Munc13-4 mutants. Similarly, to examine the effects of Rab27a mutations on MVB mobility, control or Rab27a knockout SUM159 cells were plated and co-transfected with NPY-td-Orange2-CD63 and pEGFP-C1-Rab27a WT or Rab27a mutants. After 24–48 hours post-transfection, live-cell imaging was performed using a Nikon Ti inverted TIRF microscopy system equipped with a 100× oil-immersion objective (NA 1.49) and an EMCCD camera (Andor DU897). Orange fluorescence was excited using a 532 nm laser with an exposure time of 300 ms. The diffusion coefficient (*D*), representing MVB mobility, was quantified using the Python-based trackpy library (https://soft-matter.github.io/trackpy/dev/tutorial/walkthrough.html).

#### Protein expression and purification

Protein expression of human Munc13-4 and its mutants was performed using the Bac-to-Bac™ baculovirus expression system (Invitrogen). Briefly, recombinant bacmid DNA extracted from DH10Bac was transfected into *Spodoptera frugiperda* clone 9 (Sf9) cells using X-tremeGENE™ 9 DNA Transfection Reagent (Roche) to produce P1 baculovirus. Sequential infection of Sf9 cells with P1 baculovirus generated P2 and P3 baculoviruses. For protein expression, Sf9 cells were infected with P3 baculovirus and cultured for 48 hours. Harvested cells were resuspended in lysis buffer (20 mM Tris-HCl, pH 8.1, 150 mM NaCl) supplemented with protease inhibitors (2 μg/ml aprotinin, 1 μg/ml leupeptin, 1 μg/ml pepstatin, and 1 mM PMSF). Cells were lysed using an AH-1500 Nano Homogenizer (ATS Engineering Inc.) at 800 bar under 4 °C. The lysate was clarified by centrifugation at 16,000 rpm using a JA-25.50 rotor (Beckman Coulter) at 4 °C. Supernatants were incubated with nickel-nitrilotriacetic acid (Ni-NTA) agarose (Qiagen) for 1 hour at 4 °C, followed by two washes with wash buffer (20 mM Tris-HCl, pH 8.1, 150 mM NaCl, 20 mM imidazole). Bound proteins were eluted using buffer containing 20 mM Tris-HCl, pH 8.1, 150 mM NaCl, and 300 mM imidazole. Eluted proteins were further purified by size-exclusion chromatography using a Superdex™ 200 10/300 GL column (Cytiva).

Other proteins were expressed in *Escherichia coli* BL21 (DE3). Protein expression was induced with isopropyl β-D-1-thiogalactopyranoside (IPTG). Harvested cells were lysed as described above. For GST-tagged proteins, clarified lysates were incubated with glutathione Sepharose 4B (GE Healthcare), washed, and eluted using buffer containing 20 mM glutathione (neoFroxx, 1392GR025), 20 mM Tris-HCl, pH 8.1, and 150 mM NaCl. His-tagged proteins were purified as described for Munc13-4. Purified proteins were further processed using ion exchange and size-exclusion chromatography. For transmembrane proteins, 1.5% sodium deoxycholate was included throughout the purification process. Proteins were used immediately after affinity purification.

For cryo-electron microscopy (cryo-EM) complex preparation, Munc13-4 and Rab27a were mixed at a molar ratio of 1:2 in the presence of 1 mM GppNHp (Aladdin, G276465) and incubated at room temperature for 1 hour. The mixture was subjected to size-exclusion chromatography to isolate the stable protein complex.

#### GST pull-down assay

For all GST pull-down assays, 2 μM of the GST-tagged protein was incubated with 3 μM of the target protein at room temperature for 2 hours. Subsequently, the protein mixture was combined with glutathione Sepharose 4B resin (GE Healthcare) and incubated at 4 °C for 1 hour. The resin was then washed four times with wash buffer (20 mM Tris-HCl, pH 8.1, 150 mM NaCl, 0.02% Triton X- 100) to remove unbound proteins. Bound proteins were eluted using an elution buffer containing 20 mM Tris-HCl (pH 8.1), 150 mM NaCl, 0.02% Triton X-100, and 50 mM glutathione. Eluted proteins were analyzed by SDS-PAGE.

#### SNARE assembly assay

For SNARE assembly assay, VAMP-7 SNARE motif (A131C) was labeled with the Förster resonance energy transfer (FRET)-donor dye BODIPY FL (Molecular Probes, B10250) and SN-23- 6CS (S161C) was labeled with the FRET-acceptor dye 5-tetramethylrhodamine (5-TAMRA) (Molecular Probes, T6027). During the experiment, both the donor protein and acceptor protein were at a concentration of 0.5 µM. The concentration of Syx-4 ΔTM (residues 1-275) was 2 µM, and Munc13-4 was at 10 µM. The experiments were performed using a PTI QM-40 spectrophotometer, with an excitation wavelength of 485 nm and an emission wavelength of 513/580 nm, at room temperature. SNARE complex formation signals were interpreted as the FRET proximity ratio (*E_PR_*) between the donor (BODIPY) and acceptor (5-TAMRA). The *E_PR_* was determined using the following equation:

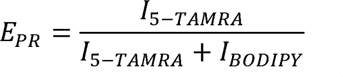

where *I_5-TAMRA_*and *I_BODIPY_* represent the fluorescence intensities of 5-TAMRA and BODIPY FL, respectively, measured under the 485/10 excitation filter.

#### Liposome fusion assay

All lipids were dissolved at an initial concentration of 10 mg/ml in chloroform, except for PI(4,5)PC, which was dissolved in a chloroform: methanol: water mixture (20:9:1) at 1 mg/ml. For Syx-4- liposome preparation, 52% 1-palmitoyl-2-oleoyl-glycero-3-phosphocholine (POPC; Avanti Polar Lipids, 850457), 20% 1-palmitoyl-2-oleoyl-sn-glycero-3-phosphoethanolamine (POPE; Avanti Polar Lipids, 850757), 15% 1,2-dioleoyl-sn-glycero-3-phospho-L-serine (DOPS; Avanti Polar Lipids, 840035), 10% cholesterol (Avanti Polar Lipids, 700000), 1% PI(4,5)PC (Avanti Polar Lipids, 840046), and 2% DiD (Molecular Probes, D307) were mixed to a final lipid concentration of 1 mM. For VAMP-7-liposomes, 38% POPC, 11% POPE, 7% 1,2-dioleoyl-sn-glycero-3-phospho-(1’-myo-inositol) (PI; Avanti Polar Lipids, 850149), 30% cholesterol, 15% sphingomyelin (Avanti Polar Lipids, 860584), and 3% DiI (Molecular Probes, D282) were mixed to the same final concentration. The lipid mixtures were vacuum-dried and resuspended in 2% sodium deoxycholate. Full-length Syx-4 protein (5 µM) and VAMP-7 SNARE-TM protein (5 µM) were incorporated into corresponding liposomes respectively, by incubation at room temperature for 20 minutes. Detergent removal was performed using PD-10 desalting columns (GE Healthcare). The resulting liposomes were combined in a 1:1 ratio with a 60% (w/v) OptiPrep™ stock solution (Serumwerk Bernburg AG), layered with 2 ml of 20% (w/v) OptiPrep™ solution, and 200 µL of 20 mM Tris-HCl (pH 8.1), 150 mM NaCl. Liposomes were centrifuged at 120,000 × *g* for 5 hours at 18°C using an SW55Ti rotor (Beckman). The top fraction was collected and dialyzed in a Slide-A-Lyzer™ dialysis cassette (Thermo Fisher, 66383) for 12 hours before use. For the fusion assay, 10 µM SN-23 was pre-incubated with Syx-4-liposomes at 37°C for 2 hours, while a negative control group was prepared without SN-23. Equal volumes of Syx-4- and VAMP-7-liposomes were mixed to a total volume of 60 µL, and 0.4 µM Munc13-4 protein was added to the experimental group. Fusion was monitored using a FluoDia T70 fluorescence plate reader (Photon Technology Incorporated) at 37°C, with excitation at 530 nm and emission at 580 nm and 667 nm. Liposome fusion signals were quantified by calculating the FRET proximity ratio (*E_PR_*) between the donor (DiI) and acceptor (DiD). The *E_PR_* was determined using the following equation:

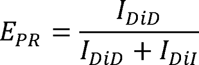

where *I_DiD_*and *I_DiI_* represent the fluorescence intensities of DiD and DiI, respectively, measured under the 530/10 excitation filter.

#### Co-flotation experiment of PD-L1-containing liposomes with Munc13-4

For the preparation of PD-L1-containing liposomes, 80% 1-palmitoyl-2-oleoyl-glycero-3- phosphocholine (POPC; Avanti Polar Lipids, 850457) and 20% 1,2-dioleoyl-sn-glycero-3-phospho-L-serine (DOPS; Avanti Polar Lipids, 840035) were combined to achieve a total lipid concentration of 1 mM. Lipid mixtures were vacuum-dried and resuspended in 2% sodium deoxycholate. PD-L1 protein (5 µM) was incorporated into the liposomes, followed by detergent removal using PD-10 desalting columns (GE Healthcare). The resulting liposome suspension (200 µL) was mixed in a 1:1 ratio with an 80% (w/v) Histodenz™ stock solution (Thermo Fisher, D2158). The mixture was layered sequentially with 350 µL of 30% (w/v) Histodenz™ and 20 µL of 20 mM Tris-HCl (pH 8.1), 150 mM NaCl. The prepared gradient was centrifuged at 240,000 × *g* for 1.5 hours at 18°C using an SW55Ti rotor (Beckman). Fractions (20 µL) were sequentially collected from the top three layers of the gradient, with an additional 20 µL sample retrieved from the bottom layer. These fractions were analyzed by Western blot to assess the co-flotation of PD-L1-containing liposomes with Munc13-4.

#### Structure prediction by AlphaFold

The protein sequences of human PD-L1 (residues 253–290) and Munc13-4 (residues 1049–1080) were obtained from the UniProt database (https://www.uniprot.org/). Three-dimensional structure predictions were performed using AlphaFold v2.3.1, implemented through the Colab-based AlphaFold notebook (https://colab.research.google.com/github/deepmind/alphafold/blob/main/notebooks/AlphaFold.ipyn b).

#### Cryo-EM sample preparation

The Munc13-4–Rab27a complex, bound to GppNHp, was prepared at a concentration of 0.5 mg/ml for cryo-EM analysis. Samples (3.5 µL) were applied to glow-discharged cryo-EM grids (Quantifoil, Cu, R1.2/R1.3, 300 mesh) in an environment of 100% humidity at 4°C. Grids were blotted for 2 seconds with a blotting force of 4 and subsequently vitrified by plunging into liquid ethane using a Vitrobot Mark IV (Thermo Fisher Scientific). Prepared grids were either screened immediately or stored in liquid nitrogen for future use.

#### Cryo-EM data acquisition and image processing

The Munc13-4–Rab27a complex with GppNHp datasets were collected by 300 kV Titan Krios electron microscope (Thermo Fisher Scientific) equipped with a Falcon4 direct electron detector coupled with a SelectrisX energy filter (10 eV slit width). The automated collection was performed using the EPU software in electron event representation (EER) mode, and all micrographs were recorded at a nominal magnification of 165,000 × with a raw pixel size of 0.73 Å on the image plane. The micrographs were recorded in a -0.8 μm to -2.4 μm defocus range, with an electron dose rate of 11.47 e^−^ /Å^2^ /s and a total dose of 50 e^−^ /Å^2^. All the EER movies were pre-processed by CryoSPARC (version 4.5.3)^68^ to perform the motion correction and CTF estimation. 1,424,230 particles were selected by Blob Picking and subsequently subjected to three rounds of 2D classification, and 317,449 particles from the selected classes were subjected to ab-initial 3D reconstruction. The initial volume was further refined by heterogenous refinement and non-uniform refinement to yield a consensus map with 3.36 Å global resolution. To acquire a map with improved characteristics of the C2A and C2B domains, a total of 102,381 particles with apparent features were selected from the 3D classification, yielding a reconstruction at 3.42 Å resolution. Particle subtraction was subsequently applied to these particles, focusing on the regions surrounding the C2A and C2B domains, respectively. Local refinement of the resulting datasets produced focused maps with resolutions of 4.39 Å and 7.2 Å. These two local volume maps were then combined during model building, producing a composite map with a resolution of 4.38 Å. All reported resolutions were estimated using the gold-standard Fourier shell correction 0.143 criterion ^69^. Data collection and refinement statistics are summarized in **Table S1**.

#### Model Building and Refinement of the Munc13-4–Rab27a Complex

Initial models of Munc13-4 (AF-Q70J99-F1) and Rab27a (AF-P51159-F1) were generated using AlphaFold 2 (https://colab.research.google.com/github/deepmind/alphafold/blob/main/notebooks/AlphaFold.ipyn b)^70^. The predicted structures were fitted into the cryo-EM density map through rigid-body docking using UCSF ChimeraX (v1.7.1)^71^. Further manual adjustments were performed using COOT (v0.9.6)^72^. Subsequent real-space refinement of the models was carried out through multiple iterative rounds using PHENIX (v1.20)^73^, followed by final model validation. Figures were prepared using PyMOL (https://pymol.org/2/) and UCSF ChimeraX. Data validation statistics are summarized in **Table S1**.

#### Quantification and statistical analysis

Statistical analyses were conducted using GraphPad Prism software (version 9.3). Specific statistical tests employed for each experiment are detailed in the corresponding figure legends. For comparisons between two groups, either a two-tailed unpaired or paired t-test was applied, as appropriate. For multiple group comparisons, one-way analysis of variance (ANOVA) followed by Tukey’s multiple comparisons test, multiple t-tests, or two-way ANOVA with Sidak’s multiple comparisons test was utilized. A p-value of less than 0.05 (P < 0.05) was considered indicative of statistical significance.

## Supplementary information

Supplementary Figure S1–S12 and Table S1.

## Supplemental Figures

**Figure S1.**
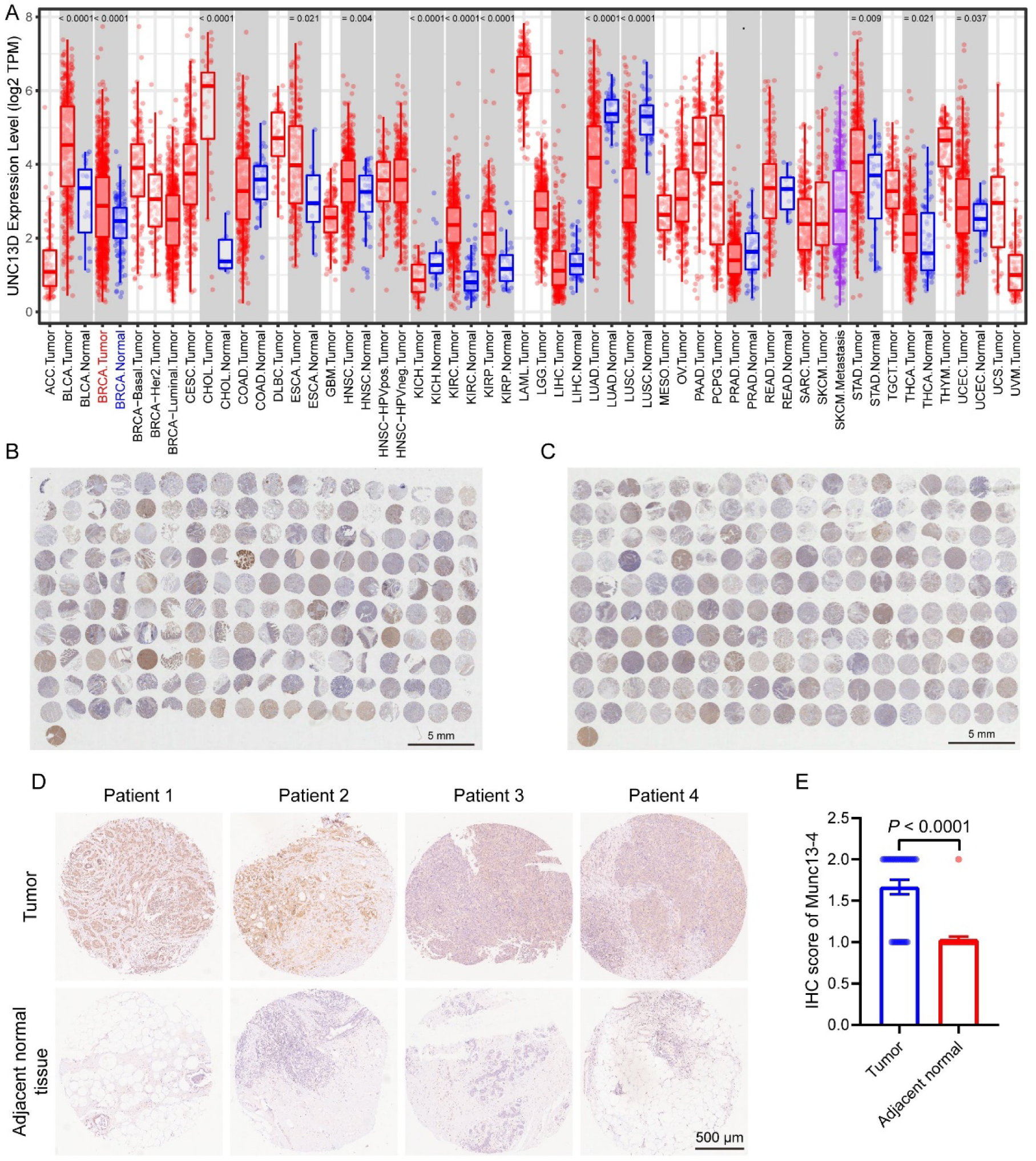
Munc13-4 expression is upregulated in multiple types of tumor tissues, related to Figure 1 **(A)** Differential expression of *UNC13D* (encoding Munc13-4) between tumor and adjacent normal tissues across all TCGA tumors. **(B)** Gross image of immunohistochemical staining for Munc13-4 in a multi-organ carcinoma tissue array comprising 180 tissue cores from 91 patients across 14 tumor types. Scale bar, 5 mm. **(C)** Gross image of immunohistochemical staining for Munc13-4 in a triple negative breast cancer (TNBC) tissue array consisting of 180 tissue cores from 91 patients across 14 tumor types. Scale bar, 5 mm. **(D)** Representative Munc13-4 IHC staining images of independent tissue cores within the TNBC tissue array. Scale bar, 500 μm. **(E)** Munc13-4 IHC scores in tumor tissues and paired adjacent normal tissues from the TNBC tissue array (n = 30). Distributions of gene expression levels are displayed using box plot (A), data are means ± SEM (E), *p-* values were calculated by Wilcoxon test (A and E).

**Figure S2.**
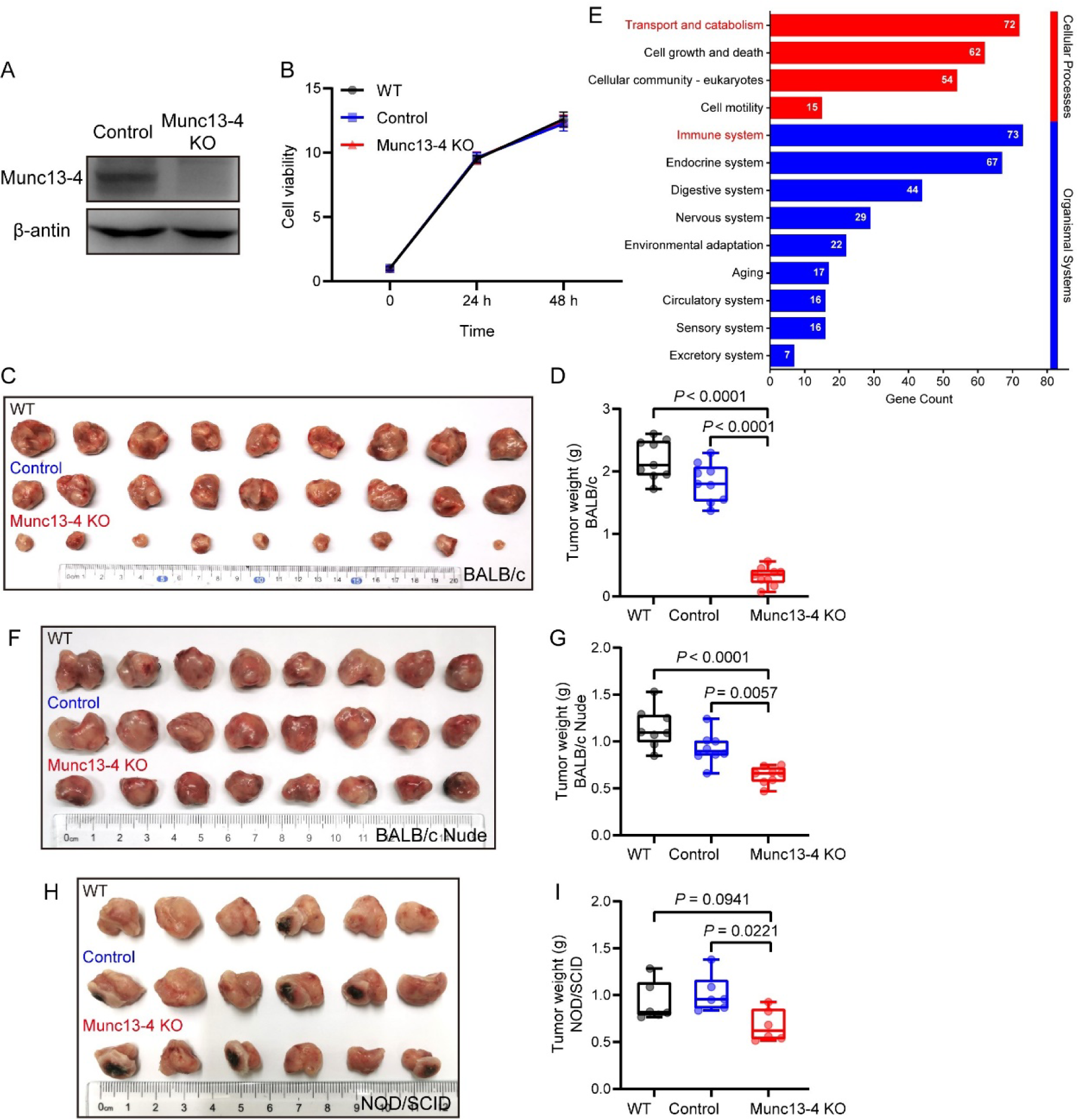
Munc13-4 deficiency in tumor cells suppresses tumor growth through an immune-dependent mechanism, related to Figure 1 **(A)** Validation of Munc13-4 knockout in 4T1 cells by western blot (n = 3). **(B)** Fold change in cell viability (n =6 from duplicate tests). **(C)** Photograph of tumors dissected from BALB/c mice in WT (top), control (middle) and Munc13-4 KO (bottom) groups. **(D)** Tumor weight for the groups shown in (C) (n = 9). **(E)** Enriched KEGG pathways for differentially expressed proteins between control and Munc13-4 knockout 4T1 cells. **(F)** Photograph of tumors dissected from BALB/Nude mice in WT (top), control (middle) and Munc13-4 KO (bottom) groups. **(G)** Tumor weight for the groups shown in (F) (n = 8). **(H)** Photograph of tumors dissected from NOD/SCID mice in WT (top), control (middle) and Munc13-4 KO (bottom) groups. **(I)** Tumor weight for the groups shown in (H) (n = 6). Data are presented as means ± SEM (B). Box plots show all data points, *p*-values were calculated by one-way ANOVA with multiple comparisons (D, G and I).

**Figure S3.**
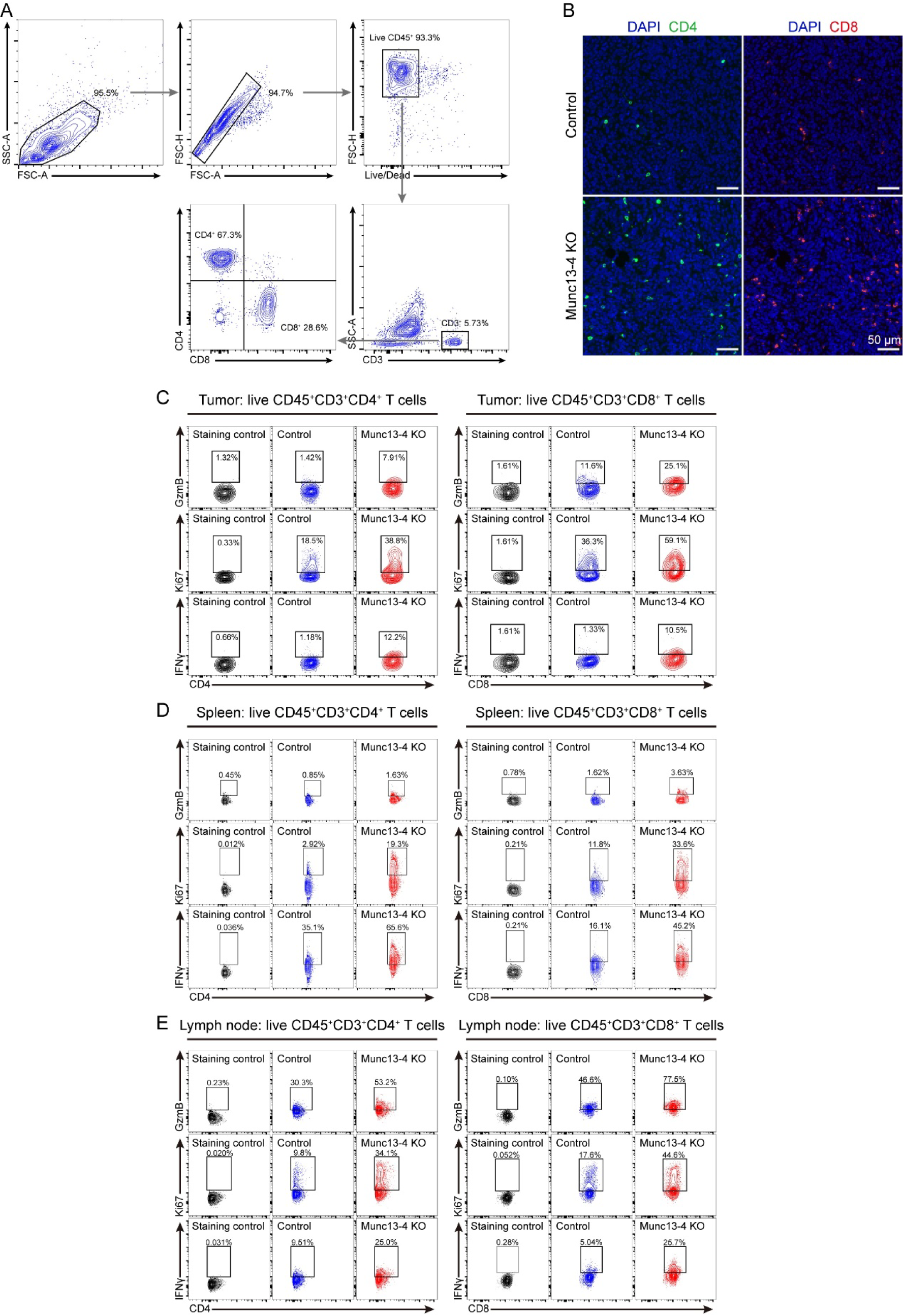
Immune profiling of tumor, spleen and draining lymph node and gating strategy, **related to** Figure 2 **(A)** Representative contour plots showing the general gating strategy for flow cytometric analysis of CD45^+^CD3^+^CD4^+^ and CD45^+^CD3^+^CD8^+^ T cells. **(B)** Representative images of immunofluorescence staining for CD4 and CD8 on tumor tissue sections from orthotopic mouse models of breast cancer established with control or Munc13-4 KO 4T1 cells. Scale bar, 50 μm. **(C–E)** Representative contour plots depicting CD45^+^CD3^+^CD4^+^ (left) and CD45^+^CD3^+^CD8^+^ (right) T cell populations within tumors **(C)**, spleens **(D)**, and draining lymph nodes **(E)** from orthotopic mouse models of breast cancer established with control or Munc13-4 KO 4T1 cells, showing the expression of granzyme B, Ki67 and IFNγ.

**Figure S4.**
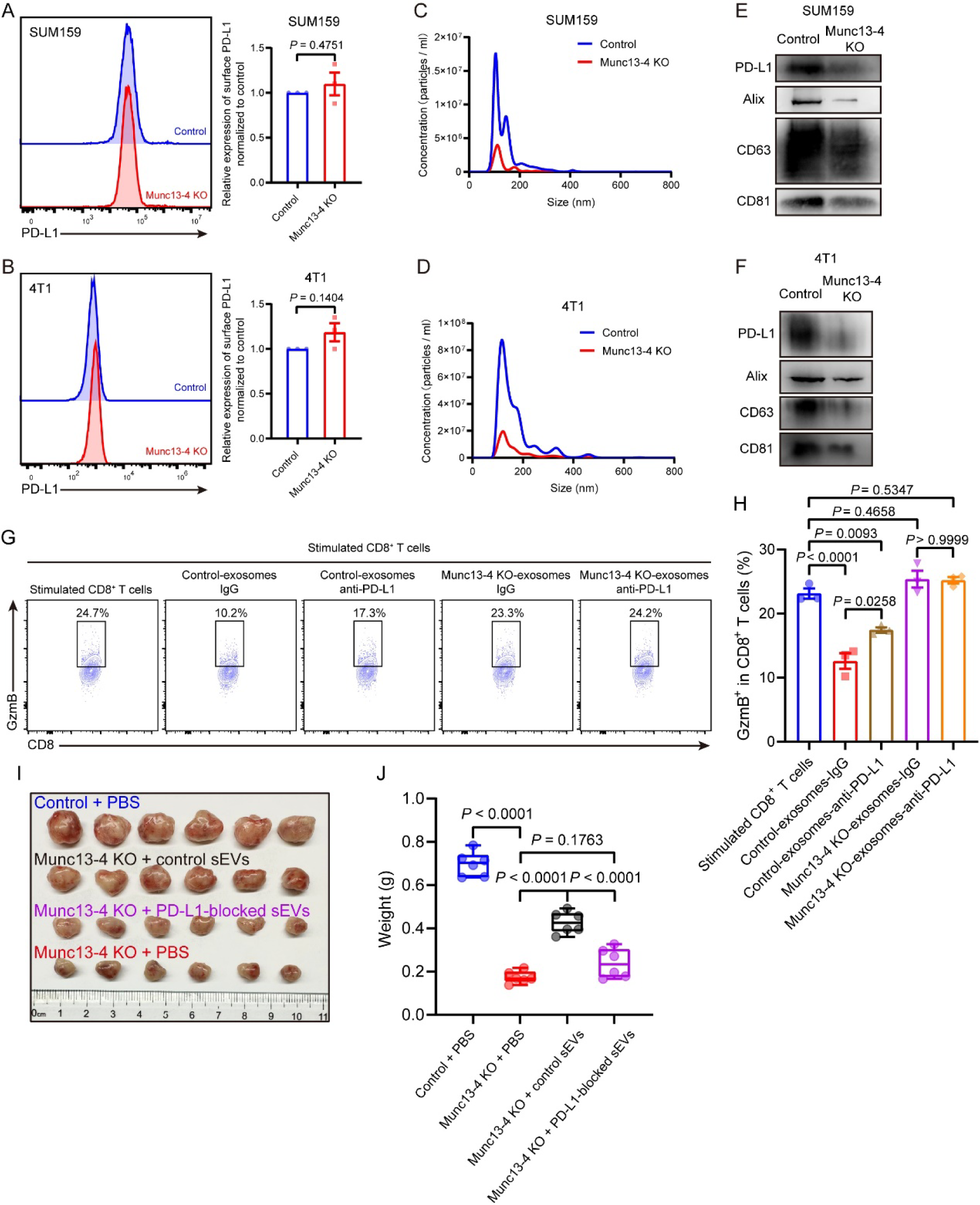
Deletion of Munc13-4 in tumor cells inhibits PD-L1 secretion, related to Figure 3 **(A)** Representative histogram (left) and quantification (right) of the fluorescence intensity of PD-L1 on SUM159 cell surface (n = 3). **(B)** Representative histogram (left) and quantification (right) of the fluorescence intensity of PD-L1 on 4T1 cell surface (n = 3). **(C and D)** Representative NTA traces of EVs secreted by equal numbers of control and Munc13-4 KO SUM159 **(C)** or 4T1 **(D)** cells. **(E and F)** Western bolt analysis of PD-L1, Alix, CD63 and CD81 in EVs secreted by equal numbers of control and and Munc13-4 KO SUM159 **(E)** or 4T1 **(F)** cells (n = 3). **(G and H)** Detection of the effects of exosomal PD-L1 on the cytotoxicity of CD8^+^ T cells. **(G)** Representative contour plots showing the expression of granzyme B within CD8^+^ T cells with indicated exosome treatment. **(H)** Quantification of the percentage of GzmB^+^ cells among CD8^+^ T cells (n = 3). **(I)** Photograph of tumors dissected from orthotopic mouse models of breast cancer with indicated treatment. **(J)** Tumor weight for the groups shown in (I) (n = 6). Data are presented as means ± SEM (A, B and H), box plot shows all data points (J), *p*-values were calculated by unpaired t test (A and B), and one-way ANOVA with multiple comparisons (H and J).

**Figure S5.**
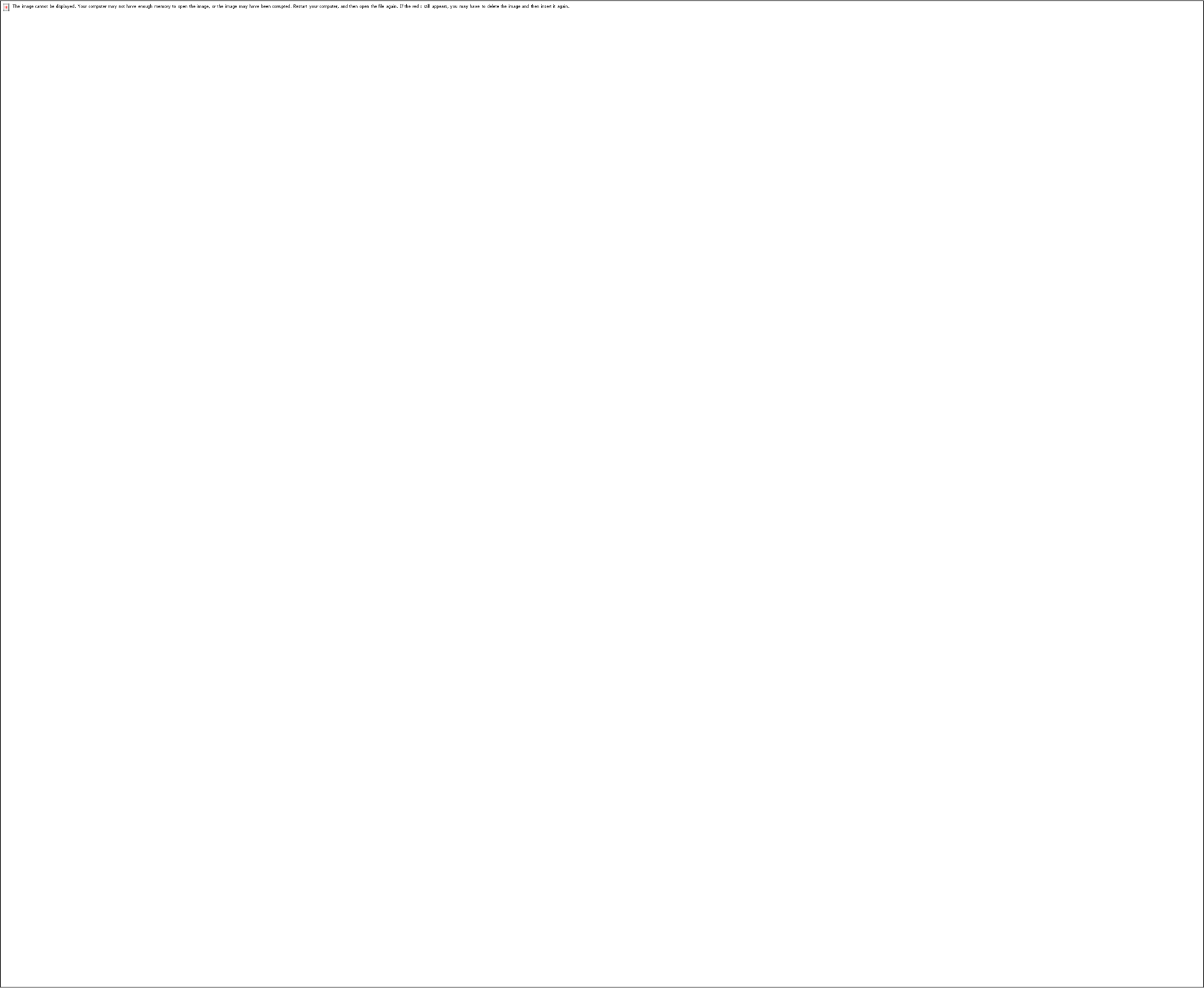
Munc13-4 deficiency in tumor cells boosts immune checkpoint blockade therapy effectiveness, related to Figure 3 **(A)** Schematic of experimental design. **(B)** Tumor growth curves following mammary gland inoculation of control or Munc13-4 KO 4T1 cells, with subsequent treatment of IgG, αPD-1 or αPD-L1 (n = 5). **(C)** Photograph of tumors dissected from BALB/c mice with indicated treatment. **(D)** Tumor weight for the groups shown in (C) (n = 5). Data are presented as means ± SEM (B and D), *p*-values were calculated by one-way ANOVA with multiple comparisons (B) and Multiple t tests (D).

**Figure S6.**
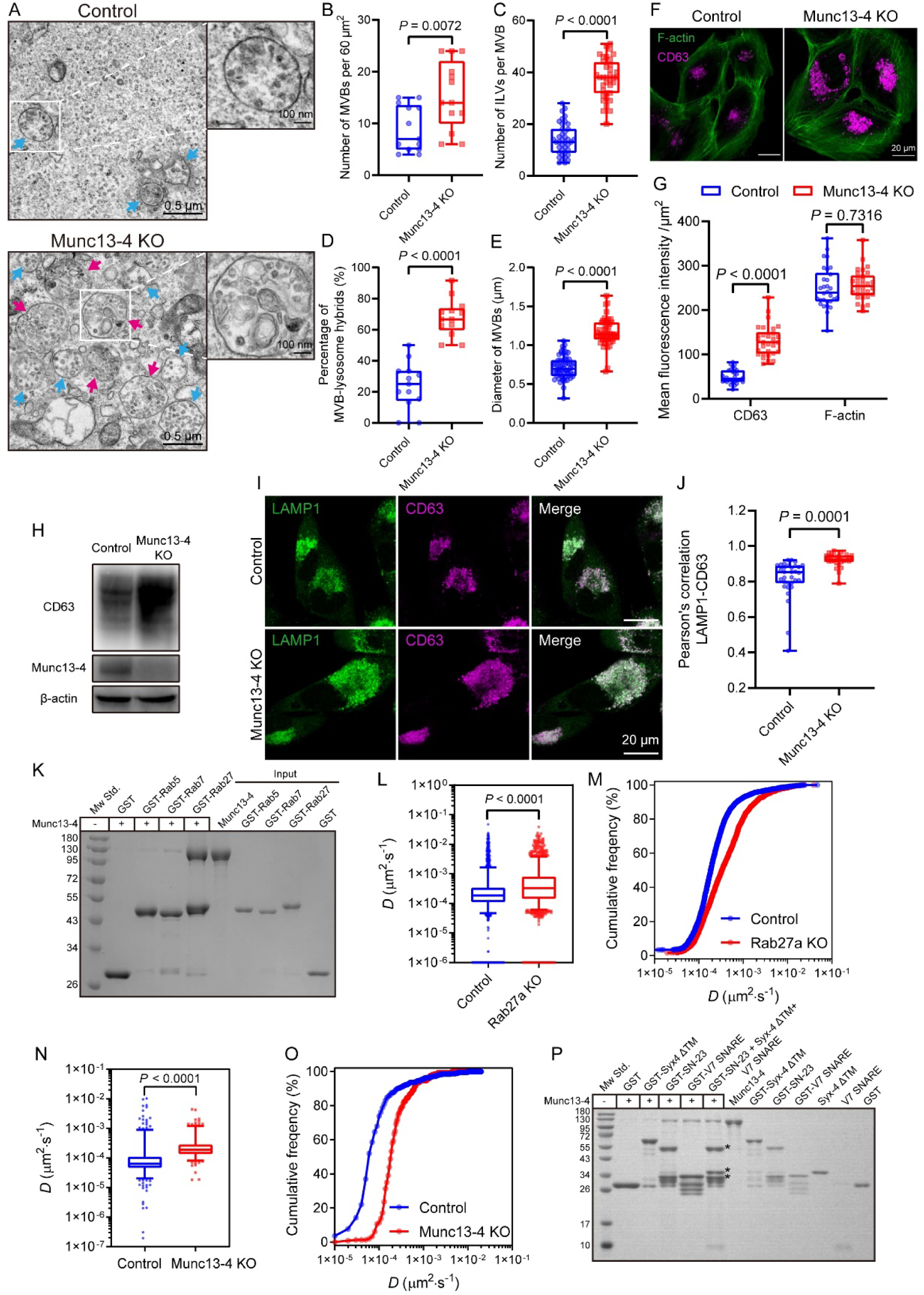
Munc13-4 does not influence MVB biogenesis, related to Figure 4 **(A)** Representative TEM images of MVBs in control and Munc13-4 KO SUM159 cells. Blue arrows indicate pure MVBs, while rosy arrows indicate MVB-lysosome hybrids. Scale bar, 0.5 μm in wide filed and 100 nm in enlarged view. **(B)** Quantification of MVB numbers in control and Munc13-4 KO SUM159 cells (n = 13). **(C)** Quantification of ILV numbers within MVBs in control and Munc13-4 KO SUM159 cells (n = 39 for control group and 36 for Munc13-4 KO group). **(D)** Quantification of the percentage of MVB-lysosome hybrids among total MVBs in control and Munc13-4 KO SUM159 cells (n = 13). **(E)** Quantification of the diameter of MVBs in control and Munc13-4 KO SUM159 cells (n = 58 for control group and 52 for Munc13-4 KO group). **(F)** Representative confocal microscopy images of control and Munc13-4 KO SUM159 cells. Cells were stained with FITC-conjugated phalloidin to label F-actin and immunolabeled with a mouse anti-CD63 primary antibody, followed by Alexa Fluor™ 647-conjugated goat anti-mouse secondary antibody. Scale bar, 20 μm. **(G)** Quantification of the mean fluorescence intensity per μm^2^ of CD63 and F-actin in control and Munc13-4 KO SUM159 cells (n = 30 from triplicate tests.) **(H)** Western blot analysis of total CD63 levels in control and Munc13-4 KO SUM159 cells (n = 3). **(I)** Representative confocal microscopy images of control and Munc13-4 KO SUM159 cells co-immunostained for LAMP1 and CD63. **(J)** Quantification of Pearson’s correlation between LAMP1 and CD63 in control and Munc13-4 KO SUM159 cells (n = 32 for control group and 52 for Munc13-4 KO group, from triplicate tests). **(K)** GST pull-down analysis of the interactions between Munc13-4 and different Rabs (n = 3). **(L)** Quantification of the mean diffusion coefficient (*D*), an index of MVB mobility, in control and Rab27a KO SUM159 cells (n = 2236 for control group and 1854 for Rab27a KO group, from triplicate tests). **(M)** Cumulative frequency distribution of diffusion coefficient (*D*) from (L). **(N)** Quantification of the mean diffusion coefficient (*D*) of MVBs in control and Munc13-4 KO SUM159 cells (n = 387 for control group and 246 for Munc13-4 KO group, from triplicate tests). **(O)** Cumulative frequency distribution of diffusion coefficient (*D*) from (N). **(P)** GST pull-down analysis of the interactions between Munc13-4 and different SNAREs, asterisks indicate that GST-SN-23, Syx-4 (residues 1-274, the cytoplasmic fragment) and VAMP-7 SNARE motif assemble into a 1:1:1 complex (n = 3). Box plots show all data points (B, C, D, E, G and J), or show the 5–95% percentile range of all data, with outliers represented as individual dots (L and M), *p*-values were calculated by unpaired t test (B, C, D, E and J), two-way ANOVA (G) and Mann-Whitney U test (L and N).

**Figure S7.**
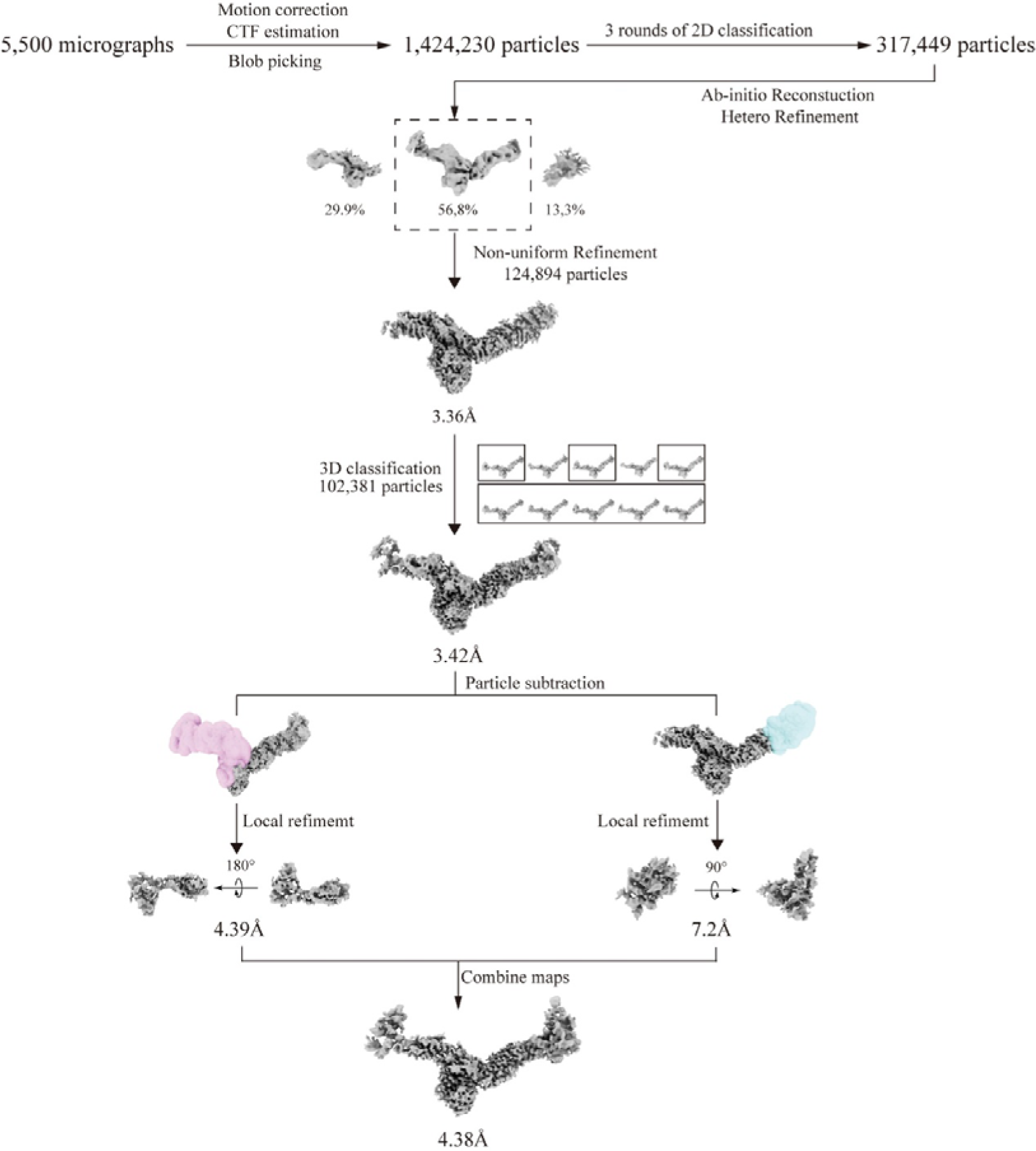
Cryo-EM data processing of the Munc13-4–Rab27a complex, related to. Figure 4 Flowchart illustrating the cryo-EM data processing pipeline using cryoSPARC. The final map was obtained following non-uniform refinement, with a composite map generated through focused refinement on individual regions of interest.

**Figure S8.**
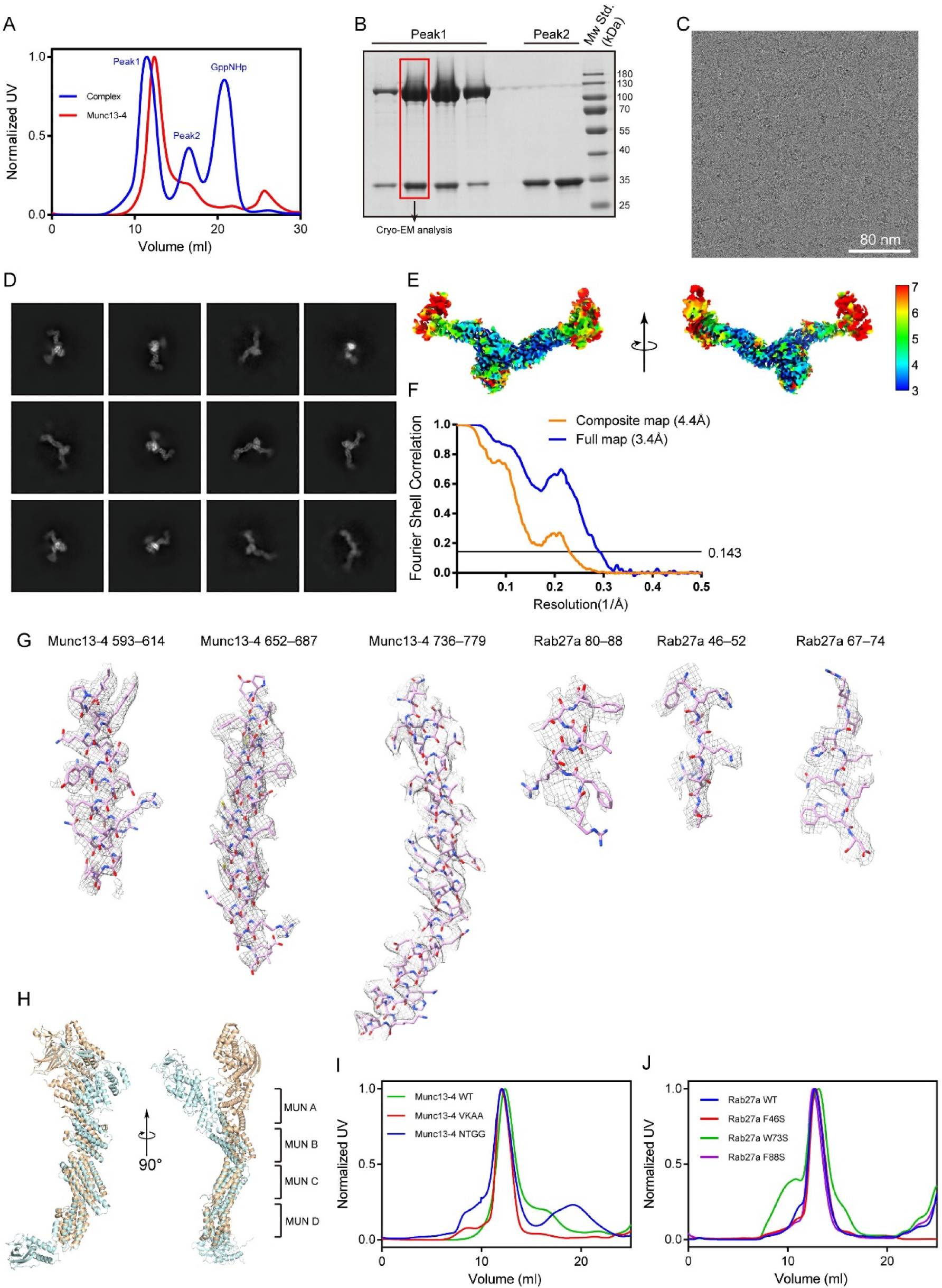
Cryo-EM data analysis of Munc13-4–Rab27a complex, related to Figure 4 **(A and B)** Gel filtration analysis of the Munc13-4–Rab27a complex in the presence of GppNHp **(A)**, followed by SDS-PAGE analysis of the fractions obtained after gel filtration **(B)**. **(C)** Representative 300 kV cryo-EM micrograph of the Munc13-4–Rab27a complex. Scale bar, 80 nm. **(D)** Representative 2D classifications of Munc13-4–Rab27a complex particles obtained from cryo- EM data. **(E)** Cryo-EM density map of the Munc13-4–Rab27a complex, colored according to the estimated local resolution. **(F)** Gold-standard Fourier shell correlation (FSC) curves calculated in cryoSPARC for the full and composite maps. The resolution is determined at FSC = 0.143. **(G)** Representative cryo-EM density map highlighting the helices and sheets at the interface of the Munc13-4–Rab27a complex. **(H)** Comparison of the C1-C2B-MUN domains of Munc13-1 (orange, PDB: 5UE8) and Munc13-4 (cyan). **(I)** Gel filtration analysis of Munc13-4 and its mutants. **(J)** Gel filtration analysis of GST-Rab27a and its mutants.

**Figure S9.**
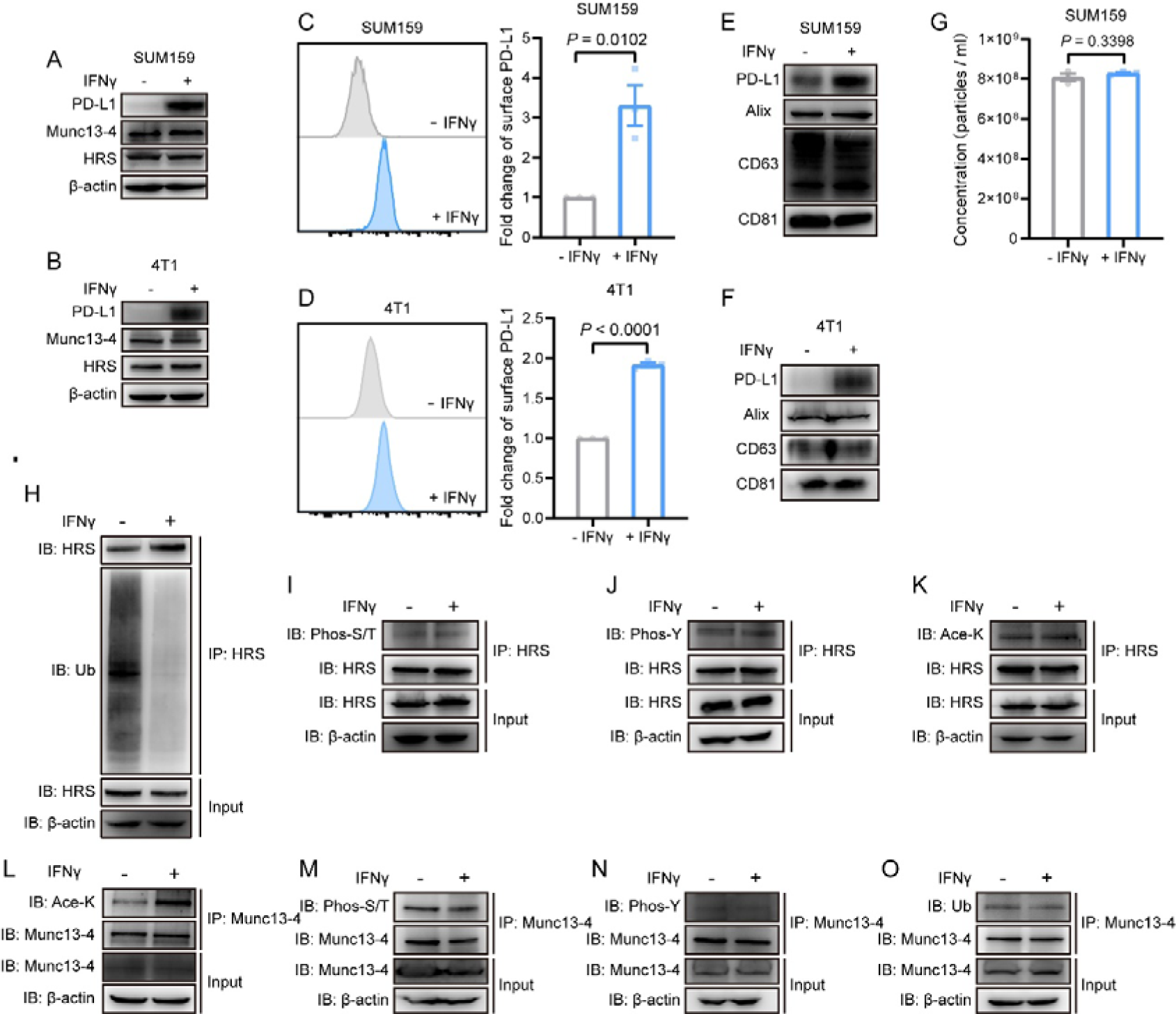
**IFN**γ **induces Munc13-4 acetylation and HRS deubiquitylation, related to** Figure 6 **(A and B)** Western blot analysis of the overall amount of PD-L1, Munc13-4 and HRS in SUM159 cells **(A)** and 4T1 cells **(B)** with or without IFNγ treatment (n = 3). **(C)** Representative histogram (left) and quantification (right) of the fluorescence intensity of PD-L1 on SUM159 cell surface with or without IFNγ treatment (n = 3). **(D)** Representative histogram (left) and quantification (right) of the fluorescence intensity of PD-L1 on 4T1 cell surface with or without IFNγ treatment (n = 3). **(E and F)** Western blot analysis of PD-L1, Alix, CD63 and CD81 abundance on equal numbers of exosomes secreted by equivalent SUM159 cells **(E)** or 4T1 cells **(F)** with or without IFNγ treatment (n = 3). **(G)** Quantification of exosomes secreted by equal numbers of SUM159 cells with or without IFNγ treatment (n = 3). **(H–K)** IP and IB analysis in SUM159 cells to investigate the effect of IFNγ treatment on HRS ubiquitylation **(H)**, serine/threonine phosphorylation **(I)**, tyrosine phosphorylation **(J)** and acetylation **(K)** (n = 3). **(L–O)** IP and IB analysis in SUM159 cells to investigate the effect of IFNγ treatment on Munc13-4 acetylation **(L)**, serine/threonine phosphorylation **(M)**, tyrosine phosphorylation **(N)** and ubiquitylation **(O)** (n = 3). Data are represented as means ± SEM (C, D and G), *p*-values were calculated by unpaired t test (C, D and G).

**Figure S10.**
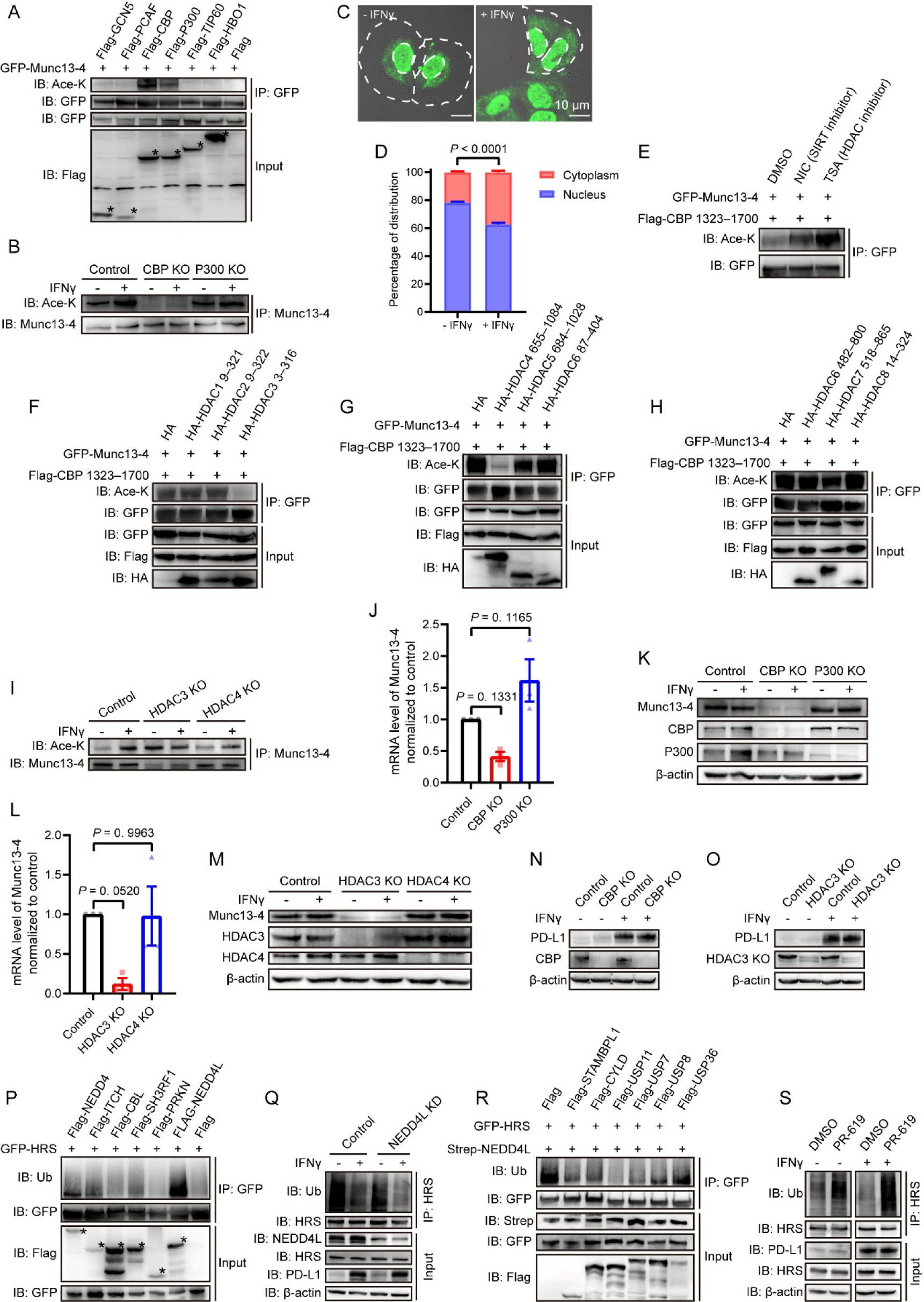
Screen of enzymes regulating Munc13-4 acetylation and HRS ubiquitylation, related to Figure 6 **(A)** IP and IB analysis of Munc13-4 acetylation in HEK293T cells transfected with indicated constructs to search the potential acetyltransferases of Munc13-4 (n = 3). Asterisks denote the blot bands corresponding to the indicated Flag-tagged proteins. **(B)** IP and IB analysis of Munc13-4 acetylation in control, CBP KO and P300 KO SUM159 cells with or without IFNγ treatment to identify the physiological acetyltransferase of Munc13-4 in SUM159 cells (n = 3). **(C and D)** Examination of the effect of IFNγ stimulation on the subcellular distribution of CBP in SUM159 cells. Representative confocal microscopy images of SUM159 cells immunostained for CBP, with or without IFNγ treatment **(C)**, and quantification of the percentage of CBP localized to the nucleus versus the cytoplasm **(D)** (n = 118 for the - IFNγ group and 88 for the + IFNγ group, from triplicate experiments). The white dotted line delineates the outline of the cell and nucleus. Scale bar, 10 µm. **(E)** IP and IB analysis of Munc13-4 acetylation in HEK293T cells transfected with indicated constructs and treated with DMSO, NIC or TSA to identify the type of deacetylase targeting Munc13-4 (n = 3). **(F–H)** IP and IB analysis of Munc13-4 acetylation in HEK293T cells transfected with indicated constructs to identify the potential deacetylases of Munc13-4. Screen of HDAC1–3 **(F)**, HDAC4–6 **(G)** and HDAC6–8 **(H)** (n = 3). **(I)** IP and IB analysis of Munc13-4 acetylation in control, HDAC3 KO and HDAC4 KO SUM159 cells with or without IFNγ treatment to identify the physiological deacetylases of Munc13-4 in SUM159 cells (n = 3). **(J)** Quantification of the mRNA level of Munc13-4 in control, CBP KO and P300 KO SUM159 cells (n = 3). **(K)** Western blot analysis of the total amount of Munc13-4 in control, CBP KO and P300 KO SUM159 cells with or without IFNγ treatment (n = 3). **(L)** Quantification of the mRNA level of Munc13-4 in control, HDAC3 KO and HDAC4 KO SUM159 cells (n = 3). **(M)** Western blot analysis of the total amount of Munc13-4 in control, HDAC3 KO and HDAC4 KO SUM159 cells with or without IFNγ treatment (n = 3). **(N)** Western blot analysis of the total amount of PD-L1 in control and CBP KO SUM159 cells with or without IFNγ treatment (n = 3). **(O)** Western blot analysis of the total amount of Munc13-4 in control and HDAC3 KO SUM159 cells with or without IFNγ treatment (n = 3). **(P)** IP and IB analysis of HRS ubiquitylation in HEK293T cells transfected with indicated constructs to search the potential E3 ubiquitin ligase of HRS (n = 3). Asterisks denote the blot bands corresponding to the indicated Flag-tagged proteins. **(Q)** IP and IB analysis of Munc13-4 acetylation in control and NEDD4L knockdown (KD) SUM159 cells with or without IFNγ treatment to examine the role of NEDD4L in the regulation of HRS ubiquitylation (n = 3). **(R)** IP and IB analysis of HRS ubiquitylation in HEK293T cells transfected with indicated constructs to search the potential deubiquitinases of HRS (n = 3). Asterisks denote the blot bands corresponding to the indicated Flag-tagged proteins. **(S)** IP and IB analysis of HRS ubiquitylation in SUM159 cells with or without PR-619 treatment in the absence and presence of IFNγ (n = 3). Data are represented as means ± SEM (D, J and L), *p*-values were calculated by two-way ANOVA (D) and one-way ANOVA with multiple comparisons (J and L).

**Figure S11.**
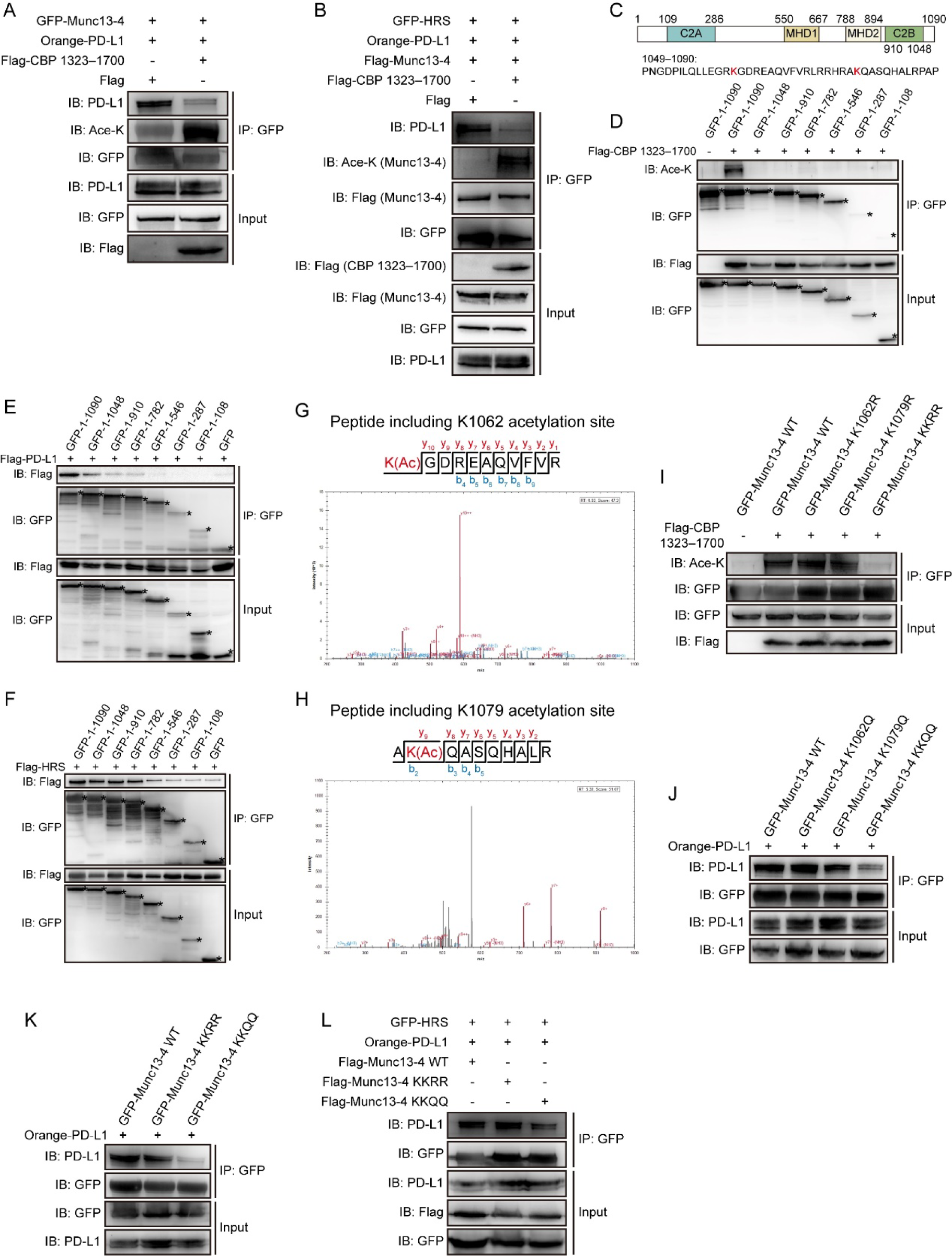
The acetylation of Munc13-4 disrupts its interaction with PD-L1, related to Figure 6 **(A)** Co-IP and IB analysis in HEK293T cells transfected with indicated constructs to investigate the effect of Munc13-4 acetylation on its interaction with PD-L1 (n = 3). **(B)** Co-IP and IB analysis in HEK293T cells transfected with indicated constructs to investigate the effect of Munc13-4 acetylation on the interaction of HRS with PD-L1 as well as Munc13-4 (n = 3). **(C)** Diagram of domains of Munc13-4 and the sequence of the Munc13-4 1049–1090 motif. **(D)** IP and IB analysis of the acetylation of Munc13-4 truncations in HEK293T cells transfected with indicated constructs to search the acetylation sites within Munc13-4 (n = 3). **(E and F)** Co-IP and IB analysis in HEK293T cells transfected with indicated constructs to identify the motif in Munc13-4 responsible for interacting with PD-L1 **(E)** and HRS **(F)** (n = 3). **(G and H)** Acetylated peptides of Munc13-4 containing either K1062 **(G)** or K1079 **(H)** acetylation sites were identified by LC-MS/MS. Panels (G and H) show the peptide sequences, with the LC-MS/MS-detected fragments labeled above the sequence for C-terminal y ions and below the sequence for N-terminal b ions. **(I)** IP and IB analysis of the acetylation of Munc13-4 mutants in HEK293T cells transfected with indicated constructs to identify the acetylation sites within Munc13-4 (n = 3). **(J)** Co-IP and IB analysis in HEK293T cells transfected with indicated constructs to determine the effects of acetylation-mimicking mutations at Munc13-4 acetylation sites on the Munc13-4–PD-L1 interaction (n = 3). **(K)** Co-IP and IB analysis in HEK293T cells transfected with indicated constructs to determine the effects of mutations mimicking or disrupting Munc13-4 acetylation on the Munc13-4–PD-L1 interaction (n = 3). **(L)** Co-IP and IB analysis in HEK293T cells transfected with indicated constructs to determine the effects of mutations mimicking or disrupting Munc13-4 acetylation on the interaction of HRS with PD-L1 (n = 3).

**Figure S12.**
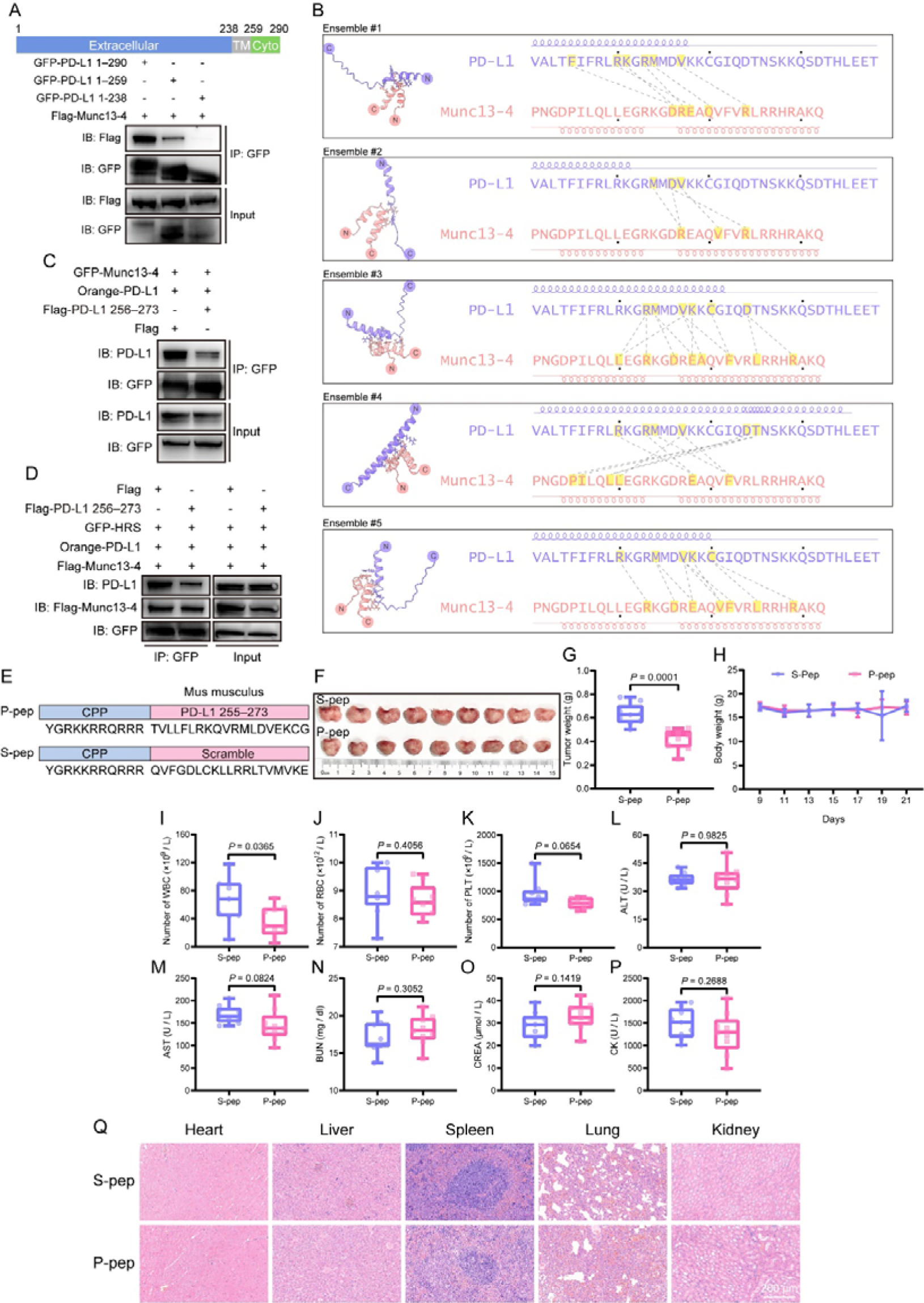
A peptide disrupting PD-L1**–**Munc13-4 interaction elicits no toxicity, related to Figure 7 **(A)** Diagram of domains of PD-L1 (upper panel) and co-IP and IB analysis in HEK293T cells transfected with indicated constructs to investigate the motif in PD-L1 required for binding to Munc13-4 (lower panel). **(B)** Predicted residue contacts between PD-L1 (residues 253–290) and Munc13-4 (residues 1049–1080). Five ensembles generated by AlphaFold multimer are displayed on the left, where N- and C-terminus of the proteins are indicated. Residue-to-residue contacts are manually curated and shown by dashed lines. Residues that show potential contacts are shaded in yellow. The secondary structure elements for each ensemble are indicated alongside the sequences. **(C)** Co-IP and IB analysis in HEK293T cells transfected with indicated constructs to investigate the effect of PD-L1 256–273 motif on the interaction between Munc13-4 and PD-L1. **(D)** Co-IP and IB analysis in HEK293T cells transfected with indicated constructs to assess the effect of PD-L1 256–273 motif on the interaction of HRS with PD-L1 and Munc13-4. **(E)** Diagram of the sequences for P-pep and S-pep used *in vivo*. P-pep comprises a cell-penetrating peptide (CPP) fused to the mouse PD-L1 255–273 motif, whereas S-pep consists of a CPP linked to a scrambled sequence containing the same amino acid composition as the mouse PD-L1 255–273 motif. **(F)** Photograph of tumors dissected from mice treated with S-pep (top) or P-pep (bottom). **(G)** Tumor weight for the groups shown in (F) (n = 9). **(H)** Body weight curves of mice treated with S-pep or P-pep over the experimental period (n = 9). **(I–K)** Blood routine examinations of mice treated with S-pep or P-pep. Quantification of white blood cells (WBC, **I**), red blood cells (RBC, **J**), and platelets (PLT, **K**) (n = 9). **(L–P)** Blood biochemical analysis of mice treated with S-pep or P-pep. Quantification includes alanine transaminase (ALT, **L**), aspartate transaminase (AST, **M**), blood urea nitrogen (BUN, **N**), creatinine (CREA, **O**), and creatine kinase (CK, **P**) levels (n = 9). **(Q)** Representative histopathological examination of dissected heart, liver, spleen, lung, and kidney tissues stained with hematoxylin and eosin (H&E) (n = 3). Scale bar, 200 μm. Data are represented as means ± SEM (H), box plots show all data points (G and I–P), *p*-values were all calculated by unpaired t test.

**Table S1.** Cryo-EM data collection, refinement, and validation statistics, related to STAR Methods.

| Methods |  |  |
| --- | --- | --- |
|  | Munc13-4-Rab27a complex<br>Consensus map (EMDB-63239) | Munc13-4-Rab27a complex (EMDB-62922) (PDB 9LA9) |
| **Data collection and processing** |  |  |
| Magnification | 165,000 | 165,000 |
| Voltage (kV) | 300 | 300 |
| Electron exposure (e-/Å<sup>2</sup>) | 50 | 50 |
| Defocus range (µm) | -2.4 to -0.8 | -2.4 to -0.8 |
| Pixel size (Å) | 0.73 | 0.73 |
| Symmetry imposed | C1 | C1 |
| Initial particle images (no.) | 1,424,230 | 1,424,230 |
| Final particle images (no.) | 102,381 | 102,381 |
| Map resolution (Å) | 3.42 | 4.38 |
| FSC threshold | 0.143 | 0.143 |
| Map resolution range (Å) | 2.8 to 4.7 | 3.2 to 8 |
| **Refinement** |  |  |
| Initial model used (PDB code) |  | AlphaFold2 |
| Model resolution (Å) |  | 3.4 |
| FSC threshold |  | 0.143 |
| Model resolution range (Å) |  |  |
| Map sharpening *B* factor (Å<sup>2</sup>) |  | -100.07 |
| Model composition |  |  |
| Non-hydrogen atoms |  | 9285 |
| Protein residues |  | 1158 |
| Ligands |  | GNP: 1 |
| *B* factors (Å<sup>2</sup>) |  |  |
| Protein |  | 103.98 |
| Ligand |  | 109.921 |
| R.m.s. deviations |  |  |
| Bond lengths (Å) |  | 0.011 |
| Bond angles (°) |  | 1.012 |
| Validation |  |  |
| MolProbity score |  | 1.95 |
| Clashscore |  | 12.18 |
| Poor rotamers (%) |  | 0.59 |
| Ramachandran plot |  |  |
| Favored (%) |  | 94.95 |
| Allowed (%) |  | 4.80 |
| Disallowed (%) |  | 0.26 |

